# Evolutionary transfer learning enables organism-wide inference of mammalian enhancer landscapes

**DOI:** 10.64898/2026.04.07.717039

**Authors:** Chengxiang Qiu, Riza M. Daza, Ian C. Welsh, Rupali P. Patwardhan, Beth K. Martin, Tony Li, Shizhao Yang, Camiel C.A. Mannens, Seppe De Winter, Niklas Kempynck, Megan L. Taylor, Olivia Fulton, Truc-Mai Le, Diana R. O’Day, Jean-Benoît Lalanne, Silvia Domcke, Stephen A. Murray, Stein Aerts, Cole Trapnell, Jay Shendure

## Abstract

Understanding and modeling how a single human genome concurrently encodes gene regulatory programs for thousands of cell types remains a central challenge in genomics and machine learning. Most human cell types emerge during embryonic, fetal, and pediatric development which are inaccessible to comprehensive molecular profiling. To circumvent this, we hypothesized that the mismatch in evolutionary rates between *cis-*acting enhancers that modulate gene expression (fast) and the *trans*-acting regulatory factors that specify cell types (slow) creates an opportunity for ‘evolutionary transfer learning’. Specifically, models trained to predict cell type-specific enhancers in one species should generalize to the orthologous cell types and enhancers of related species. To test this, we generated a single-cell atlas of chromatin accessibility spanning mouse embryonic day 10 (E10) to birth (P0). Using combinatorial indexing^1^, we profiled 3.9 million nuclei from 36 staged embryos, resolving genome-wide accessibility in 36 cell classes and 140 cell types. We then trained a series of multi-output deep learning models (CREsted^2^), each addressing limitations of the preceding approach, towards the goal of genome-wide prediction of distal enhancers across major developmental lineages. An ‘evolution-naive’ model achieved strong performance on heldout peaks, but exhibited two failure modes during genome-wide inference: overprediction at tandem repeats and conflation of promoter and distal enhancer grammars. An ‘evolution-aware’ model resolved these by regrouping accessible regions based on their retention and functional coherence across mammalian evolution, but failed to generalize across species. Finally, an ‘evolution-augmented’ model, STEAM (Synteny-aware Transfer learning for Enhancer Activity Modeling), incorporated enhancer orthologs from 241 mammalian genomes (Zoonomia^3^) in a synteny-supervised manner. This increased the effective data scale by as much as 195-fold, markedly improving generalization across mammals despite greater label noise. We applied STEAM to the genome-wide inference of cell class-specific distal developmental enhancers in humans, mice (*HumMus*) and 239 additional mammals^3^ (*BabaGanoush*), *i.e.* 32 × 241 = 7,712 genome-wide distal enhancer prediction tracks. Together, our results unify advances in single-cell profiling, deep learning, and comparative genomics into a framework for the evolutionary transfer learning of noncoding regulatory grammars. More broadly, our work supports the view that model organisms and evolutionarily diverse genomes are indispensable resources for accelerating and enhancing the AI-enabled exploration of human biology.

**Note:** An interactive version of this preprint, together with count matrices, CREsted models, prediction tracks, code and reproducible figures, is available at [this link](https://doi.org/10.62329/hxkk6249) (ref ^4^).

## INTRODUCTION

Recent breakthroughs in protein structure prediction, exemplified by AlphaFold^5^ and RoseTTAFold^6^, are often credited to the synergy of transformer architectures, large-scale computation, and a growing repository of atomic-resolution structures^7^. But the most indispensable ingredient was arguably evolution. Protein orthologs can diverge dramatically in primary amino acid sequence while maintaining a nearly identical structure and function, and protein orthologs can functionally complement one another across vast evolutionary distances (*e.g.* yeast ↔ human)^8^. This mismatch—sequences change quickly, while structure-function changes slowly—is the supervisory signal that makes high-fidelity structure prediction possible. Modern structure predictors achieved their accuracy only once densely populated multiple sequence alignments (MSAs) from genome/metagenome sequencing, generated in abundance by the genomics community for unrelated goals, allowed them to extract co-evolutionary couplings that no amount of structural data alone could provide^9,10^.

Surprisingly, no analogous principle has risen to the forefront of regulatory genomics. Despite a decade of deep learning in this field^11,12^, nearly all major efforts lack an evolutionary dimension (with some important exceptions^13–20^). Models such as Borzoi, BPnet, Enformer, Basenji and AlphaGenome predict biochemical readouts from DNA sequence with impressive fidelity, yet are trained on a single reference genome and a static set of molecular profiles^21–25^. A parallel class of DNA models (*e.g.* Evo, Nucleotide Transformer, GPN) draws on multi-species corpora during pretraining, internalizing evolutionary statistics implicitly^26–29^, analogous to how ESMFold^30^ learns co-evolution from protein sequence alone, without an MSA at inference. But whether implicit exposure substitutes for principled comparative supervision remains an open question.

We reasoned that an analogous mismatch exists in gene regulation: the sequences of *cis-*regulatory elements (*e.g.* enhancers, promoters) change quickly, but the *trans-*acting programs that act through them (*i.e.* repertoires of cell type-specific transcription factors) change slowly. A striking demonstration of this contrast in evolutionary tempos comes from transgenic mice bearing human chromosome 21^31^. Hepatocytes from these animals almost perfectly reproduce the epigenomic and transcriptomic signatures of this chromosome in human hepatocytes^32–37^. Put another way, the *trans*-acting milieus of mammalian cell types are ancient and stable, while the *cis*-regulatory landscape rewires continuously. In principle, this contrast should provide a rich supervisory signal for modeling regulatory grammars. A proof-of-concept comes from Oh and Beer, who recently exploited it in paired human and mouse datasets to discover thousands of conserved enhancers missed by prior alignment methods^19^.

Why not simply learn human regulatory grammars on human data? First, data availability: nearly all human cell types arise during embryonic, fetal, and pediatric stages that are inaccessible to deep molecular profiling. A comprehensive human epigenomic atlas—from zygote to embryo to fetus to child to adult, at whole-body scale and with high temporal resolution—may never be obtainable. In contrast, we and others have constructed whole-organism single-cell timelapses of zebrafish^38–40^, mouse^41–45^ and rhesus macaque^46^ development. Given that cell type-specific enhancer grammars are far more conserved than the genome sequences on which they act, models trained on such comprehensive model organism atlases and then applied to the human genome could provide ‘virtual access’ to *in vivo* human regulatory landscapes that are otherwise off-limits.

A second motivation is statistical. Single-species models are limited not only by the number of regulatory elements in a genome, but also by restricted sequence diversity. Even if the full human enhancer catalog were known, the number active in any given cell type may be insufficient to learn high-dimensional grammars involving combinatorial motif syntax, positional preferences, and long-range dependencies. Orthologous enhancers provide a natural solution: sequence variation under shared functional constraint. Because purifying selection on enhancers is weak at the nucleotide level, their orthologs frequently encode highly divergent yet functionally equivalent sequences^47,48^. And although excessive divergence can prevent alignment, the initial human-mouse genome comparison showed that syntenic regions—identical by descent, even if neutrally evolving—remain alignable at ∼0.50 substitutions per position^49^, while other studies have shown functional complementation of the positional orthologs of enhancers (here termed ‘syntenic enhancers’) across even greater phylogenetic distances (*e.g.* mammals ↔ chicken or zebrafish^47,48^). In this sense, synteny provides the critical bridge between sequence divergence and functional conservation, allowing evolutionarily related, functionally conserved elements to be identified even when sequence similarity has heavily eroded. Collectively, the genomes of >6,500 extant mammals are likely to contain over 1 billion examples of developmental enhancers, the vast majority of which are likely to have positional orthologs with equivalent specificities elsewhere in the set.

We set out to train a multi-class deep learning model of regulatory grammars that spanned all major cell lineages of mammalian development, specifically isolated distal enhancer grammars, and generalized robustly to genome-wide inference across mammalian species. We first generated a single-cell, whole-animal atlas of chromatin accessibility spanning mouse embryonic day 10 to birth, resolving 36 cell classes and 140 cell types across 3.9 million nuclei. Using CREsted^2^, we trained an evolution-naive sequence model that exposed key failure modes in genome-wide inference, then an evolution-aware successor model that resolved these through positional ortholog retention and coherence. Finally, we developed STEAM (Synteny-aware Transfer learning for Enhancer Activity Modeling), augmenting training with synteny-anchored orthologs from up to 241 mammalian genomes^3^, and applied it to inference of the genome-wide enhancer landscape across all major developmental lineages in human, mouse, and 239 additional mammals.

## RESULTS

### Single-nucleus chromatin accessibility profiling of whole mouse embryos from E10 to P0

For the molecular profiling of the emergence and diversification of mammalian cell types at the whole organism scale, we focused on *Mus musculus* (mouse). In a recent study, we collected and ontogenetically staged 523 mouse embryos spanning E8 to P0^42^. We then applied scRNA-seq to profile 11.4 million nuclei from 74 of these embryos, spaced at 2- to 6-hour intervals. For this study, we selected 36 embryos from this same staged set, spaced at 6-hour intervals from E10 to P0, and extracted nuclei (**Fig. 1a; Supplementary Table 1**; E17.75 bin absent). We preferentially selected female embryos to minimize sex-linked mapping artifacts^50^ (29 female; 7 male) and processed whole animals without tissue dissection, as previously^42–44,51^

**Figure 1.**
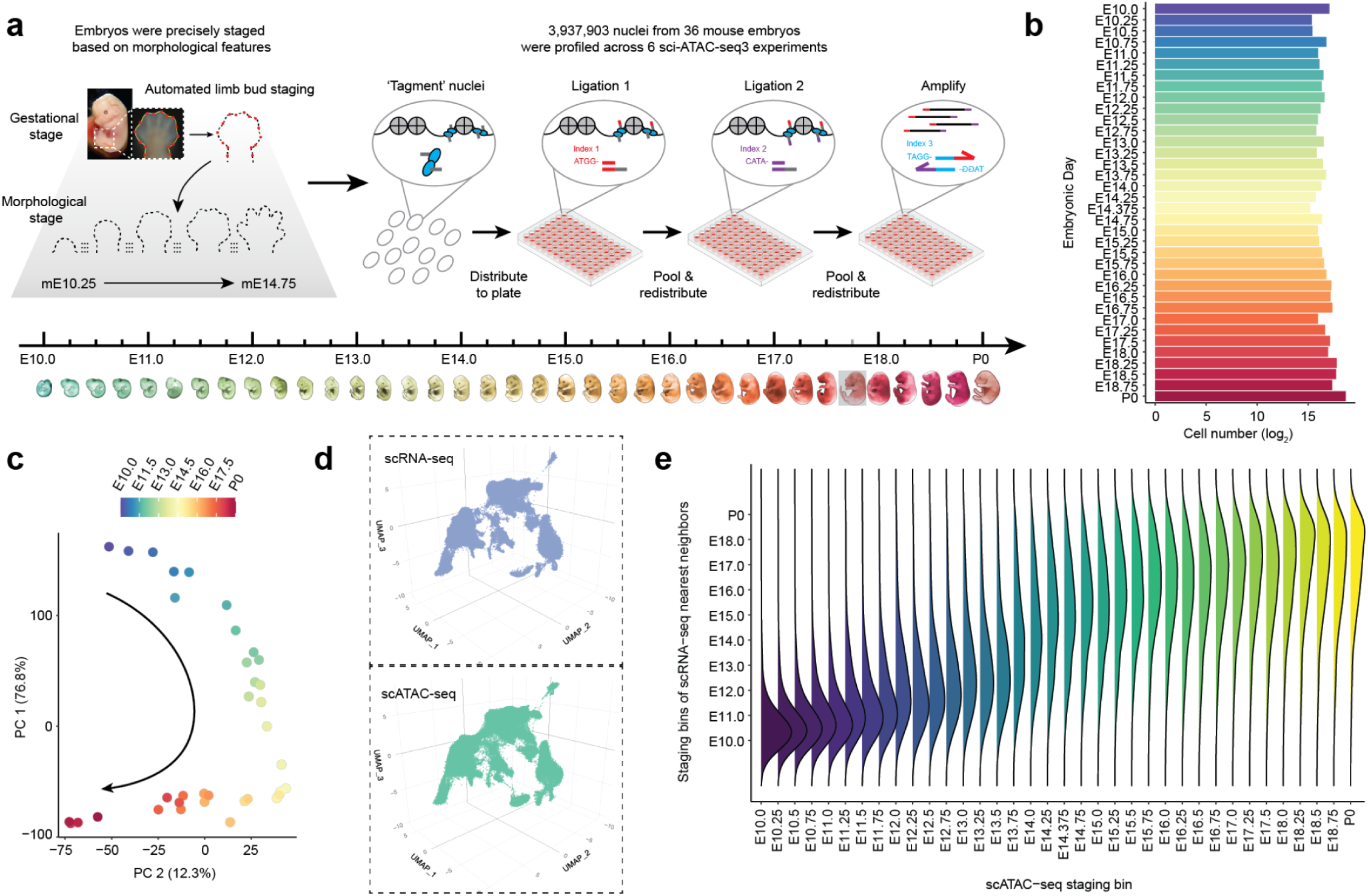
A single-cell chromatin-accessibility timelapse of mouse development from E10 to P0. **a,** Embryos were collected and staged based on morphological features, including an automated method leveraging limb bud geometry for E10–E15^42^. We selected 36 embryos for sci-ATAC-seq3, exactly one per 6-hour bin from E10 to P0 with the exception of E17.75, which remains missing from the series (gray box). Across six experiments, we profiled chromatin accessibility in 3.9 million cells from 36 embryos. The staging diagram and sci-ATAC-seq3 workflow are adapted from Domcke et al.^53^ and Qiu et al.^42^. **b,** The log_2_ scaled number of cells profiled per embryo, sorted by staging bin. **c,** Embeddings of pseudo-bulk ATAC-seq profiles of 36 embryos in PCA space, plotting PC1 vs. PC2. Briefly, single cell profiles were downsampled to a uniform number per embryo, aggregated to 36 pseudo-bulk profiles, and subjected to dimensionality reduction (PCA). Embryos are colored by staging bin; the arrow highlights developmental progression. **d,** 3D UMAP visualization of co-embedded cells from scRNA-seq^42^ and scATAC-seq (this study) data. From each dataset, we selected 15,000 cells from each of 36 shared staging bins, and then performed integration with scGLUE^52^. The same UMAP is shown twice, highlighting cells from either scRNA-seq (top) or scATAC-seq (bottom). **e,** For scATAC-seq cells from each staging bin (*x-*axis), the density of staging-bins-of-origin of their nearest scRNA-seq derived neighbors (n = 10) in the scGLUE-derived co-embedding is plotted (*y*-axis). The density curves on the *y*-axis, generated by geom_density_ridges, were smoothed using a bandwidth of 0.6. P0 was treated as E19.0 for this analysis.

Six sci-ATAC-seq3 experiments were conducted across defined stage ranges (E10–E13.75 × 3; E14–E15.75; E16–E18; E18.25–P0). Following sequencing, demultiplexing, embryo assignment, mapping to the mm10 reference genome, and duplicate removal (avg. 24%; **Supplementary Table 2**), peaks were called per embryo and merged into a master set of 376,574 peaks. After aggressive filtering of low-quality nuclei and doublets (**Supplementary Fig. 1a**), the final matrix comprised 3,937,903 cells × 376,574 peaks, averaging 109,386 cells per embryo (median 90,804; range 37,287–420,085; **Fig. 1b**). Median unique reads per cell was 3,922 (35.7% overlapping peaks; 15.0% within 1 kb of a transcription start site (TSS)). Of note, we observed modest shifts in fragment size distributions across timepoints and mild experiment-level batch structure across pseudobulk profiles (**Supplementary Figs. 1b, 2a**); while these may reflect a mixture of biological and technical factors, we did not detect any substantial influence on clustering, trajectory structure, or model behavior.

**Figure 2.**
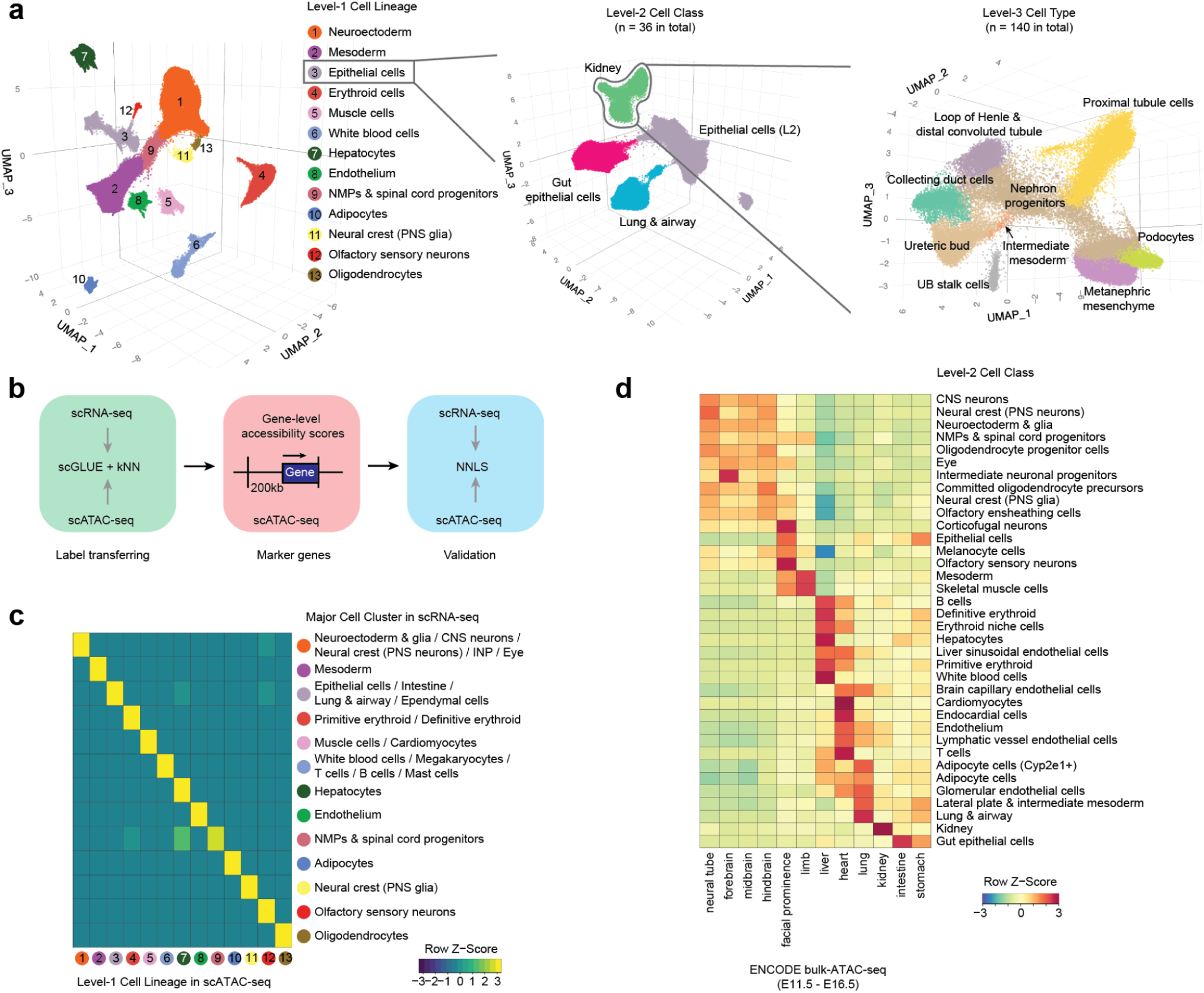
Iterative annotation of a single cell mouse developmental chromatin accessibility atlas. **a,** Left: 3D UMAP visualization of 3,937,903 single cell chromatin accessibility profiles, colored by Level-1 annotation. Middle & Right: Three rounds of annotation were performed through dimensionality reduction, clustering and label transfer, illustrated by epithelial cells (Level-1) → kidney (Level-2) → various renal epithelial cell types (Level-3). **b,** To annotate cell lineages, classes or types, we applied a *k*-NN heuristic to an scGLUE^52^-derived co-embedding with scRNA-seq data from matching timepoints (Fig. 1d). Next, we manually refined these annotations by verifying that marker genes for each cell lineage exhibited high gene-level accessibility scores, calculated by counting reads overlapping gene bodies (plus upstream 200 kb). Finally, we validated these annotations by performing non-negative least squares (NNLS) regression between gene-level accessibility scores from scATAC-seq and gene expression levels from scRNA-seq^44^. **c,** Heatmap of combined regression coefficients (NNLS; row-scaled) for 13 major scRNA-seq cell clusters^42^ (rows) vs. 13 Level-1 scATAC-seq cell lineages (columns; annotation key in panel **a**). CNS: central nervous system. PNS: peripheral nervous system. INP: intermediate neuronal progenitors. NMPs: Neuromesodermal progenitors. **d,** Overlap of 36 peak sets from Level-2 scATAC-seq cell classes (rows) vs. 12 peak sets from bulk ATAC-seq of dissected mouse embryonic tissues by ENCODE^65^. Briefly, peaks identified by ENCODE for each tissue were merged across timepoints (E11.5–E16.5). We also downsampled each Level-2 cell class to 10,000 cells, recalled peaks, calculated their specificity scores using Jensen-Shannon divergence, and selected the 2,000 most specific peaks. For each Level-2 cell class peak set × ENCODE tissue peak set, we counted the number of base pairs overlapping base pairs. Finally, we performed row-based normalization to account for total length, and plotted the resulting Z-scores.

Upon performing principal components analysis (PCA), pseudobulked ATAC profiles showed top PCs strongly correlated with developmental time (PC1+PC2 = 89%; **Fig. 1c**) and high correlation between neighboring timepoints (**Supplementary Fig. 2a**). Integration of downsampled scATAC-seq and scRNA-seq matrices (540,000 cells each; 15,000 cells × 36 timepoints) via scGLUE^52^ confirmed that nearest neighbors largely originated from matched or neighboring timepoints (**Figs. 1d,e**), further validating data quality.

### Iterative annotation of 13 cell lineages, 36 cell classes and 140 cell types

We iteratively annotated these 3.9 million scATAC-seq profiles through three rounds of dimensionality reduction, clustering, and label transfer. For the first round, we applied BPCells^54^ to perform latent semantic indexing (LSI)-based dimensionality reduction, embedded cells in 3D UMAP space, and used Louvain clustering to define major cell lineages (**Fig. 2a**). Labels were transferred from our matched scRNA-seq atlas^42^ via a *k*-nearest neighbors (*k*-NN) heuristic applied to the scGLUE co-embedding (**Fig. 1d**), and then manually refined based on marker gene accessibility scores (**Fig. 2b**; **Supplementary Table 3**).

This first round yielded 13 <u>cell lineages</u> (Level-1): neuroectoderm, mesoderm, epithelial cells, erythroid cells, muscle cells, white blood cells, hepatocytes, endothelium, NMPs & spinal cord progenitors, adipocytes, neural crest (PNS glia), olfactory sensory neurons, and oligodendrocytes (**Fig. 2a-c**). Lineage compositions were highly consistent between scATAC-seq and scRNA-seq modalities (**Supplementary Fig. 2b-c**), and Level-1 annotations were robust to the choice of features for dimensionality reduction (**Supplementary Fig. 2d**)^55,56^. Two additional rounds of iterative subclustering, with steps as in the first round, yielded 36 <u>cell classes</u> (Level-2) and 140 <u>cell types</u> (Level-3) (**Fig. 2a**; **Supplementary Figs. 3-5**). We note that these annotations are preliminary and may be refined as we and others further analyze the data.

Of 204 cell types previously annotated in our scRNA-seq atlas^42^, 114 mapped one-to-one to 114 Level-3 annotations, 56 mapped many-to-one to 24 Level-3 annotations (reflecting lower resolution in scATAC-seq data, *e.g.* diencephalon/midbrain/MHB → midbrain), and 34 were absent, either because they predate E10 (*e.g.* notochord) or are too rare to detect (<0.25% of the scRNA-seq dataset, which profiled three times as many cells) (**Supplementary Fig. 5**; **Supplementary Table 4**). Conversely, 138 of 140 Level-3 scATAC-seq cell types had at least one scRNA-seq counterpart. A first exception was ureteric bud stalk cells, present in the scRNA-seq atlas but not resolved there as a distinct cluster (**Supplementary Fig. 6a-c**). The other exception was a population we term ‘erythroid niche cells’, absent from the scRNA-seq atlas entirely (**Supplementary Fig. 6d-e**). These cells cluster with the yolk sac-derived primitive erythroid lineage but are distinguished by a delayed temporal profile persisting through P0 (**Supplementary Fig. 6f**), markedly reduced accessibility at embryonic globin loci (*Hba-x*, *Hbb-y*), and increased accessibility at lysosomal (*Lamp1*, *Ctsd*, *Ctsb*, *Ctsl*, *Hexa*), heme catabolism (*Hmox1*), and erythropoiesis-related cytokine genes (*Epo*, *Csf2ra*) (**Supplementary Fig. 6g**; **Supplementary Table 5**)^57–63^. Their absence from scRNA-seq data could reflect a transcriptionally low-output state. Alternatively, they could correspond to primitive pyrenocytes—the extruded nuclei of enucleating primitive erythrocytes^64^. We favor the former interpretation, as pyrenocytes would be expected to lack active chromatin remodeling machinery, which is difficult to reconcile with the coordinated accessibility gains observed, together with robust quality metrics for this population (**Supplementary Fig. 6h**).

To recover accessible regions specific to rarer cell types that may be absent from the initial master peak set, we re-called and merged peaks from pseudobulked profiles at each annotation level, which yielded 695,910 (495 Mb), 795,998 (583 Mb), and 793,816 (568 Mb; 21% of mm10) peaks at Levels 1, 2, and 3, respectively. As expected, these annotation-aware peak sets overlap substantially, *e.g.* 89% and 77% of Level-3 intervals are also spanned by Level-2 and Level-1 intervals, respectively. Reassuringly, they also overlap extensively with peaks identified by bulk ATAC-seq of mouse embryonic tissues by the ENCODE Consortium^65^ (**Fig. 2d**).

These data, peak sets and annotations constitute a high-resolution, single-cell, whole-organism map of chromatin accessibility, sampled at 6-hour resolution from the onset of organogenesis (E10) through birth (P0) in the mouse, the most widely used mammalian model of human biology, and at 3.9 million nuclei, is also among the largest scATAC-seq datasets released to date. Together with our matched scRNA-seq atlas from equivalently staged mouse embryos^42^, they provide a powerful resource for modeling the emergence and diversification of cell type-specific *cis-*regulatory programs at the scale of the whole organism and throughout prenatal development.

### Articulation of modeling objectives

Training a sequence model to predict chromatin accessibility on held-out peaks drawn from the same distribution is a necessary but insufficient benchmark for our purposes. As outlined in the introduction, our goal is the <u>genome-wide</u> inference of <u>distal enhancer grammars</u> for <u>all major developmental cell types</u> and <u>all</u> <u>available mammalian genomes</u>. This is a harder target, and one for which we can set four quantifiable expectations. <u>First</u>, the top-scoring regions under the model for any given cell type should be enriched for sequences accessible in that cell type. <u>Second</u>, the model should specifically learn distal enhancer grammar, rather than a conflation of enhancer, promoter and/or grammars. <u>Third</u>, the model should not assign high scores to tandem repeats, which are highly represented and widely distributed throughout mammalian genomes, and whose subsequences can resemble transcription factor binding sites. <u>Fourth</u>, inference should generalize across mammalian genomes, *i.e.* orthologous enhancers should exhibit similar and well-calibrated patterns of predicted cell type-specificity.

We articulate these goals at the outset because performance on held-out peaks (goal #1), while tracked throughout, is an incomplete guide to performance on goals #2 through #4. Our path to a model well-suited to all four criteria was iterative, as each successive model (evolution-naive → evolution-aware → STEAM) was developed in response to failure modes revealed by its predecessor. Having arrived at STEAM, we evaluate the primary and intermediate models against a set of metrics that collectively represent our four goals, which aims to clarify what each successive iteration contributes and what limitations remain (**Supplementary Fig. 7**). We refer back to these metrics throughout the manuscript as we work through the model iterations.

### An evolution-naive model performs well on held-out peaks, but not on genome-wide inference

As a first step, we trained a sequence model on the mouse atlas alone, without leveraging any comparative genomic information. For this and throughout, we used CREsted, a framework for decoding enhancers through sequence-to-function deep learning^2^. A key attribute of CREsted is its ability to jointly learn and predict quantitative chromatin accessibility from DNA sequence across many cell types within a single model (**Fig. 3a**).

**Figure 3.**
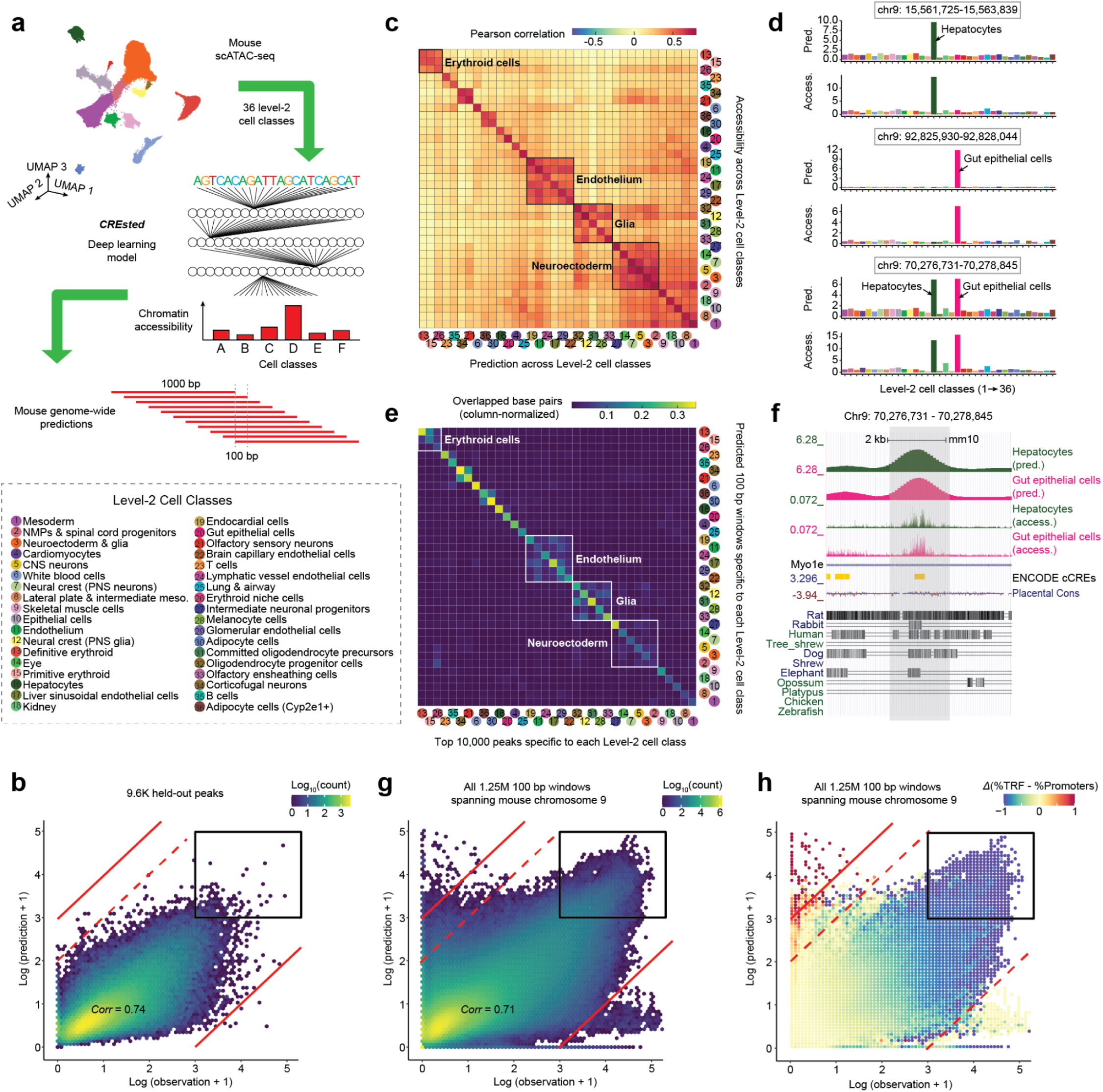
Evolution-naive modeling of chromatin accessibility across 36 Level-2 cell classes. **a,** We trained CREsted^2^, a multi-output convolutional neural network framework that predicts quantitative chromatin accessibility from DNA sequence, on the mouse scATAC-seq timelapse atlas. For evolution-naive modeling, we trained on Level-2 cell class peaks and then sought to make genome-wide predictions. For this, the mouse genome was segmented into sliding 1,000 bp windows, and predictions averaged to 100-bp resolution. Candidate 100-bp windows for each cell class were identified by requiring both a Z-transformed predicted score ≥ 3 and a Z-transformed specificity ≥ 3. **b,** Scatterplots of observed (*x*-axis; normalized Tn5 cut-site counts; log-n scaled) vs. predicted (*y*-axis; evolution-naive CREsted model; log-n scaled) chromatin accessibility across 9,583 peaks held out during fine-tuning, shown for all 36 Level-2 cell classes and colored by log_10_ scaled density. Solid red lines demarcate grossly overpredicted (above *y* > *x* + 3) and grossly underpredicted (below *y* < *x* - 3) windows. A dotted red line marks moderately overpredicted windows (above *y* > *x* + 2). The black rectangle highlights windows that are both predicted and observed to be highly accessible (*x* > 3 && *y* > 3). The same data, stratified by cell class, are shown in **Supplementary Fig. 8**. **c,** Pairwise Pearson correlations were computed among the 36 Level-2 cell classes between normalized Tn5 cut-site counts from scATAC-seq data (rows) and model predictions (columns) for 9,583 held-out peaks. Correlation coefficients are shown in the heatmap, with rows and columns ordered by hierarchical clustering. The black rectangles highlight subsets of related cell classes exhibiting elevated cross-correlation. Cell class labels are shown to the left of panel **e**. **d,** Observed (top) and predicted (bottom) chromatin accessibility is plotted for each of the 36 Level-2 cell classes, for three representative regions from the test set, including two regions exhibiting accessibility in one cell class, and one region exhibiting accessibility in two related cell classes. In all three cases, we observe strong agreement between observed and predicted accessibility. **e,** For each 100-bp bin, we averaged predictions from overlapping windows to obtain a 36-dimensional accessibility vector, and computed cell-class specificity scores using a softmax-based metric (**Supplementary Fig. 9a-b**). Plotted here are pairwise overlaps (bp) between each class’s 10,000 peaks used for initial training (columns) and 100-bp windows with both high predicted accessibility and specificity (rows; both Z-transformed scores ≥ 3; **Methods**), column-normalized by the total size of each class’s training peaks. Cell class labels are shown to the left. **f,** Genome browser view of predicted (top two rows; evolution-naive CREsted model) and observed (normalized Tn5 cut-site counts) chromatin accessibility for the gut epithelial and hepatocyte cell classes, together with ENCODE cCRE, placental conservation and synteny tracks (bottom three rows). This example corresponds to the bottommost example highlighted in panel **d**. Generated using the UCSC Genome Browser (mm10). **g,** Same as panel **b**, but across 1.25M × 100 bp windows spanning mouse chr. 9 for all 36 Level-2 cell classes. **h,** Same as panel **g,** but colored by the difference between the percentage of windows overlapping tandem repeats and the percentage overlapping promoters (± 2.5 kb of annotated TSS). The scale is oriented such that red indicates tandem repeat enrichment and blue indicates promoter enrichment. Gross overpredictions are highly enriched for tandem repeats (top left of plot), while very highly accessible observations/predictions are highly enriched for promoters (black rectangle).

We trained this ‘evolution-naive’ CREsted model to predict chromatin accessibility at candidate regulatory regions across 36 cell classes. After excluding promoter-proximal peaks from the Level-2 peak set (± 2.5 kb of annotated TSS), we trained on 299,281 × 2,114-bp windows (CREsted’s input length^2^) centered on the 10,000 most specific peaks per class, followed by fine-tuning on 106,813 windows centered on the 3,000 most specific peaks per class. For both stages, peaks were split into training (chrs. 1-7, 11-17, 19, X, Y), validation (chrs. 8, 10), and test (chrs. 9, 18) sets.

By the standard benchmark of held-out peak prediction, the resulting model exhibited strong performance on held-out peaks (Pearson’s *r* = 0.74, range = 0.63–0.82 across 36 cell classes; **Fig. 3b**; **Supplementary Fig. 8**). Cross-correlation analysis revealed largely cell-class-specific signals alongside coherent blocks corresponding to closely related lineages (**Fig. 3c**). Inspection of specific loci found that the model accurately predicted both cell class-specific peaks and peaks shared across related lineages (**Fig. 3d**).

Most sequence-based models of regulatory activity are trained and evaluated on predefined sets of candidate enhancers^12,22,24^, as we do above. However, whether such models can be meaningfully applied beyond these curated regions has received comparatively little attention. This omission may be consequential: strong performance within a training distribution does not guarantee that a model has learned regulatory logic that generalizes throughout the rest of the genome, let alone across to the genomes of other species. To assess this, we tiled the mouse reference genome with overlapping 1-kb windows (100-bp step size; predictions averaged across overlapping windows) and applied the ‘evolution-naive’ model, yielding 36 genome-wide tracks of predicted accessibility and specificity for 28M × 100 bp windows spanning mm10 (**Supplementary Fig. 9a-b**).

At a coarse level, we recovered cell class-specific regulatory landscapes. Highly scoring windows were enriched for overlap with distal peaks used during training and recapitulated observed accessibility patterns at representative loci (**Fig. 3e-f**). However, while overall performance remained comparable to held-out peaks (Pearson’s *r* = 0.71 for held-out chr. 9), scatterplots revealed numerous windows that were grossly overpredicted (log-n[predicted/observed] > 3) or underpredicted (log-n[predicted/observed] < -3) (**Fig. 3g**). Gross overpredictions were strongly enriched for tandem repeats (odds ratio (OR): 24-fold), (**Fig. 3h**; **Supplementary Fig. 9c-d**). We speculate this reflects motif-like matches within repeating units that mimic enhancer grammar but without regulatory activity. Such false positives are highly problematic for genome-wide inference, as tandem repeats and other low complexity sequences appear frequently throughout the genome.

A second failure mode relates to promoters, which are among the most highly accessible regions in the genome, yet whose sequences encode little of the regulatory grammar underlying cell type-specific expression. Housekeeping promoters are constitutively accessible across cell types, while cell type-specific promoter accessibility reflects the activity of distal enhancers acting in *trans*, rather than specificity encoded in the promoters themselves. Despite our exclusion and zeroing of peaks within 2.5 kb of annotated TSSs, the most highly accessible observations and predictions were very strongly enriched for promoters (OR: 14-fold for windows with log-n[predicted] > 3 and log-n[observed] > 3; **Fig. 3g-h**). Although technically accurate — these regions are indeed highly accessible — this conflation is problematic, as our broad objective is to learn distal enhancer grammars rather than raw accessibility, and promoter masking was intended to enforce this. We interpret this as evidence that the evolution-naive model learned a mixed grammar spanning both distal enhancers and promoters, possibly because many promoters are unannotated, ambiguously annotated, or otherwise not captured by simple distance-to-TSS exclusion.

To summarize, the evolution-naive model reproduces a familiar pattern in regulatory genomics, in that strong performance on the in-distribution benchmark masks behavior that is unsuitable for the inference task we actually care about. Although the model generalizes well across most of the genome, the highest scoring genome-wide predictions overwhelmingly correspond to non-enhancer signals. For example, among the 0.01% highest scoring windows per cell class on held-out chromosome 9 (n = 4,465), 95% overlapped promoters, despite our the exclusion of promoter-overlapping peaks and zeroing of accessibility within 2.5 kb of annotated TSS, while 65% of the remaining high-scoring windows overlapped tandem repeats (**Supplementary Fig. 7a-c**, columns 2-3).

We emphasize that annotation-guided filtering of enhancer predictions overlapping tandem repeats and/or promoters is not a viable solution. Tandem repeats are too densely distributed; although only 3.4% of mm10 is annotated as tandem repeat sequence, 83% of the genome lies within 2.5 kb of one or more of these 1.1 million elements. Filtering predictions near promoters is more tractable in principle, as only 8% of mm10 falls within 2.5 kb of an annotated TSS, but cell-type-specific enhancers are themselves highly enriched in promoter-proximal regions and would be lost alongside the confounders. Together, these failure modes preclude the robust genome-wide inference of distal enhancer grammars in the mouse genome with the evolution-naive model, let alone other mammalian genomes.

### Evolutionary coherence separates enhancer grammars from promoter grammars

We set out to develop a revised strategy to address the confounding of enhancer and promoter grammars by explicitly incorporating evolutionary signals, as well as to implement a heuristic solution to tandem repeat-induced false positives. For the former, we reasoned that most bona fide enhancers that play an important role in mammalian development should satisfy two criteria. First, they should have orthologs in most mammalian genomes (‘syntenic retention’). Second, such syntenic orthologs should exhibit consistent predicted activity profiles across cell classes when scored by the evolution-naive model, regardless of the extent of nucleotide-level sequence conservation (‘evolutionary coherence’).

To quantify syntenic retention, we began with the 2.79 million 100-bp windows (∼10% of the mouse genome; Z ≥ 3 for both predicted accessibility and specificity in at least one cell class; **[Supplementary File 1](https://shendure-web.gs.washington.edu/content/members/cxqiu/public/backup/jax_atac/download/Supplementary_File_1_100bp_Windows_Specific_to_36_Level2_Cell_Classes.bed.gz)**) and performed liftovers from mm10 to each of 240 Zoonomia genomes^3^ (**Fig. 4a**). The resulting counts were clearly bimodal, and we retained 1.47 million windows (53%) for which syntenic orthologs were identified in at least half of species (**Fig. 4b**). To quantify evolutionary coherence, we extracted 2,114 bp (CREsted input length) from each mammalian genome for which an ortholog was identified (296 million sequences; mean 201 orthologs per window). We then scored these orthologs with the evolution-naive model and compared the mouse ortholog’s 36-value predicted accessibility vector against each non-mouse ortholog’s 36-value vector. The resulting correlations were again clearly bimodal, and we retained 547,317 windows (37%) with a median Pearson’s *r* ≥ 0.6 (**Fig. 4c**).

**Figure 4.**
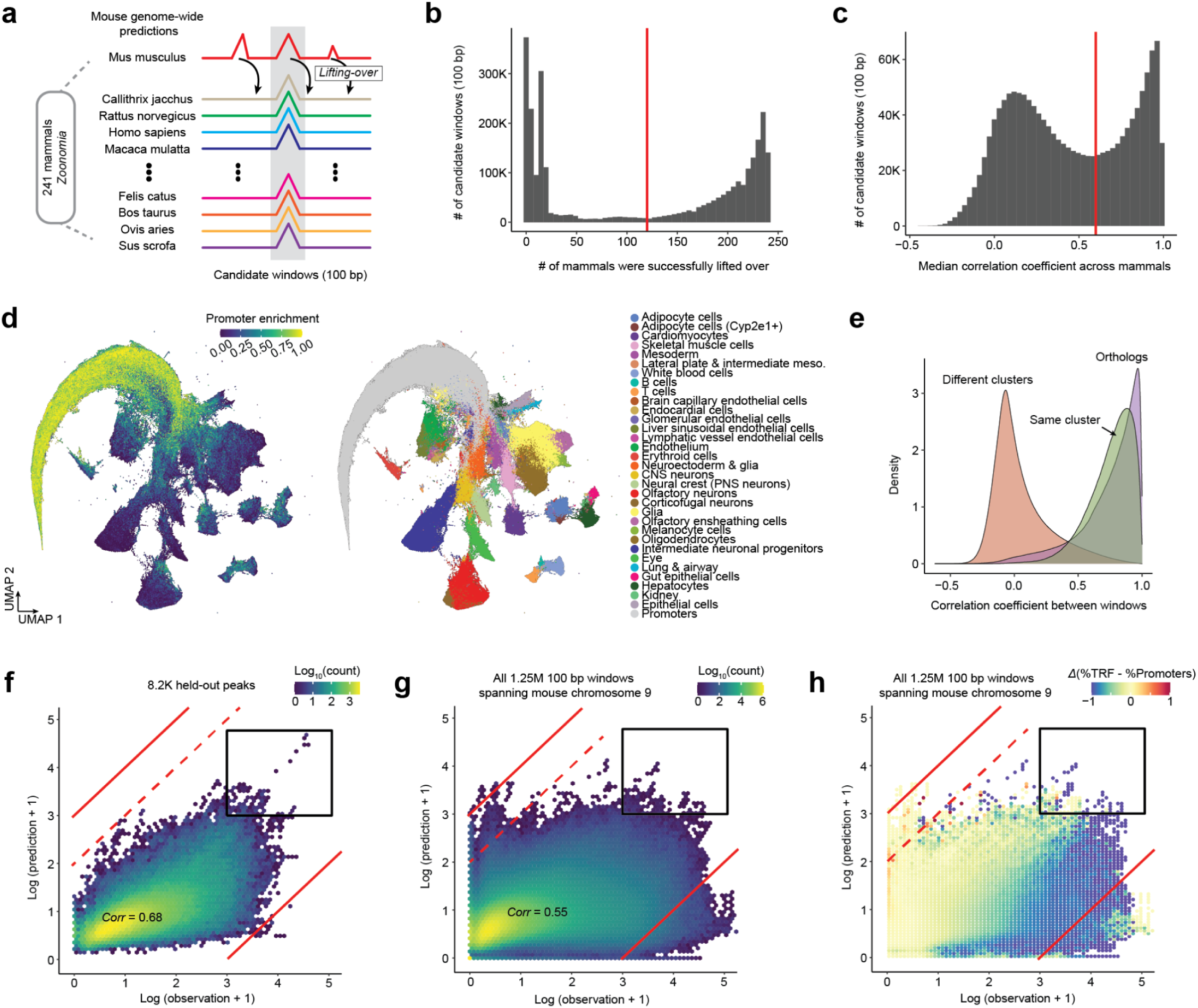
Evolutionary coherence separates distal enhancer grammar from promoter & repeat-associated signals. **a,** Schematic. We performed mouse genome-wide inference with the evolution-naive model. Windows with high predicted accessibility and specificity were filtered for evolutionary retention and coherence of model predictions across Mammalia^3^. **b,** To 2.79M windows predicted to be highly (Z ≥3) and specifically (Z ≥3) accessible in ≥1 cell class, we performed liftover to 240 other mammalian genomes^3^. 1.47M windows were successfully lifted over to ≥120 other species (red vertical line). **c,** To those 1.47M windows, we applied the evolution-naive CREsted model to syntenic regions of other mammals, and calculated the pairwise Pearson’s *r* between the resulting 36-value predicted accessibility vectors for mouse vs. ortholog. Windows exhibiting a median correlation coefficient greater than 0.6 (red vertical line) across species were retained. **d,** Predicted score matrix for the remaining 547,317 windows × 36 cell classes, subjected to UMAP embedding and Leiden clustering. 2D UMAP, colored by promoter enrichment (left) or annotation (right), is shown. Promoter enrichment calculated as % of a window’s 30 nearest neighbors (Euclidean) within 2.5 kb of an annotated TSS. **e,** We randomly sampled 500 windows from each of the 32 non-promoter clusters and computed Pearson’s *r* between their 36-value predicted accessibility vectors vs. those of windows from the same cell class cluster (red), different cell class cluster (green), or syntenic orthologs (purple). The resulting distributions are visualized as density plots. **f,** Scatterplots of observed (normalized Tn5 cut-site counts; log-n scaled; *x*-axis) vs. predicted (evolution-aware CREsted model & tandem repeat mitigation heuristic; log-n scaled; *y*-axis) accessibility at 8,223 held-out peaks, shown for all 32 cell classes and colored by log_10_ scaled density. Similar to Fig. 3b, which was generated with the evolution-naive model. **g,** Same as panel **f**, but across 1.25M × 100 bp windows spanning mouse chr. 9 for 32 cell classes. Similar to Fig. 3g, which was generated with the evolution-naive model and without the tandem repeat mitigation heuristic. **h,** Same as panel **f**, but across 1.25M × 100 bp windows spanning mouse chr. 9 for 32 cell classes and with points colored by the difference between % of windows overlapping tandem repeats and % overlapping promoters (± 2.5 kb of annotated TSS). Red indicates tandem repeat enrichment. Blue indicates promoter enrichment. Similar to Fig. 3h, which was generated with the evolution-naive model and without the tandem repeat mitigation heuristic.

To organize these syntenically retained, evolutionarily coherent windows, we performed Leiden clustering on their 36-value predicted accessibility vectors, which resulted in 33 groups: one large cluster highly enriched for promoters (190,035 windows) and 32 smaller clusters (1,018-45,655 windows) that largely mapped one-to-one to our Level-2 cell classes (**Fig. 4d**; **Supplementary Fig. 10a**). A few closely related classes were not separable and their labels therefore merged (primitive & definitive erythroid → erythroid; oligodendrocyte progenitors & committed oligodendrocyte precursors → oligodendrocytes). A few others mapped to the promoter cluster and were discarded (NMPs; erythroid niche cells). After filtering ENCODE blacklisted regions, the 32 non-promoter clusters comprised 354,450 × 100-bp windows (74,621 regions spanning 35.4 Mb after merging adjacent windows within each cluster). Reassuringly, the strength of correlation between prediction vectors of mouse windows within a cluster approaches that of mouse windows vs. their orthologs (**Fig. 4e**).

We then trained an ’evolution-aware’ CREsted model to predict chromatin accessibility across the 32 non-promoter clusters, using the same chromosome splits as the evolution-naive model (initial training on the 10,000 most specific windows per cluster, fine-tuning on the top 3,000). As with the evolution-naive model, we once again zeroed out TSS-proximal accessibility during training to further suppress the learning of promoter grammar. To address tandem repeat-induced false positives, we introduced an *in silico* randomization of annotated tandem repeat sequences within each scored window during inference, which destabilizes their contribution to predicted signal while still allowing windows that overlap tandem repeats to be scored (**Supplementary Fig. 10b**).

We performed genome-wide inference on mm10 with the evolution-aware model. Of note, although the resulting predictions exhibit poorer performance than the evolution-naive model in predicting accessibility at held-out peaks (*r =* 0.74 vs. 0.68; **Fig. 3b** vs. **Fig. 4f**) and genome-wide (*r =* 0.71 vs. 0.55; **Fig. 3g** vs. **Fig. 4g**), this reflects the deliberate suppression of promoter and other off-target regulatory grammars — signals that inflate correlation with observed accessibility but are not the intended targets of inference. In particular, the evolution-aware model substantially downweights promoter accessibility (**Fig. 4g,h**), eliminating their dominance among the most highly predicted regions upon genome-wide inference (black-boxed regions in **Fig. 3g,h** vs. **Fig. 4g,h**). Furthermore, we observe a marked reduction in tandem repeat-associated overprediction (upper-left quadrant of **Fig. 3g,h** vs. **Fig. 4g,h**; **Supplementary Fig. 10c**). These results further support our conclusion that both key failure modes encountered with the evolution-naive model are substantially mitigated by the evolution-aware model.

To summarize, the evolution-aware model substantially mitigates the failure modes of its predecessors. On its own, our filtering and reclustering of the training set to syntenically retained, functionally coherent windows failed to substantially reduce confounding by promoters (**Supplementary Fig. 7d**, column 2). But when combined with the zeroing of TSS-proximal accessibility, the proportion of the 0.01% highest scoring windows per cell class on held-out chromosome 9 that overlap promoters substantially dropped, while also stabilizing across thresholds (**Supplementary Fig. 7e**, column 2). Yet neither measure addressed our second confounder, as most highly scoring non-promoter windows still overlapped tandem repeats (**Supplementary Fig. 7e**, column 3). Only with all three measures in place (pre-training synteny/coherence-based filtering/reclustering, pre-training exclusion/zeroing of TSS-proximal regions, and *in silico* repeat perturbation during inference), do we achieve relief from confounding by promoters and tandem repeats (**Supplementary Fig. 7f**, columns 2-3). Although it comes at the cost in terms of performance in predicting accessibility at held-out peaks (*r =* 0.70 → 0.58), we reduce and stabilize the proportion of highest-scoring windows overlapping promoters from 97% to 39%, and the proportion of remaining high-scoring windows overlapping tandem repeats from 79% to 4% (**Supplementary Fig. 7a vs. 7f**, columns 1-3).

### Candidate developmental enhancers are conserved and predict cell type-specific gene expression

Because we apply the evolution-aware model genome-wide rather than to a predefined peak set, raw model outputs lack an intuitive reference frame for ranking predicted enhancers across the genome. To address this, we applied a scoring rubric similar to our Phred-scaled, non-parametric CADD scores for genome-wide variant effect prediction^66,67^. For each of the 32 cell classes, genomic windows were ranked by predicted accessibility, and for each score we defined P as the fraction of genomic windows with lower predicted values — a genome-wide percentile rank. We then converted these ranks into a <u>G</u>enome-wide <u>P</u>hred-like <u>S</u>core (GPS = −10 × log₁₀[1 − P]), facilitating their immediate interpretation (*e.g.* GPS 30 = top 0.1% of genome-wide predictions). To call peaks, we focused on the top 100,000 × 100-bp windows per cell class (10 Mb; GPS > 24.5; 0.355% of the genome). Adjacent high-scoring windows were merged, and for each merged region we extracted a 2,114-bp sequence centered on the region (matching the model input length), whose predicted accessibility was assigned to the region. Finally, we applied a trimming heuristic to identify the ‘core region’ of each candidate developmental enhancer, defined as the smallest contiguous subsequence whose predicted accessibility is at least 90% of the full 2,114-bp score (**Supplementary Fig. 10d**). Across all 32 cell classes, this procedure yielded 318,423 core candidate developmental enhancers (mean ± SD: 530 ± 316 bp) whose union spans 65.4 Mb (2.3%) of the mouse reference genome (**[Supplementary File 2](https://shendure-web.gs.washington.edu/content/members/cxqiu/public/backup/jax_atac/download/Supplementary_File_2_Peaks_Specific_to_32_Clusters.bed.gz)**).

Although the evolution-aware model was trained on chromatin accessibility, we introduced modifications to isolate distal enhancer grammars (training on syntenically retained, evolutionarily coherent peaks; zeroing out TSS-adjacent regions) and to mitigate overprediction artifacts (tandem repeat randomization). A focused examination of ∼100 kb encompassing the fetal hepatocyte marker genes *Alb* and *Afp* illustrates the impact of these adjustments (**Fig. 5a**). Observed chromatin accessibility is minimal in erythroid cells, while hepatocytes exhibit prominent peaks at promoters and candidate distal enhancers, together with accessibility throughout their highly transcribed gene bodies (**Fig. 5a**, top row-pair). The evolution-naive CREsted model recapitulates peaks at putative distal enhancers, but also peaks at both the *Alb* and *Afp* promoters in hepatocytes, as well as at regions inaccessible in both cell types, some of which, including the prominent peak in the erythroid track, are coincident with tandem repeats (**Fig. 5a**, second row-pair).

**Figure 5.**
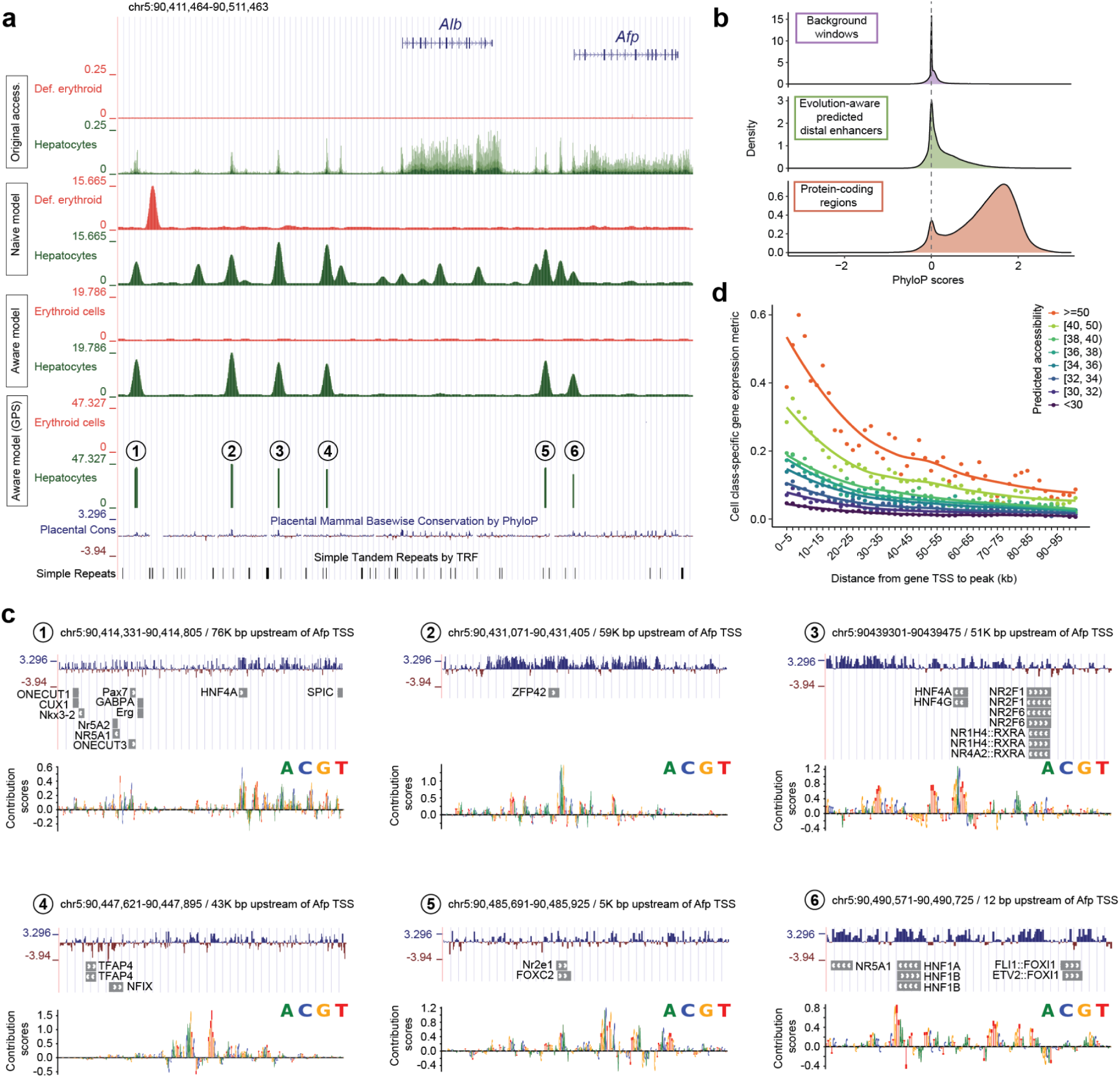
Evolution-aware modeling identifies distal enhancers that predict cell-type-specific gene expression. **a,** Genome browser view of 100 kb region encompassing *Alb* and *Afp* loci (mm10, chr5:90,411,464-90,511,463). Four pairs of definitive erythroid and hepatocyte tracks are shown (top to bottom): normalized Tn5 cut-site counts (pseudobulk), evolution-naive predicted accessibility, evolution-aware predicted accessibility, and GPS-scaled, trimmed enhancer predictions derived from the evolution-aware model. Additional annotation tracks are shown above (gene structure) and below (placental conservation, simple tandem repeats). **b,** Distribution of phyloP scores (60 placental mammals), computed by averaging nucleotide-level scores within each element, for background regions (n = 300,000 randomly selected 200-bp genomic windows; top row), evolution-aware, core-trimmed distal enhancer predictions (n = 318,423; middle row) and protein-coding regions (n = 262,025; bottom row). **c,** Nucleotide-resolution features of the 6 hepatocyte candidate enhancer core regions shown in bottom track of panel **a**. Tracks show basewise conservation (phyloP; 60 placental mammals) and model-derived saliency scores (evolution-aware model, CREsted: *crested.tl.contribution_scores*). **d,** Relationship between cell-class-specific enhancer prediction scores vs. cell-class-specific expression of nearby genes. We regrouped our mouse developmental scRNA-seq timelapse data^42^ into 32 cell classes, and generated normalized pseudobulk expression profiles for each. For each of ∼22,000 protein-coding genes across 32 cell classes, cell-class-specific expression was defined as the log_2_ fold-change relative to the gene’s median expression across all cell classes. Each of the 318,423 cell-class-specific enhancer predictions of the evolution-aware model was linked to each gene within 100 kb, and such pairings assigned the corresponding cell-class-specific expression value. Enhancer-gene pairs were then jointly stratified by distance (*x*-axis; 5-kb bins with 2-kb sliding steps) and GPS (colors). Within each distance-score bin, the *y*-axis shows the mean cell-class-specific gene expression across all enhancer-gene pairs in that bin, after subtracting a background estimate obtained from 100 permutations in which enhancer-cell class assignments were randomly shuffled. Smoothed trends were fit using *geom_smooth* in ggplot2.

In contrast, the evolution-aware model yields a markedly simplified landscape at this locus: six discrete hepatocyte-specific peaks (**Fig. 5a**, third row-pair), trimmed to six core elements (**Fig. 5a**, fourth row-pair; **Supplementary Fig. 10d**; lengths 154 to 474 bp). Tandem-repeat-coincident predictions and the *Alb* promoter peak are eliminated. One candidate enhancer (#6), lying 12-166 bp upstream of the *Afp* TSS (**Supplementary Fig. 11a**), is predicted as hepatocyte-specific, raising the question of whether it represents a ‘very promoter proximal’ enhancer or a cell-type-specific promoter — a distinction we revisit in the next section.

Genome-wide, the 318,423 candidate developmental enhancers exhibit nucleotide constraint intermediate between background and coding regions (**Fig. 5b**). Within the six candidate hepatocyte enhancers at the *Alb-Afp* locus, high-saliency positions^2,68^ often coincide with bases exhibiting high phyloP scores^69,70^ and predicted binding sites for hepatocyte-related transcription factors (**Fig. 5c**). This pattern is also observed at the *Tcf20* and *Sox2* loci, which, in contrast with *Alb-Afp,* harbor a diversity of cell classes among their predicted enhancers (**Supplementary Fig. 11b-c**). Although high-saliency bases (> 0.1) within predicted developmental enhancers comprise only 1%, 3% and 3% of bases at the *Alb-Afp*, *Tcf20* and *Sox2* regions, they account for 8%, 20% and 7% of nucleotides with phyloP > 2, respectively, after excluding coding sequence.

To evaluate whether enhancers predicted by the evolution-aware model inform cell-class-specific gene expression, we matched our mouse developmental scRNA-seq annotations^42^ to the 32 cell classes and calculated cell class-specific expression vectors for ∼22,000 protein-coding genes as their log_2_-fold change relative to the gene’s median across all cell classes. Candidate enhancers were binned by distance to each TSS and GPS. Aggregate plots show a clear dependence of gene expression on both predicted enhancer strength and proximity (**Fig. 5d**; **Supplementary Fig. 11d**), with both contributing independently within a given cell class. This result supports that evolution-aware model-predicted enhancers inform gene expression across major developmental lineages, and yields an aggregate estimate of distance-dependent decay in enhancer influence — albeit one confounded by genome architecture, as apparent decay with distance may partly reflect the increasing probability of intervening insulators or boundary elements. Notwithstanding this caveat, these results demonstrate that single-cell chromatin accessibility data, coupled with evolution-aware modeling, yields cell class-specific regulatory grammars that predict *in vivo* patterns of cell type-specific gene expression at the scale of the whole organism.

### Evolutionary augmentation enhances the robustness of cross-species inference

Although the evolution-aware model performed to our satisfaction for mouse-genome-wide inference of cell class-specific distal regulatory grammars, when we sought to apply it in a cross-species manner, the resulting predictions were miscalibrated. For example, human orthologs of the 354,450 mouse windows scored in a correlated but attenuated manner (**Supplementary Fig. 12a-b**; Pearson’s *r* = 0.50; log_2_ fold-difference *Hs*/*Mm*: median -0.64, mean: -0.59). Further concerning, when we applied it to a 200-kb window centered on the *Afp* TSS across 136 mammalian genomes for which this region was contiguously assembled (the 100 kb region shown in **Fig. 5a**, extended ∼20 kb upstream and ∼80 kb downstream; mean recovery: 191 ± 21 kb), many of the hepatocyte-specific enhancers predicted in mouse were largely restricted to other *Mus* species (**Supplementary Fig. 12c**). Regardless of the cause(s) (*e.g.* subtle differences in genome composition; genuine evolutionary divergence), this miscalibration challenges our goal of cross-species inference.

At the same time, in planning an advance from cell class (n = 36) to cell type (n = 140) resolution, we realized that many cell types and even some cell classes are represented by modest numbers of type/class-specific enhancers. Together, these observations highlight a core constraint: robust inference of mammalian *cis*-regulatory landscapes across hundreds to thousands of cell types likely requires a substantially larger and more diverse training corpus than any single genome can provide.

To overcome this, we sought to leverage the functional coherence of syntenic enhancers in a deeper way. Functionally constrained orthologous enhancers densely sample permissible sequence variation while preserving regulatory activity, and synteny enables their identification even in the face of substantial sequence divergence (**Fig. 4e**; **Fig. 5b**)^47,48^. We therefore expanded our training corpus to include syntenic orthologs of mouse-predicted enhancers from other mammals, increasing sequence diversity by ∼200-fold while preserving cell class-specific labels (**Fig. 6a**) — analogous to the role of diverse MSAs in protein structure prediction.

**Figure 6.**
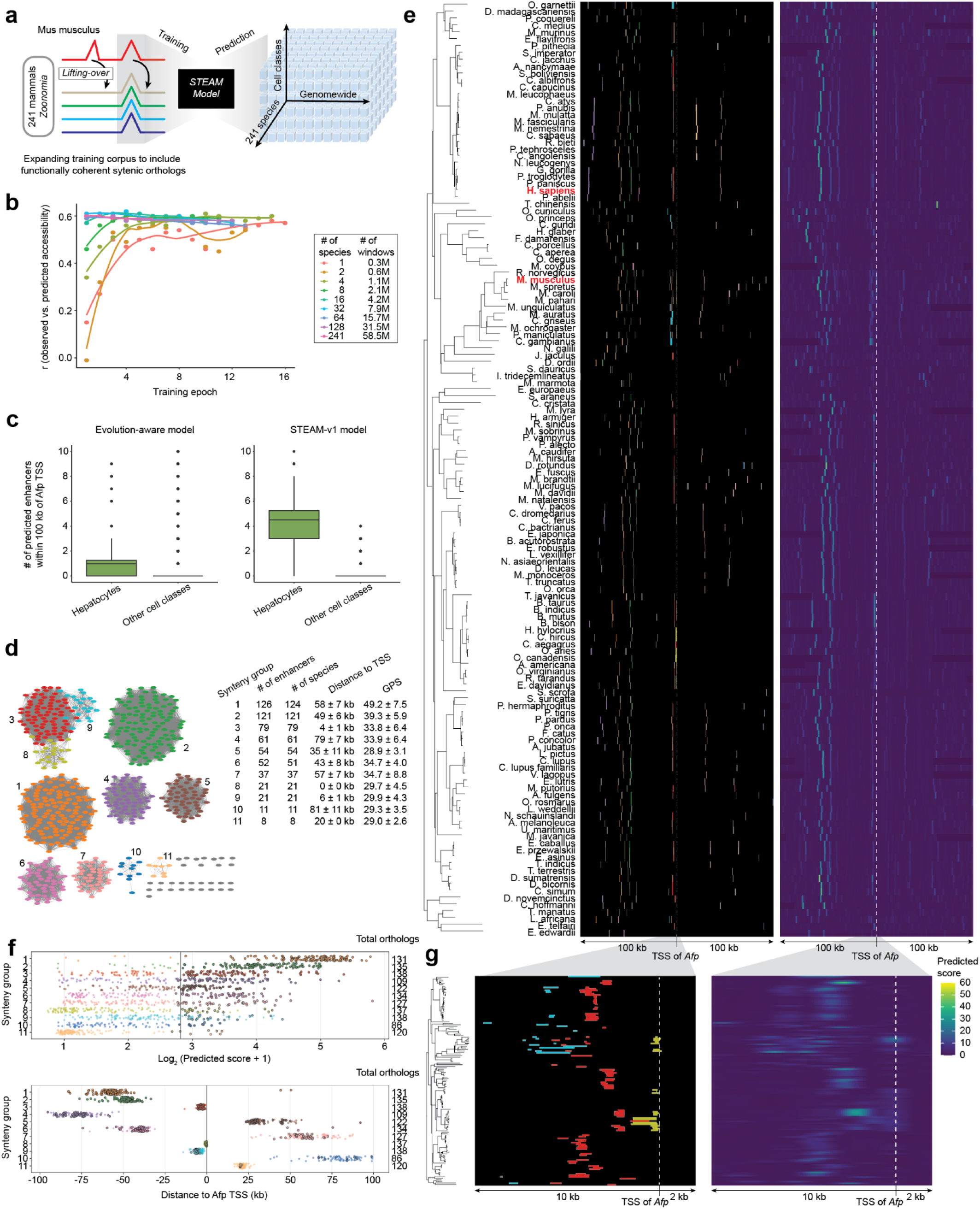
Evolutionary augmentation generalizes genomewide regulatory grammar inference across Mammalia. **a,** Evolutionarily coherent, syntenic orthologs of 354,450 × 100-bp mouse windows from 241 Zoonomia mammals^3^ were incorporated to train STEAM-v1, which was then applied to generate enhancer predictions spanning 32 cell classes for a 200 kb region centered on the *Afp* TSS in 136 species (panel **e**), as well as for whole-genome inference (human hg38/hg19, mouse mm10, and 239 additional Zoonomia genomes^3^). **b,** Models were trained on mouse windows and qualifying orthologs from progressively more species (1, 2, 4, …, 128, 241), expanding the corpus from 0.3M to 58.5M sequences. The 1-species model used mouse only; the 2-species model added human; subsequent models added randomly selected mammals. All models trained for up to 20 epochs with early stopping, except the 128- and 241-species models (manually stopped at epochs 12 and 6). Performance was evaluated at each checkpoint on a held-out mouse set of 22,014 sequences. **c,** The mouse *Afp* TSS was lifted over to 240 Zoonomia^3^ genomes and ±100 kb extracted; 136 species with ≥50 kb of contiguous recoverable sequence on each side were retained. The evolution-naive (left) and STEAM-v1 (right) models were applied to each syntenic locus (tiling, tandem repeat mitigation, GPS-based enhancer calling calibrated on mouse genome-wide scale, model-based trimming), and hepatocyte enhancer counts per species compared against the other 31 cell classes. Boxplot center lines: medians; box limits: 25th–75th percentiles. **d,** 621 hepatocyte enhancers predicted by STEAM-v1 across 136 species within 200 kb of the *Afp* TSS were lifted over between species; overlapping pairs were considered connected. The resulting graph forms 11 major synteny groups (clusters 1–11); remaining enhancers form smaller groups or singletons (grey). For each major synteny group: number of predicted enhancers, number of species represented, mean distance to *Afp* TSS, and mean GPS score are reported. **e,** Predicted hepatocyte chromatin accessibility from STEAM-v1 across 136 mammalian species within 200 kb of the *Afp* TSS. Contigs harboring the *Afp* TSS were reoriented to match the top-strand (forward) orientation of the *Afp* transcript in mm10. Left: phylogenetic tree for the 136 species. Middle: 621 hepatocyte enhancers colored by synteny group (as in panel **d**). Right: Predicted hepatocyte chromatin accessibility. **f,** For each of the 11 major synteny groups, orthologs in species lacking a called enhancer from that cluster were lifted over, merged, and treated as non-enhancer elements, yielding 591 enhancers and 786 non-enhancer elements. Top: log2-scaled predicted hepatocyte chromatin accessibility for enhancers (black-bordered) and non-enhancers, one row per synteny group. Bottom: distances to the *Afp* TSS for enhancers (black-bordered) and non-enhancers, one row per synteny group. In both plots, colors as in panel **d**. **g,** Subview of panel **e** spanning -10 kb to +2 kb of the *Afp* TSS across 136 species.

Specifically, starting from the 354,450 mouse windows used to train the evolution-aware model, we incorporated orthologs from 240 additional mammals^3^ that satisfied: (i) syntenic retention (successful liftOver and expansion to 2,114 bp) and (ii) functional coherence (pairwise Pearson’s *r* ≥ 0.6 between evolution-naive prediction vectors; **Fig. 4b,e**). Orthologs were assigned the chromatin accessibility profiles of their corresponding mouse sequence, inevitably introducing label noise. However, we reasoned that the value of expanded sequence diversity would outweigh this noise — analogous to the expectation that a model trained to identify bicycles based on 20,000 low-resolution images taken will outperform a model trained on 100 high-resolution images, provided signal-to-noise remains sufficient.

We trained a series of ‘evolution-augmented’ models on the 10,000 most cell class-specific mouse windows and their qualifying orthologs, with increasing numbers of species (1, 2, 4, … 128, 241) expanding the corpus from 0.3M to 58.5M sequences (195-fold). The 1-species model was trained on mouse only, the 2-species model added human orthologs, and subsequent models added orthologs from additional randomly selected mammals. Models were trained for up to 20 epochs with early stopping based on validation performance, except for the 128- and 241-species models, which were manually stopped at epochs 12 and 6, respectively. Chromosome-based train/validation/test splits were shared between mouse and orthologs to prevent leakage, and mouse TSS-proximal regions were zeroed out as before, now together with their syntenic orthologs.

Performance on held-out mouse sequences improved with greater species inclusion but plateaued around 32 species (**Fig. 6b**). Notably, the primary impact of corpus expansion was to accelerate convergence rather than improve performance, possibly reflecting inherent limits to predicting accessibility from 2,114-bp windows alone. We selected the best-performing checkpoint on the validation dataset (32-species, epoch 4) and fine-tuned on the top 3,000 most cell-class-specific mouse windows and their orthologs, yielding the Synteny-aware Transfer learning for Enhancer Activity Modeling model (STEAM-v1).

To assess whether STEAM-v1 mitigates cross-species miscalibration, we applied it to the 200-kb region centered on the *Afp* TSS across 136 mammalian genomes for which the evolution-aware model had failed to predict hepatocyte-specific enhancers beyond *Mus* (**Supplementary Fig. 12c-d**). For each syntenic locus, we performed predictions as previously described, including tiling, tandem repeat mitigation, GPS-based enhancer calling (calibrating on mouse genome-wide scale), and model-based trimming (**Supplementary Fig. 10b,d**).

The results were striking. While the evolution-aware model predicts 1.2 ± 1.7 hepatocyte enhancers per species (0.4 ± 1.5 for other cell classes), STEAM-v1 predicts 4.6 ± 1.7 (0.3 ± 0.5 for other cell classes) (**Fig. 6c**). As we have a strong expectation of hepatocyte-specificity at this locus, this corresponds to a 3-fold (evolution-aware) vs. 15-fold (STEAM-v1) signal-to-noise ratio. Moreover, the strong bias of the evolution-aware model for *Mus* was substantially mitigated with STEAM-v1 (mean predicted hepatocyte enhancers within *Mus* vs. all species: 8.2 vs. 1.2 (evolution-aware), 8.5 vs. 4.6 (STEAM-v1). Importantly, the substantial improvements in the consistency of cross-species inference with STEAM-v1 are not limited to the Afp locus, and we showcase the additional example of the β-globin LCR in **Supplementary Fig. 13**,

Altogether, 621 hepatocyte enhancers in 136 species are predicted by STEAM-v1 at the *Afp* locus (vs. 35 ± 19 for other cell types). However, 591 of these (95%) belong to 11 synteny groups, *i.e.* they have shared ancestry (**Fig. 6d**). To better understand these groups, we plotted their predicted accessibility scores and relative positions, together with those of an additional 570 orthologs from these same species and synteny groups that are not predicted as hepatocyte-specific enhancers (**Fig. 6e-f**). For some synteny groups, there is a clear separation of predicted-functional vs. predicted-non-functional orthologs that correlates with their phylogenetic distribution. For example, synteny group #11 contains only 8 predicted hepatocyte enhancers; these have STEAM prediction scores that are well separated from 102 non-functional syntenic orthologs and moreover all derive from old world monkeys (**Fig. 6e-f**). For other synteny groups, there is a more continuous distribution of predicted accessibilities that spans our threshold, with called enhancers more broadly distributed throughout the phylogeny (*e.g.* synteny group #5; **Fig. 6e-f**). As our threshold for calling enhancers is somewhat arbitrary, we cannot conclude with confidence whether elements falling just short of this threshold are nonfunctional.

Three synteny groups (#3, #8, #9) reside within 10 kb upstream of the *Afp* TSS (**Fig. 6f**). Upon closer inspection, synteny group #8 contains the aforedescribed *Afp* promoter-proximal enhancer (**Fig. 6g**; candidate enhancer #6; **Supplementary Fig. 11a**), centered 300 +/- 337 upstream of *Afp* TSS. The hepatocyte GPS scores for this synteny group follow a long-tailed distribution across its 137 orthologs (median: 6.3; MAD: 8.8), with only 21 species (15%) exceeding the enhancer-calling threshold (median GPS among these: 26.7). Accordingly, candidate enhancer #6 in *Mus musculus* may be better classified as a ’very proximal enhancer’ rather than a cell-type-specific promoter, as its hepatocyte specificity appears separable from the *Afp* TSS. This case also highlights the importance of distinguishing enhancer vs. promoter vs. other grammars, rather than solely focusing on accessibility as the target of inference.

### STEAM-v1 enables inference of human genome-wide enhancer landscapes

We applied STEAM-v1 to the human (hg38) and mouse (mm10) reference genomes at 100-bp tiling resolution to generate genome-wide distal enhancer prediction tracks across 32 cell classes, which collectively encompass all major lineages of the developing mammal (*HumMus-v1* tracks). Repeating the human-mouse ortholog comparison previously performed on the 354,450 evolution-aware windows (**Supplementary Fig. 12a-b**), we find that both correlation and attenuation are substantially improved with STEAM-v1 (Pearson’s *r =* 0.67; log_2_ fold-difference *Hs*/*Mm*: median -0.23, mean: -0.28; **Supplementary Fig. 12e-f**). Similar comparisons on held-out windows from mouse chromosome 9 confirm a marked improvement in interspecies agreement at syntenic windows with evolutionary augmentation, with the mean log_2_ fold-difference for *Hs*/*Mm* predictions dropping from -0.21 to -0.07, the proportion of windows in which *Mm > Hs* falling from 63% to 53%, and the proportion of *Mm* vs. *Hs* prediction vectors at called enhancers exhibiting strong correlation (Pearson’s *r >* 0.8) rising from 32% to 41% (**Supplementary Fig. 7f-g**, columns 4-5).

Across all cell classes, STEAM-v1 predicts 341,552 mouse enhancers (470 ± 288 bp) and 336,378 human enhancers (542 ± 310 bp), in aggregate covering 85 Mb and 90 Mb of the mouse and human genomes, respectively (**[Supplementary File 3](https://shendure-web.gs.washington.edu/content/members/cxqiu/public/backup/jax_atac/download/Supplementary_File_3_Evolution_Augmented_Model_Predict_On_Mouse_Genome.bed.gz)**, **[Supplementary File 4](https://shendure-web.gs.washington.edu/content/members/cxqiu/public/backup/jax_atac/download/Supplementary_File_4_Evolution_Augmented_Model_Predict_On_Human_Genome.bed.gz)**). However, this close agreement is expected given our fixed-footprint peak-calling heuristic. Following liftover of mouse enhancer predictions to the human genome, 23% have a syntenic ortholog predicted as an enhancer for the same cell class, 49% have a syntenic ortholog that is not a matched prediction, 28% have no syntenic ortholog. Nearly identical proportions are obtained following liftover of human enhancer predictions to the mouse genome (24%, 49%, 27%).

Merging overlapping predictions yields 167,193 mouse (158,047 human) regions, of which 61% (62%) are predicted enhancers for a single cell class, 21% (21%) for two cell classes, and 17% (17%) for three or more cell classes (**Fig. 7a**). In both species, multi-class predictions are enriched for coincidence with TSSs (± 2.5 kb) (**Fig. 7b**), suggesting that the challenge of contaminating promoter grammar, although mitigated, may not be fully resolved. Remarkably, Jaccard-based co-occurrence analysis reveals a nearly identical structure for mouse and human enhancer predictions, with developmentally or functionally related lineages co-localizing most strongly (**Fig. 7c**).

**Figure 7.**
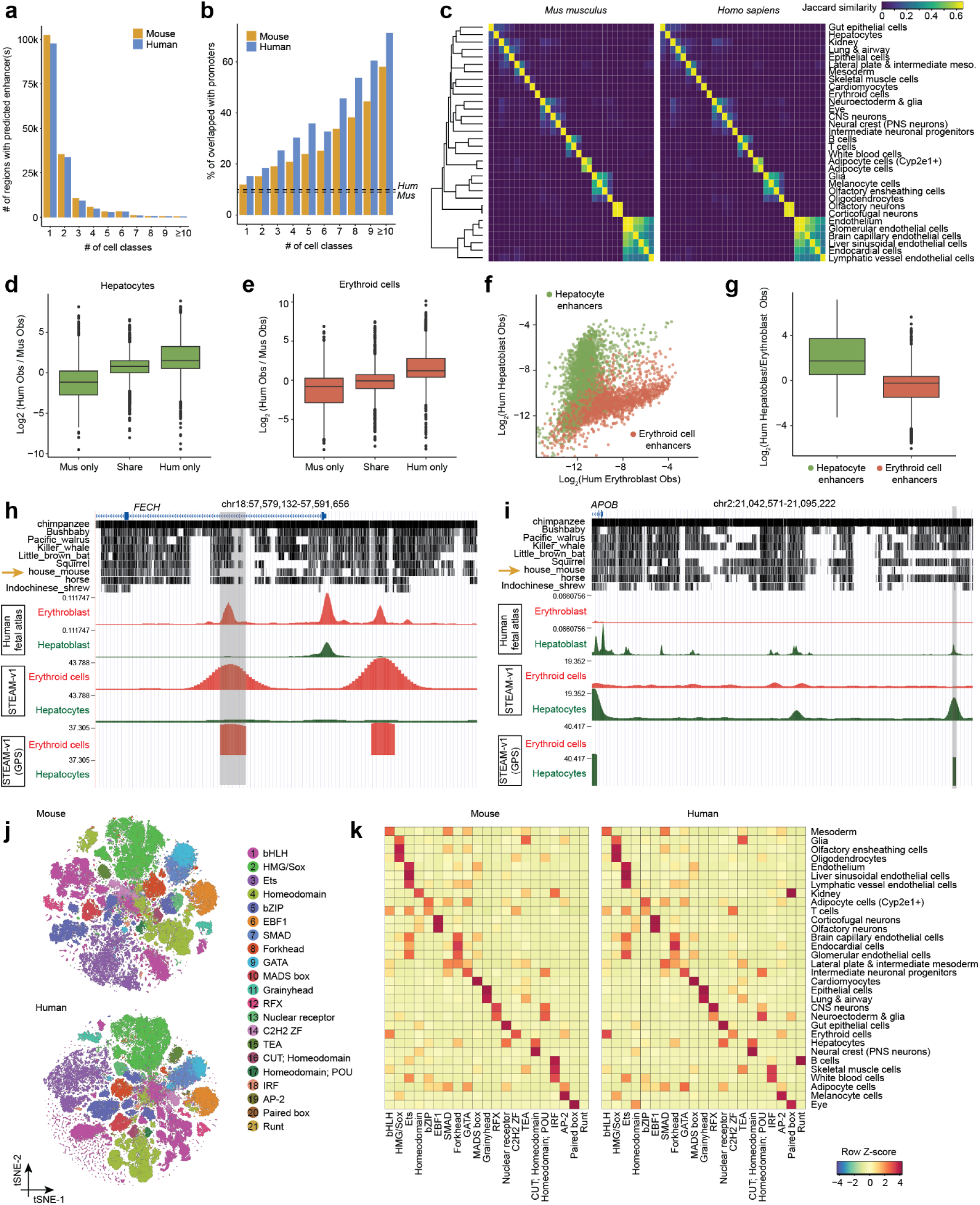
STEAM-v1 predicts human and mouse genome-wide enhancer landscapes. **a,** Genome-wide STEAM-v1 enhancer predictions were merged across cell classes, yielding 167,193 (mouse) and 158,047 (human) regions. Histogram of merged regions by the number of cell classes for which they are predicted enhancers, in mouse and human. **b,** For each category in panel **a**, the percentage of merged regions overlapping a promoter (± 2.5 kb around an annotated TSS) is shown for mouse and human. Background (dashed lines) was estimated from 10,000 randomly selected regions matched for size distribution. **c,** Jaccard similarity between all pairwise cell class combinations, calculated as the intersection over union of predicted STEAM-v1 enhancers, shown for mouse (left) and human (right). Merged groups composed of ≥7 cell classes (see panel **b**) were excluded from this analysis. The diagonal was set to the maximum non-diagonal value. Rows and columns for both species are ordered identically by hierarchical clustering on the mouse cell classes. **d,** Human and mouse predicted hepatocyte enhancers were reciprocally lifted over, and successful liftovers intersected to define three sets: mouse-only (n = 4,948), human-mouse shared (n = 2,597), and human-only (n = 5,790) predictions. Observed chromatin accessibility from human fetal liver hepatoblasts^53^ and mouse hepatocytes (this study) was extracted at averaged base resolution, and the distribution of log_2_-scaled human/mouse fold-changes is plotted for the three enhancer sets. Boxplot center lines: medians; box limits: 25th-75th percentiles. **e,** Same as panel **d** for erythroid enhancers: mouse-only (n = 3,492), human-mouse shared (n = 1,045), and human-only (n = 4,353) predicted enhancers with successful cross-species liftover. Observed chromatin accessibility from human fetal liver erythroblasts^53^ and mouse definitive erythroid cells (this study) was used. **f,** Human-specific hepatocyte (n = 3,239; green) and erythroid (n = 3,842; red) predicted enhancers failing liftover to the mouse genome are plotted by observed chromatin accessibility in human fetal liver erythroblasts (*x*-axis) vs. hepatoblasts (*y*-axis), log_2_-scaled^53^. **g,** For the same sets of human-specific enhancer predictions failing liftover to the mouse genome shown in panel **f**, the log_2_-scaled ratio of observed human hepatoblast to erythroblast accessibility is compared between hepatocyte-predicted (left) and erythroid-predicted (right) enhancers. Boxplot center lines: medians; box limits: 25th–75th percentiles. **h,** Genome browser view of a human erythroid-specific enhancer with no syntenic mouse ortholog, located in the first intron of *FECH*. Tracks show (top to bottom): Zoonomia 241-species alignment^3^, observed liver erythroblast and hepatoblast accessibility from the human fetal atlas^53^, STEAM-v1 predicted erythroid and hepatocyte accessibility, and GPS-scaled core enhancer predictions. Generated using the UCSC Genome Browser (hg38). **i,** Same as panel **h** for a human hepatocyte-specific enhancer with no syntenic mouse ortholog, located ∼50 kb upstream of the *APOB* TSS. **j,** For each class in the STEAM-v1 model, the top 2,000 class-specific mouse and human enhancers were selected by ranking their prediction scores. Contribution scores were then calculated, seqlets extracted using TF-MINDI^71^ and tangermeme^72^, and then scored against the SCENIC+ motif collection^73^. The resulting similarity matrix was clustered using PCA, nearest-neighbor graph construction, and Leiden clustering. Each cluster was annotated by its most prevalent TF DNA-binding domain family. t-SNE visualizations of the 300,748 mouse (top) and 295,871 human (bottom) seqlets are shown, labeled by each cluster’s assigned TF DNA-binding domain family. **k,** The frequency of seqlets from each cluster across cell classes (based on the class-specific enhancer in which each seqlet was identified) was calculated. Frequencies were first normalized by column totals and then converted to row-wise z-scores. Heatmaps for both mouse (left) and human (right) are shown, with rows and columns in the same order.

To assess the human STEAM-v1 predictions, we turned to the Domcke *et al.* human fetal chromatin accessibility atlas, which we previously generated from midgestational organs via sci-ATAC-seq3^53^. As the Domcke *et al.* pseudobulk tracks were generated on hg19, we also applied STEAM-v1 to hg19 at 100-bp tiling resolution, yielding 336,745 human enhancer predictions (**[Supplementary File 5](https://shendure-web.gs.washington.edu/content/members/cxqiu/public/backup/jax_atac/download/Supplementary_File_5_Evolution_Augmented_Model_Predict_On_Human_Genome_hg19.bed.gz)**). For our evaluation, we focused on erythroblast and hepatoblast accessibility profiles from fetal liver — the human counterparts of our mouse definitive erythroid and hepatocyte cell classes, respectively.

We first examined predicted human and mouse enhancers for these cell classes that have a syntenic ortholog in the other species — regardless of whether that ortholog was itself a predicted enhancer — as for these regions we have cell-class-matched observed accessibility data in both species. Reassuringly, predicted human-only enhancers were substantially more accessible than predicted mouse-only enhancers in the corresponding human cell class: an 8.8-fold difference for hepatocyte predictions, and an 8.5-fold difference for erythroid predictions (difference in geometric means; **Fig. 7d-e**), consistent with STEAM-v1 correctly predicting species-specific regulatory differences at syntenically shared regions.

As a more stringent test, we also examined predicted human enhancers that failed liftover to mouse entirely, *i.e.* regions for which there were neither human accessibility measurements nor even labeled mouse syntenic ortholog sequences available to STEAM-v1. As we also lack mouse accessibility measurements for these regions, we instead examined human hepatoblast/erythroblast accessibility ratios for both hepatocyte (n = 3,239) and erythroid (n = 3,842) predicted enhancers, and observe a mean 7.1-fold difference in the expected direction (difference in geometric means; **Fig. 7f-g**). Collectively, these results provide direct empirical validation of evolutionary transfer learning: STEAM-v1 infers human cell class-specific regulatory activity despite having been trained exclusively on mouse molecular data.

We highlight four examples of STEAM-v1 human-specific predicted enhancers with no mouse ortholog: (i) *<u>FECH</u>:* In the first intron of *FECH*, which encodes an erythroid-specific enzyme performing the final step of heme production, STEAM-v1 predicts a strong erythroid-specific enhancer (GPS 36.5), aligning precisely with an observed accessibility peak specific to human erythroblasts (**Fig. 7h**). Syntenic orthologs of this enhancer are intact in macaque, disrupted in other mammals, and absent in *Mus musculus.* (ii) *<u>TFRC</u>:* A similar case (erythroid GPS 36.1) resides ∼26 kb upstream of *TFRC*, which encodes the erythroid-specific, iron-uptake-essential transferrin receptor, here within a broader region lacking mouse synteny (**Supplementary Fig. 14a**). (iii) *<u>APOB</u>:* STEAM-v1 predicts a strong human hepatocyte enhancer (GPS 36.9) ∼50 kb upstream of the TSS of *APOB*, a key determinant of circulating LDL-cholesterol levels (**Fig. 7i**). This element is syntenically retained in nearly all mammalian genomes, with the notable exception of *Mus musculus*. (iv) *<u>CYP2C19</u>:* This locus encodes an enzyme that metabolizes many commonly prescribed drugs, and like many cytochrome P450 genes, it lacks a 1:1 mouse ortholog. Around 15 kb upstream of its promoter, STEAM-v1 predicts a strong hepatocyte-specific enhancer (GPS 34.7) which may explain *CYP2C19’s* liver-specific expression (**Supplementary Fig. 14b**). This is the only accessible peak, and the only STEAM-v1 predicted enhancer, in the ∼20 kb region between *CYP2C18* and *CYP2C19*, for which full synteny appears restricted to a subset of primates.

### Regulatory interpretation of the cell class-specific enhancer grammars of the STEAM-v1 model

As a preliminary confirmation that the grammars learned by STEAM-v1 matched expectation, we examined motif enrichment among predicted cell-class-specific mouse and human enhancers, and found reassuring agreement between the top enriched motifs and the key transcription factors associated with each cell class (**Supplementary Figs. 15-16**).

Next, we sought to extract *cis*-regulatory sequence-to-function rules with a recently developed computational package, TF-MINDI^71^. For each cell class in the STEAM-v1 model, we selected the top 2,000 mouse and enhancers, extracted seqlets (short, high-attribution subsequences representing motif instances) from model attribution scores, clustered them by similarity to known TF motifs, and mapped them back to cell classes to reveal the cell-class-specific regulatory grammar learned by the model (**Fig. 7j-k**).

Many of the observed enrichments align with known biology and are consistent between mouse and human, *e.g.* MADS-box (MEF2), a key driver of cardiac muscle identity, in cardiomyocytes; bHLH, consistent with TAL1/SCL and its erythroid partners, in erythroid cells; RFX, known for roles in neuronal gene regulation, in CNS neurons; and Grainyhead, important for epithelial differentiation and barrier function, in lung epithelial cells. We note that the apparent species-specific differences, *e.g.* differential enrichments of IRF (mouse > human) vs. RUNT (human > mouse) in B cell enhancers, may be due to the small size of these seqlet clusters (together less than 0.5% of total seqlets in each species).

### Genome-wide inference of distal enhancer landscapes of 241 mammals

Beyond the *HumMus* prediction sets for hg19, hg38, and mm10, we also applied STEAM-v1 to all Zoonomia genomes at 100-bp tiling resolution, which yielded a total of 32 □ 241 = 7,712 genome-wide enhancer prediction tracks (*BabaGanoush* tracks). These tracks were also subjected to peak calling, GPS calibration, and core-element trimming, yielding consistently generated enhancer prediction sets for each species. Finally, we leveraged the Zoonomia-associated Cactus alignments to generate a [UCSC Genome Browser track](https://genome.ucsc.edu/cgi-bin/hgTrackUi?hgsid=4021021427_ycyuSO7Aigo6JalKfHaAYVIb4veg&c=chr9&g=hub_7227887_jaxAtacMammals) that allows users to explore distal enhancer landscapes across all 241 mammalian genomes in a manner that is anchored to the human genome.

## DISCUSSION

Here we combine three key ingredients — (i) a 3.9 million single cell atlas of whole embryo chromatin accessibility spanning mouse organogenesis through birth (reported here), (ii) a multi-output convolutional neural network framework (CREsted^2^), and (iii) hundreds of mammalian genomes with defined synteny blocks (Zoonomia^3^) — to learn the deeply conserved regulatory grammars of major mammalian cell lineages and to infer distal enhancer landscapes genome-wide. A practical advantage of mouse is that the entire organism can be profiled as an intact whole, enabling all cell types to be sampled as they emerge and mature across tightly staged timepoints^41–45,74,75^. Although our initial motivation was to access enhancer landscapes for human cell types that are inaccessible to direct profiling, the resulting framework generalizes broadly, and we release genome-wide enhancer predictions across 32 cell classes for human and mouse (*HumMus*; hg19, hg38, mm10) and for 239 additional mammalian genomes (*BabaGanoush*; 7,712 tracks total).

Arriving at the STEAM framework required iterative diagnosis and resolution of failure modes that emerged at successive stages of genome-wide and cross-species inference. An evolution-naive CREsted model trained on the mouse atlas performed well on held-out peaks but failed in genome-wide inference, as the highest-scoring predictions were dominated by tandem repeats and promoter-proximal sequences, neither of which encodes the distal enhancer grammars we sought to learn (**Supplementary Fig. 7a-c**). The combination of three measures — filtering training windows for syntenic retention and functional coherence across mammals, zeroing TSS-proximal accessibility during training, and randomizing tandem repeat sequences *in silico* during inference — addressed these failures, but cross-species inference remained miscalibrated (**Supplementary Fig. 7d-f**). STEAM-v1 resolved this by augmenting training with the syntenic orthologs of mouse-predicted enhancers across other mammals, expanding effective sequence diversity by as much as ∼200-fold while preserving cell-class labels (**Supplementary Fig. 7g**). The result is analogous to the role of multiple sequence alignments in protein structure prediction. Orthologous enhancers, like orthologous protein sequences, can be treated as diverse realizations of a deeply constrained grammar, and the mismatch between the tempos of *cis-*vs. *trans-* regulatory evolution provides the supervisory signal that single-genome models lack. By scaling this supervisory signal across hundreds of mammalian genomes, STEAM-v1 transforms a single-species molecular atlas into a framework capable of inferring regulatory grammars throughout placental mammals.

A central motivation for this work is the difficulty of interpreting human regulatory variation across the full diversity of *in vivo* cellular states, given the limited human contexts accessible to functional experiments^76^. Progress on the causal variants underlying tens of thousands of GWAS associations has arguably stalled on precisely this constraint. Predictive models that capture *cis*-regulatory grammars across the full range of cell types offer a path through it, enabling *in silico* exploration of variant effects in contexts that cannot be directly profiled, including the embryonic, fetal, and pediatric stages in which most human cell types arise. In this view, large-scale functional genomics datasets serve not merely as end products but as training substrates for increasingly generalizable models of context-aware gene regulation, learned by evolutionary transfer from model organisms whose developmental biology can be profiled comprehensively. By learning from the temporal unfolding of mammalian development across many species simultaneously, we can extract a consensus ontogeny of mammalian regulation and project it onto the human genome to predict human-specific gene regulation, without ever requiring access to human embryonic, fetal, or pediatric tissues. Beyond variant interpretation, the BabaGanoush catalog itself constitutes a model-driven resource of unprecedented scale, as the 7,712 cell-class-specific enhancer tracks across 241 mammals will serve as a systematic hypothesis generator for where enhancers have been gained, lost, redeployed across lineages, or rewired between cell types — questions previously addressable only at a handful of loci or for a small number of species pairs.

Several limitations of this work warrant emphasis. <u>First</u>, the evolutionary coherence filter used to curate training data relies on predictions from the evolution-naive model itself, raising the risk of circularity. We do not regard this as vicious; because the filter is applied independently across hundreds of diverse genomes, mouse-specific spurious predictions are unlikely to be consistently reinforced. The cross-species calibration improvements observed with STEAM-v1, including predictions at human-only enhancers that validate against held-out human fetal chromatin accessibility data, also argue against systematic propagation of mouse-specific bias. However, a more rigorous evaluation would involve cross-validation across both chromosomes and clades. <u>Second</u>, despite augmentation with up to 240 mammalian genomes, the training corpus is ultimately anchored on mouse chromatin accessibility labels, and some residual species bias remains, as evidenced by the mild but persistent attenuation of human predictions relative to mouse (**Supplementary Fig. 12e-f**). Matched atlases in a handful of phylogenetically strategic additional species would likely yield substantial improvements. <u>Third</u>, our heuristic for suppressing promoter grammar — zeroing chromatin accessibility within 2.5 kb of annotated TSSs during training — is a blunt instrument. We do not fully understand why it appears necessary, only that it was a critical ingredient for suppressing the confounding of enhancer and promoter grammars. Looking forward, a more principled replacement remains an important goal.

STEAM-v1 represents a first step, and several directions stand out as particularly promising. The first is scale. Beyond the nearly 4 million scATAC-seq profiles reported here, other large-scale mammalian studies contribute another ∼4 million covering cell type contexts and developmental stages not sampled by our atlas, including pre-implantation and early embryonic development, extraembryonic tissues, adult tissues, brain regions, and immune stimulation^50,53,75,77–83^. Extending STEAM to these data, while moving from cell class (n=36) to cell type (n=140) resolution, should yield an increasingly complete hierarchical graph of mammalian regulatory grammars that mirrors the ontogeny of cell types^42,74^. At the limit, this discrete hierarchy becomes continuous, analogous to a Waddingtonian landscape of cell type grammars. The same logic applies to the corpus of mammalian genomes, in that although superficial performance plateaus with the 241 mammals currently available, further expanding this corpus is likely to yield compounding benefits as we move to finer cellular resolution, where specialized grammars may be represented by far fewer enhancers per genome. Evolutionary transfer learning is also not inherently limited to mammals. Among vertebrates, birds offer hundreds of sequenced genomes^84^ but lack time-resolved single-cell molecular data, while teleost fishes like zebrafish offer rich developmental single-cell chromatin accessibility data^38,40^ but a sparser genomic corpus. Both gaps are addressable. Looking further ahead, frameworks that accommodate phylogenetic distance explicitly, allowing models to both constrain and permit regulatory divergence as a function of evolutionary separation, would integrate and extend the approach across deeper taxonomic distances. Taken to its limit, sufficiently dense collections of syntenic sequences may permit regulatory grammars to be inferred even without direct functional measurements, through unsupervised or weakly supervised approaches that exploit the tendency of syntenic sequences to preserve regulatory function.

Our results suggest a renewed and modernized framing of the role of model organisms in the study of human biology. Model organisms remain the foundation of molecular biology: nearly everything we know about human development, physiology, and disease at the level of genes, molecules, and cells has its roots in insights first made in mice, flies, worms, fish, yeast, or bacteria. Yet this knowledge transfer has historically been ad hoc, and has not been systematically scaled to the era of AI and machine learning. Evolutionary transfer learning provides a framework for systematically leveraging the experimental richness of model organism data, together with the growing diversity of sequenced genomes, to make principled inferences about human biology. The current study employs the genome sequences of ∼4% of extant mammalian species and a unimodal single-species molecular atlas spanning only half of mouse prenatal development. The headroom for improvement is therefore substantial. In this light, sequencing additional mammalian genomes and multimodal profiling the molecular, cellular, and developmental biology of key model organisms represent not merely exercises in cataloging biodiversity or satisfying scientific curiosity — though we wholeheartedly support both of those justifications in their own right — but critical investments in the data infrastructure needed to model human biology effectively. In short, Dobzhansky’s adage that nothing in biology makes sense except in the light of evolution is likely to be reinforced, not dampened, in the age of AI.

## AUTHOR INFORMATION

## Acknowledgments

We thank the members of the Shendure Lab for helpful discussions, R. Waterston (UW), D. Kelley (Calico) and F. Huang (Calico) for thoughtful feedback, R. Cockett (Curvenote) for piloting an interactive version of this work, and Q-T-π (*Canis familiaris*), Tater and Tot (*Feline catus*) for inspiration. We thank C. Cenik and Z. Ma (M. Snyder laboratory) for providing the CH12.LX cell line. This work was supported by the Brotman Baty Institute for Precision Medicine and grants from the Paul G. Allen Frontiers Group (Allen Discovery Center for Cell Lineage Tracing to J.S. and C.T.), Seattle Hub for Synthetic Biology, a collaboration between the Allen Institute, Chan Zuckerberg Initiative and University of Washington (award number CZIF2023-008738 to J.S. and C.T.), Alex’s Lemonade Stand Foundation (Grant 19-15730 to J.S.), Crazy 8 Initiative (to J.S.), and National Institutes of Health (R01HG010632 to J.S. and C.T.). I.C.W. and S.A.M were supported by N.I.H. grant UM1OD023222, R01DE032319, and the JAX Director’s Innovation Fund. C.Q. was supported by Dartmouth’s Center for Quantitative Biology through a grant from NIGMS (P20GM130454). J.S. is an Investigator of the Howard Hughes Medical Institute.

## Author Contributions

J.S., C.Q. designed the research.

I.C.W. collected and staged the mouse embryos, and wrote the corresponding parts of the manuscript.

R.M.D. performed the sci-ATAC-seq3 experiments and generated the data (with assistance from B.K.M., T-M.L., M.L.T., O.F., D.R.O.), and wrote the corresponding parts of the manuscript.

C.Q., R.P.P. performed all computational analyses. S.Y., T.L., S.D., C.T. assisted with data analysis. I.C.W., J.B.L., S.A.M., C.T. assisted with results interpretation.

N.K., S.A. developed the CREsted package and assisted with its implementation.

S.D.W., C.M., S.A. developed the TF-MINDI package and performed the corresponding analyses.

C.Q., J.S. collaboratively explored and annotated the data, and wrote the manuscript except for sections corresponding to mouse staging and data generation.

J.S. supervised the project.

## Data & code availability

An interactive version of this preprint is available at [this link](https://doi.org/10.62329/hxkk6249) (ref ^4^). Data, including count matrices, CREsted models, and prediction tracks, are available at https://chengxiangqiu.github.io/jax-atac-website. Code and scripts are available at https://github.com/ChengxiangQiu/jax-atac-code. Data can be downloaded in raw and processed forms from the NCBI Gene Expression Omnibus under accession number GSE325776.

## AI Disclosure Statement

We disclose that language editing, proofreading and coding were supported by AI-based tools; these were not used for conceptual development or primary manuscript writing. The authors take full responsibility for the contents of this manuscript.

## Competing Financial Interests Statement

J.S. is on the scientific advisory board, a consultant, and/or a co-founder of Guardant Health, Phase Genomics, Sixth Street Capital, Pacific Biosciences, Cellular Intelligence and 10x Genomics. C.T. is a consultant for 10x Genomics. All other authors declare no competing interests.

## METHODS

### Data reporting

For newly generated mouse data, no statistical methods were used to predetermine sample size. Embryo collection and sci-ATAC-seq3 data generation were performed by different researchers in different locations.

### Mouse embryos collection and staging

All animal use at The Jackson Laboratory was done in accordance with the Animal Welfare Act and the AVMA Guidelines on Euthanasia, in compliance with the ILAR Guide for Care and Use of Laboratory Animals, and with prior approval from The Jackson Laboratory animal care and use committee (ACUC) under protocol AUS20028.

The details of collecting 523 mouse embryos, ranging from embryonic day (E) 8.0 to P0, were described in a previous publication^42^. Briefly, C57BL/6NJ (strain #005304) mice were obtained from The Jackson Laboratory and maintained under standard husbandry conditions. Timed matings were set up in the afternoon, and vaginal plugs were checked the following morning. Noon on the day a plug was detected was designated as E0.5 and harvests were performed at intervals throughout the day or evening to ensure collection of samples within the desired 6 hour staging bins. At each developmental stage, individual decidua were removed and placed in ice cold PBS during harvest. Embryos were dissected free of maternal tissue and extraembryonic membranes, imaged, and then snap frozen in liquid nitrogen. Yolk sac tissue from each embryo was used to genotype for embryo sex, and all samples were stored at −80°C until further processing.

We employed a combination of staging approaches depending on the gestational age at harvest. To maximize temporal coherence, resolution, and accuracy, individual embryos were staged primarily on the basis of well-defined morphological criteria rather than gestational day alone (see ref ^42^ for full details). Briefly:

1. For E10.25 to E15.0, hindlimb bud morphology was used to estimate the absolute developmental stage of each sample via a combination of tools developed by Marco Musy in the Sharpe lab (embryonic Mouse Ontogenetic Staging System (eMOSS); https://limbstaging.embl.es/ and https://github.com/marcomusy/welsh_embryo_stager)^85,86^. Samples staged using eMOSS are denoted with “mE” to indicate morphometric embryonic day (*e.g.* mE13.5).
2. For E15.0 to E16.75, embryos were manually staged into bins based on changes in digit morphology as defined in ref ^42^.
3. Embryos from E17.0 to E18.75 were staged solely by gestational age.
4. P0 samples were harvested from litters at noon of the day of birth (parturition for C57BL/6NJ occurs between E18.75 and E19.0).

The images of individual embryos at each timepoint can be found in **Extended Data Fig. 1** of ref ^42^. Of note, only 36 samples from E10.0 to P0 were profiled using sci-ATAC-seq3 in this study.

*Generating data using an optimized version of sci-ATAC-seq3*

A total of six sci-ATAC-seq3 experiments were performed on 36 individual mouse embryos. At least one sample was included for every ∼6 hour time interval from E10.0 to P0, except for the timepoint at E17.75. We tried to ensure that as many female mice as possible appear (29 females vs. 7 males) to avoid potential bias caused by sex differences, which we observed in a previous study^87^. A detailed summary of individual embryos can be found in **Supplementary Table 1**.

We generated the dataset using the sci-ATAC-seq3 protocol as described in a recent study^53^. The detailed protocol is also available at: https://www.protocols.io/view/sci-atac-seq3-ewov18xn7gr2/v1/v1. Briefly, snap-frozen tissues were pulverized and processed for nuclei fixation then banked for future experiments. On the day of experiment, frozen fixed nuclei were thawed, repermeabilized in Omni lysis buffer^88^, and diluted in ATAC-resuspension buffer (RSB: 10 mM Tris-HCl pH 7.4, 10 mM NaCl, 3 mM MgCl₂) supplemented with 0.1% Tween-20. For three-level indexing experiments at 384 × 384 × 384 resolution, 4.8 million nuclei (50,000 per well) were distributed across 96 wells. In this study, each experimental batch varied in the number of tissues per experiment to aim for at least 100,000 nuclei for early embryonic timepoint and increasing recovery for late developmental stage tissue. As per previous study, a mixture of GM12878 human and CH12-LX mouse cells were included as internal barnyard control.

For each tissue sample, nuclei were pelleted and resuspended in tagmentation master mix (Nextera TD buffer, 1X DPBS, 0.01% Digitonin, 0.1% Tween-20), aliquoted into wells (50,000 nuclei per well) of LoBind plates, and 2.5 μl Nextera v2 enzyme was added. Plates were incubated at 55°C for 30 min, and reaction was stopped by adding a stop solution (40 mM EDTA, 1 mM Spermidine) then incubated at 37°C for 15 min. Tagmented nuclei were pooled, pelleted, washed, and resuspended in ATAC-RSB with 0.1% Tween-20.

Phosphorylation master mix (1X PNK buffer, 1 mM rATP, T4 PNK) was added to tagmented nuclei, and distributed across four 96-well LoBind plates (12,500 nuclei per well) and then incubated at 37°C for 30 min. Ligation master mix (1X T7 ligase buffer, N5_splint, T7 DNA ligase) and N5 oligos (384 distinct barcodes) was added and incubated at 25°C for 1 hour, then stop solution (40 mM EDTA, 1 mM Spermidine) was added then incubated at 37°C for 15 min. Upon completion, reactions were pooled, pelleted, washed, and resuspended in N7 ligation master mix (1X T7 ligase buffer, N7_splint, T7 DNA ligase). Nuclei in ligation master mix was aliquoted across four 96-well LoBind plates, N7 oligos (384 distinct barcodes) added, and incubated at 25°C for 1 hour. Reactions were stopped and incubated at 37°C for 15 min.

Pooled nuclei were pelleted, resuspended in Qiagen EB buffer, and 1,000-4,000 nuclei were distributed across four 96-well plates. Cross-links were reversed by adding EB buffer, Proteinase K, and 1% SDS, then incubated at 65°C for 16 hours. Test PCR with SYBR green determined the optimal cycle number^1^, after which full amplification was performed using NEBNext HF 2x PCR Mastermix, BSA, and indexed P5/P7 oligos. Products were pooled, purified using Zymo Clean & Concentrate-5 and 1X AMPure beads to get rid of adapter dimers, and quantified on an Agilent 4200 Tapestation. Libraries were equimolar pooled at 2 nM and sequenced on an Illumina NextSeq 550 first and then sequenced deeply on a NovaSeq 6000 with custom primers and sequencing recipe (read 1: 51 cycles, read 2: 51 cycles, index 1: 10+15 dark+10 cycles, index 2: 10+15 dark+10 cycles). All splint and barcode sequences are provided in **table S7** of ref ^53^. The PCR duplication rate of individual samples ranges from 19% to 28% (**Supplementary Table 2**).

### Processing of sci-ATAC-seq3 sequencing reads

Sequencing data was processed using the pipeline that we developed for sci-ATAC-seq3^53^ (https://github.com/shendurelab/sciatac_pipeline). Briefly, BCL files were converted to FASTQ format using Illumina’s bcl2fastq/v2.20. Each read was assigned a cell barcode composed of four components: the P5 end of the molecule included a row address for tagmentation and one for PCR; the P7 end of the molecule included a column address for tagmentation and one for PCR. To correct for the errors in these barcodes, we broke them into their 4 constituent parts and corrected them to the closest barcode within an edit distance of 2. If any of the four barcodes could not be corrected to a known barcode, the corresponding read pair was discarded. Reads with corrected barcodes contained in the read name were written out to a separate R1/R2 file for each of the samples in the dataset. Low-quality bases or adapter sequences were trimmed with Trimmomatic/v0.36^89^, and reads were mapped to the mm10 reference genome using Bowtie2^90^ with “-X 2000 -3 1” as options. Read pairs that did not map uniquely to autosomes or sex chromosomes, or had a mapping quality below 10, were filtered out using Samtools^91^. BAM files were then sorted, merged across sequencing lanes using Sambamba, and indexed for downstream analysis.

PCR duplicates within each cell were identified by determining the unique set of fragment endpoints. Within each sample, the resulting BED file of unique fragment endpoints for each cell was used for peak calling with MACS3^92^. Peak calls from all samples were then merged using bedtools to generate a master peak set (*n* = 376,574). Cell barcodes were distinguished from background using a modified approach from the 10x Genomics scATAC pipeline. A mixture of two negative binomials (noise vs. signal) was fitted to determine unique-read thresholds for calling cells in individual samples^53^, with a global cutoff of 500 unique reads per cell applied (**Supplementary Table 2**). Finally, sparse matrices counting unique reads within the master peak set were generated for each sample.

### Initial cell-level QC and doublet removal

To identify high-quality cells, we applied the following criteria to each sample:

1. Unique reads: >1,000
2. Unique reads in peaks: 200 - 200,000
3. Fraction of reads in peaks: above a sample-specific cutoff (15 - 37%)
4. Fraction of reads in TSSs (+/-1 kb): above a sample-specific cutoff (8 - 13%)
5. Fraction of reads in ENCODE blacklist regions: 0.001 - 0.1
6. Nucleosome signal (calculated using the NucleosomeSignal function in Signac/1.9.0^93^): <10

After filtering, we merged the peak-by-cell sparse matrices from all 36 samples, which included six sci-ATAC-seq3 experiments across five NovaSeq runs, resulting in 4,344,905 cells. Dimensionality reduction was performed using Latent Semantic Indexing (LSI) followed by Principal Component Analysis (PCA). Peaks overlapping ENCODE blacklist regions, on sex chromosomes (to avoid batch effects between sexes), or present in <0.1% of cells were excluded. LSI was applied to the binarized peak-by-cell matrix by first weighting all peaks in each cell by log(total number of peaks accessible in that cell) (log-scaled “term frequency”), then multiplying these values by log(1 + the inverse frequency of each peak across all cells) (“inverse document frequency”). Singular value decomposition of the TF-IDF matrix produced a lower-dimensional PCA representation, retaining only dimensions 2-50, as the first dimension correlated strongly with read depth. The PCA matrix was then L2-normalized to account for differences in unique fragment counts per cell and used for downstream analyses. Most analyses were performed with BPCells/v0.1.0^54^. The L2-normalized PCA matrix was also used to perform Leiden clustering using sc.pp.neighbors(n_neighbors = 50) and sc.tl.leiden (resolution = 1) functions implemented in Scanpy/v1.6.0^94^, resulting in 27 cell clusters.

A limited number of doublet removal tools have been specifically developed for scATAC-seq (see: https://github.com/yuelaiwang/CEMBA_AMULET_Scrublet). To our knowledge, these tools fall into two main categories: 1) read count-based methods, such as AMULET^95^, and 2) algorithms adapted from scRNA-seq doublet detection, such as Scrublet^96^. We applied both AMULET and a modified Scrublet algorithm to identify potential doublets. Among the 27 cell clusters, two clusters (6 and 8) showed a high likelihood of being doublets, as identified by Scrublet or AMULET (**Supplementary Fig. 1a**). Cells with AMULET p-value < 0.05 or belonging to these two clusters were removed from downstream analysis, representing 9.4% of the total dataset.

Of note, we observed a reversal in the relative abundance of subnucleosomal fragments (<200 bp) versus mononucleosomal fragments (∼200 bp) across timepoints and experiments. In early stages (before E16.0) or experiments 3-6, subnucleosomal fragments were more abundant than mononucleosomal fragments. In later stages (after E16.0) or experiments 7-8, this pattern was reversed (**Supplementary Fig. 1b**). This reversal was consistent across cell types, suggesting it may be driven by a technical factor.

### Cell clustering and cell type annotations

After filtering low-quality cells and potential doublets, 3,937,903 cells from 36 embryos were retained, with a median of 3,922 reads and 828 peaks detected per cell. We repeated the filtering as described above, excluding peaks in ENCODE blacklist regions, on sex chromosomes, or present in <0.1% of cells, and applied LSI followed by PCA for dimensionality reduction. The L2-normalized PCA matrix (dimensions 2-50) was used for 3D Uniform Manifold Approximation and Projection (UMAP) embedding with n_components = 3, metric = “cosine”, n_neighbors = 50, and min_dist = 0.1. Seurat/v5^97^ was then used to compute a neighborhood graph (n_neighbors = 20) and Louvain clustering (resolution = 0.1), resulting in 29 major cell clusters.

Our recently generated scRNA-seq dataset^42^, from the same set of mice and spanning the same stages of mouse embryogenesis with comparable temporal resolution (∼6 hours), technology (single-cell combinatorial indexing), and scale (11M nuclei RNA-seq vs. 4M nuclei ATAC-seq), is well annotated with >200 cell types, making it a valuable resource for multi-omics integration. We first matched timepoints between the two datasets and randomly selected 15,000 cells per timepoint from each dataset to reduce computational load. Dataset integration was performed using scGLUE/v0.3.2^52^ with default parameters, resulting in a 50-dimensional co-embedding space. Cell type labels were transferred from scRNA-seq to scATAC-seq using a kNN approach, with each scATAC-seq cell assigned the most frequent label among its 10 nearest scRNA-seq neighbors in the co-embedding space. Based on these labels, clusters in scATAC-seq were manually annotated and similar adjacent clusters were manually merged, resulting in 13 Level-1 cell lineages. In addition, we repeated dimensionality reduction using a new feature set of 545,114 genomic windows (5 kb) across the mouse genome. Cell clustering and annotation were robust to the choice of features, whether using merged peaks from each sample or tiled genomic windows (**Supplementary Fig. 2d**).

Next, for each Level-1 cell lineage, we repeated dimensionality reduction, clustering, and cell type annotation, slightly adjusting UMAP parameters (n_neighbors = 30, min_dist = 0.3), resulting in 36 Level-2 cell classes. The same procedure was then applied to each Level-2 cell class, resulting in 140 Level-3 cell types. The three hierarchical levels of cell type annotations are summarized in **Supplementary Table 3**. Of note, when annotating Level-2 cell class and Level-3 cell types, we re-performed integration between scATAC-seq and scRNA-seq, including only scRNA-seq cells expected to correspond to the same populations. For example, scRNA-seq cells from the major cell clusters of neuroectoderm & glia, intermediate neuronal progenitors, eye & other, CNS neurons, and neural crest (PNS neurons) were integrated with neuroectoderm cells in scATAC-seq.

These cell type labels are temporal and require further investigation. Therefore, we created a gene-by-cell matrix with gene-level accessibility scores, counting reads within gene bodies extended by 2 kb upstream. Using these gene-level accessibility scores, analogous to gene expression values, we manually verified whether cell-type-specific marker genes were specifically accessible in the corresponding cell types, revising a few annotations accordingly. The gene markers used for annotating each cell type are summarized in **Supplementary Table 3**.

### Identification of inter-datasets correlated cell types using non-negative least-squares (NNLS) regression

To identify correlated major cell clusters between scRNA-seq and scATAC-seq datasets, we first calculated aggregate expression values (for scRNA-seq) or gene-level accessibility scores (for scATAC-seq) for each gene in each major cell cluster by summing the log-transformed normalized UMI counts of all cells in that cluster. For scATAC-seq, each Level-1 cell lineage was treated as a major cell cluster. To align with scATAC-seq, we manually grouped scRNA-seq cells into 13 major clusters—for example, consolidating neuroectoderm & glia, CNS neurons, neural crest (PNS neurons), intermediate neuronal progenitors, and eye & other into the neuroectoderm cluster, as shown in **Fig. 2c**.

Next, we applied non-negative least squares (*NNLS*) regression to predict gene expression in a target cell cluster (*T_a_*) in dataset A based on the gene expression of all cell clusters (*M_b_*) in dataset B: *T_a_ = β_0a_ + β_1a_M_b_*, based on the union of the 1,500 most highly expressed genes and 1,500 most highly specific genes in the target cell cluster. We then switched the roles of datasets A and B, *i.e.* predicting the gene expression of target cell cluster (*T_b_*) in dataset B from the gene expression of all cell clusters (*M_a_*) in dataset A: *T_b_ = β_0b_ + β_1b_M_a_*. Finally, for each cell cluster *a* in dataset A and each cell cluster *b* in dataset B, we combined the two correlation coefficients: *β* = 2(*β_ab_* + 0.01)(*β_ba_* + 0.01) to obtain a statistic, where high values reflect reciprocal, specific predictivity. We repeated this NNLS analysis using Level-3 cell types, manually regrouping them into six major categories: erythroid, epithelial cells, endothelium, neuroectoderm, mesoderm, and white blood cells. Before analysis, scRNA-seq cell types corresponding to individual Level-3 scATAC-seq cell types were merged as needed, based on the matched cell types summarized in **Supplementary Table 4**.

### Identification cell-type specific peaks

We reorganized fragments from each of the 140 Level-3 cell types and recalled peaks, merging them into a master list of 793,816 peaks. All subsequent analyses, including identification of cell-type-specific peaks, were based on this expanded peak set. Using the master peaks, we generated a new sparse peak-by-cell matrix by counting unique reads mapping to each peak in every cell, which was then binarized.

Cell-type-specific peaks were identified for each Level-2 cell class and each Level-3 cell type. To optimize computational efficiency and balance cell numbers across types, we randomly downsampled 10,000 cells per Level-2 cell class and 2,000 cells per Level-3 cell type. For each peak-cell type pair, a specificity score was calculated using Jensen-Shannon divergence, as previously described^50,53^.

1. Peaks overlapping ENCODE blacklist regions or exceeding two standard deviations from the mean log-scaled counts per peak were removed.
2. For each cell type, the median number of accessible sites per cell was calculated and log10-transformed. The scaling factor for each type was defined as the ratio of the average transformed value across all types to that type’s value.
3. For each peak, the proportion of accessible cells within each cell type was calculated. These proportions were normalized using the cell-type-specific scaling factors and then re-scaled across cell types within each peak to sum to 1.
4. For each peak and cell type, the re-scaled proportion was compared to a perfect distribution (1 for the target cell type, 0 for others) using Jensen-Shannon divergence. The specificity score was calculated as 1 minus the squared divergence.
5. The Jensen-Shannon specificity scores were squared and multiplied by the normalized proportion of accessible cells across cell types from step 3.

### Overlap of cell-type-specific peaks with bulk tissue peaks in ENCODE mouse embryos

We first merged ATAC-seq peaks identified across timepoints (E11.5-E16.5) for each tissue in ENCODE mouse embryos^65^. Next, we randomly downsampled 10,000 cells per Level-2 cell class from our scATAC-seq dataset and removed peaks overlapping ENCODE blacklist regions. Specificity scores for each peak across level-2 cell classes were calculated using Jensen-Shannon divergence. For each level-2 cell class, the 2,000 most specific peaks were selected, and the number of base pairs overlapping bulk ATAC-seq peaks from ENCODE was counted and normalized by the total peak length of each tissue.

### Evolution-naive modeling of chromatin accessibility across 36 Level-2 cell classes

We applied CREsted (https://github.com/aertslab/CREsted), a multi-output convolutional neural network (CNN) framework recently reported by the Stein Aerts lab^2^, to model Level-2 cell class-specific chromatin accessibility. For each of the 36 Level-2 cell classes, Tn5 cut site counts (approximated by fragment boundaries) were aggregated across all cells and normalized using CPM (counts per million; 10⁶ × raw count / total counts in the cell class). The resulting bedGraph files were converted to bigWig format using the bedGraphToBigWig tool. Fragments from each Level-2 cell class were also used to recall peaks, which were merged into a master peak list. Peaks located within ±2.5 kb of transcription start sites (TSSs) of both protein coding and long non-coding genes were excluded, resulting in a final set of 692,058 distal peaks. A peak by cell-class matrix with normalized counts was then used to identify cell-class-specific peaks by applying a softmax function to each peak. Model training was performed using the union of the top 10,000 peaks per cell class (299,281 peaks in total) together with bigWig signal tracks as input, primarily following default settings. Peaks from chromosomes 8 and 10 (∼10% of all peaks) were used as the validation set, while peaks from chromosomes 9 and 18 (∼10%) were held out as the test set. The remaining peaks (∼80%) were used for training. Input sequences were recentered to 2,114 bp, and the top_k_percent parameter was set to 0.03 to normalize peak height across cell classes. Training used the peak_regression mode with the CosineMSELogLoss loss function, and all other configurations followed default settings (batch_size = 256, max_stochastic_shift = 3, learning_rate = 1e-3). The model was trained for up to 60 epochs with early stopping if no significant improvement was observed. After initial training, the model was fine tuned on a subset of 106,813 peaks comprising the top 3,000 most specific peaks per cell type, with the batch_size adjusted to 64 and the learning_rate set to 1e-4. The final model achieved an average Pearson correlation of 0.73 across the 36 Level-2 cell classes (range 0.63 to 0.82) on the held out test set, comparing predicted and observed accessibility within each cell type.

For genome-wide prediction in mouse, chromosomes and unlocalized sequences shorter than 2,200 bp were excluded. The remaining genome was segmented into 2,114 bp windows using a 100 bp sliding step, and segments containing fewer than 2,000 non-N bases were filtered out. The model trained on 36 Level-2 cell classes was then used for prediction, with predicted values assigned to the central 1,000 bp of each input sequence. Predicted values from overlapping sequences were averaged to generate the final chromatin accessibility profile at 100 bp resolution. A softmax function was applied to each 100 bp window to calculate cell class specificity, and the most specific windows for each cell class were selected using thresholds of Z-transformed predicted score ≥ 3 and Z-transformed specificity ≥ 3. Across the whole genome, 10% of 100 bp windows (2.79M) passed these thresholds for at least one cell class.

### Evolution-aware modeling of distal regulatory grammars for 32 non-promoter clusters

We began with 2.79M 100 bp windows identified from the mouse genome that showed both high predicted accessibility and high predicted specificity (Z ≥ 3 for both metrics) in at least one cell class. As an initial filter, these windows were lifted from the mouse genome to 240 mammalian genomes from Zoonomia^3^ (https://zoonomiaproject.org/). We retained 1.47 million windows (53%) that were syntenically preserved in at least half of these mammals. As a second filter, for each retained window, we extracted 2,114 bp sequences centered on the syntenic position in each non mouse mammalian genome, resulting in a total of 296 million sequences (mean 201 orthologs per window). The CREsted model, trained only on mouse data, was applied to all sequences, and for each window we computed the Pearson correlation between the 36 dimensional predicted accessibility vector for the mouse sequence and those of its orthologs. The median correlation across orthologs for each window also showed a bimodal distribution. Using a cutoff of 0.6, we retained 547,317 windows that exhibited both strong evolutionary retention and coherent predicted regulatory activity across species.

To further organize these 547,317 windows, we constructed a kNN graph (k = 30) based on their 36 dimensional predicted accessibility vectors and applied Leiden clustering (resolution = 1). This analysis yielded 33 clusters, including one large cluster dominated by promoters. The remaining 32 clusters showed a strong one-to-one correspondence with our Level-2 cell classes and were annotated accordingly. A few exceptions were observed for cell classes that were not separable and were therefore merged (primitive and definitive erythroid; oligodendrocyte progenitors and committed oligodendrocyte precursors), or that were not distinguishable from the promoter dominated cluster and were therefore excluded (NMPs and erythroid niche cells).

A new model was trained on 354,450 windows from 32 non-promoter clusters (after filtering ENCODE blacklist regions; hereafter referred to as 32 cell classes). Model training followed a similar procedure as for the evolution-naive model. For each of the 32 non-promoter cell classes, normalized Tn5 cut site counts were extracted for individual candidate 100 bp windows, with regions within ±2.5 kb of TSSs of both protein coding and long non-coding genes masked. For cell classes derived from manually merged Level-2 cell classes (primitive and definitive erythroid; oligodendrocyte progenitors and committed oligodendrocyte precursors), the maximum value between the two merged classes was retained. A window by cell-class matrix with normalized counts was then used to identify class-specific windows by applying a softmax function to each window. Training was performed using the union of the top 10,000 windows per cell class (238,538 windows in total). Windows from chromosomes 8 and 10 were used as the validation set, while windows from chromosomes 9 and 18 were held out as the test set. The remaining windows were used for training. Input sequences were recentered to 2,114 bp, and the top_k_percent parameter was set to 0.03 to normalize window heights across classes. Training used peak_regression mode with the CosineMSELogLoss loss function, and all other settings followed defaults (batch_size = 256, max_stochastic_shift = 3, learning_rate = 1e-3). The model was trained for up to 60 epochs with early stopping if no significant improvement was observed. After initial training, the model was fine tuned on a subset of 95,139 windows comprising the top 3,000 most specific windows per cell class, with batch_size adjusted to 64 and learning_rate set to 1e-4.

We performed genome-wide prediction using the evolution-aware model. For this, as with genome-wide prediction using the evolution-naive model, the mouse genome was segmented into sliding 1,000 bp windows, and predictions averaged to 100 bp resolution. However, to mitigate false positive overpredictions induced by tandem repeats, we implemented a heuristic in the mouse reference genome (mm10) was modified by replacing tandem repeats, identified by Tandem Repeats Finder^98^ (TRF), with nearly random sequences that matched to local dinucleotide frequency (± 2,500 bp from the center position of each repeat). We repeated this procedure 10 times (*i.e.* genome-wide prediction on 10 modified versions of the mm10 mouse reference genome) and took the mean prediction for each window as its final prediction score.

### Core region identification using evolution-aware predictions

After applying genome-wide prediction on the mouse genome using the evolution-aware mode, we ranked all genomic windows for each of the 32 non-promoter cell classes by their predicted accessibility. For each score, we defined *P* as the fraction of genomic windows with lower predicted scores and computed a <u>G</u>enome-wide <u>P</u>hred-like <u>S</u>core (GPS = −10 × log₁₀[1 − P]). For each of the 32 cell classes, we selected 100-bp windows with a GPS score > 24.5 (n = ∼100,000; ∼10 Mb in aggregate) and merged adjacent windows. For each merged region, we extracted a 2,114-bp sequence (the CREsted model input length) centered on the region. We then iteratively replaced increasing amounts of sequence from the left and right ends with random DNA matched to local dinucleotide frequencies (± 2,500 bp from the center), following the pattern [20,0] → [20,20] → [40,20] → [40,40] → …, where each pair indicates the number of base pairs replaced at the left and right ends. At each step, both replaced sequence blocks were regenerated using newly sampled random DNA, and accessibility was re-predicted by the model. When the predicted score first fell below 90% of the original value, trimming from the end whose replacement block had most recently increased was halted, while trimming continued from the opposite end until it also caused the score to drop below the threshold. The core region was defined as the subsequence retained immediately before this final score decrease. Counting across all cell classes, a total of 318,423 core regions were identified, with an average length of 530 bp (± 316 bp, standard deviation). The score for each core region was defined as the predicted value immediately before dropping below 90% of its original level and was subsequently converted to a GPS score using the genome-wide predicted score distribution for the corresponding cell type.

### Predicting cell-class-specific gene expression from enhancer activity

To identify the relationship between cell-class-specific enhancer prediction scores vs. cell-class-specific expression of nearby genes, we first regrouped our mouse developmental scRNA-seq timelapse data^42^ into 32 cell classes, and generated normalized pseudobulk expression profiles for each. For each of ∼22,000 protein-coding genes across 32 cell classes, cell-class-specific expression was defined as the log_2_ fold-change relative to the median expression across all cell classes. Each of the 318,423 cell-class-specific candidate enhancers predicted by the evolution-aware model was linked to all nearby genes within 100 kb, and assigned the corresponding cell-class-specific expression value. Enhancer-gene pairs were then stratified jointly by genomic distance (5-kb bins with 2-kb sliding steps) and GPS scores. Within each distance–score bin, we computed the mean cell-class-specific gene expression across all enhancer-gene pairs, and subtracted a background estimate derived from 100 permutations in which enhancer-cell class assignments were randomly shuffled.

### Evolution-augmented modeling enables cross-species inference

We started from the 354,450 × 100-bp mouse windows used to train the evolution-aware model, and incorporated their orthologs from 240 additional mammals^3^. Inclusion required: (i) syntenic retention (successful liftOver & expansion to 2,114 bp) and (ii) functional coherence (pairwise Pearson’s *r* ≥ 0.6 between 36-value evolution-naive prediction vectors for mouse vs. ortholog). Orthologs were assigned the same chromatin accessibility profiles across 32 cell classes as the corresponding mouse sequence. The evolution-augmented CREsted model was trained by: (i) initial training on the 10,000 most cell class-specific mouse windows and qualifying orthologs; (ii) chromosome-based train/validation/test splits shared between mouse (chromosomes 8 & 10: validation; 9 & 18: test; rest: training.) and orthologs, with each ortholog assigned to the same split as its corresponding mouse window, to prevent leakage; and (iii) zeroing regions within 2.5 kb of annotated mouse TSSs, and their orthologs. We trained models with increasing numbers of species (1, 2, 4, … 128, 241), which expanded the data corpus from 0.3M to 58.5M sequences (195-fold). The 1-species model was trained on mouse only, the 2-species model added human, and subsequent models randomly added additional mammalian genomes. All models were trained for up to 20 epochs with early stopping based on validation performance, except for the 128- and 241-species models, which were manually stopped early at epochs 12 and 6, respectively. We selected the model checkpoint exhibiting the highest performance on the validation set (32-species, 4 epochs) and fine-tuned it on the top 3,000 most cell-class-specific mouse windows and their orthologs. The final model, STEAM v1 (Synteny-aware Transfer learning for Enhancer Activity Modeling), was used to predict enhancer activity genome-wide across 241 mammals.

### TF-MINDI analysis

Human and mouse enhancers from *HumMus-v1* tracks were extended to 2,114 bp, and STEAM-v1 was applied to obtain cell-class predictions. For each class in the STEAM-v1 model, the top 2,000 mouse and human enhancers, respectively, were selected based on proportion using CREsted’s *sort_and_filter_regions_on_specificity* function. Contribution scores for these enhancers were then calculated using the *expected_integrated_grad* method. Seqlets were extracted from the central 500 bp of each enhancers (with an additional 10 bp flank and a threshold of 0.5) using TF-MINDI^71^ and tangermeme^72^. Similarity scores were then computed between the extracted seqlets and the SCENIC+ motif collection^73^. The resulting similarity matrix was clustered by first reducing dimensionality with PCA (50 components), then constructing a nearest-neighbor graph (k = 15), and finally applying Leiden clustering (resolution = 3.0). For each cluster, the most prevalent TF DNA-binding domain family annotation was assigned and used as the downstream annotation.

## SUPPLEMENTARY TABLES

**Supplementary Table 1.** Summary of individual embryo samples.

**Supplementary Table 2.** Quality assessment of individual embryos after processing.

**Supplementary Table 3.** Three levels of iterative cell-type annotation, and the gene markers used for each Level-3 cell type.

**Supplementary Table 4.** Cell-type annotations from scATAC-seq and scRNA-seq show high concordance.

**Supplementary Table 5.** Genes showing significant differences in gene level accessibility scores between erythroid niche cells and primitive erythroid cells (*p* < 0.01).

## SUPPLEMENTARY FILES

**Supplementary File 1.** 2.79 million 100-bp windows (Z ≥ 3 for both predicted accessibility and specificity in at least one cell class, based on evolution-naive model predictions).

**Supplementary File 2.** 318,423 core candidate developmental enhancers were identified in the mm10 genome after prediction with the evolution-aware model and trimming to core enhancers.

**Supplementary File 3.** 341,552 core candidate developmental enhancers were identified in the mm10 genome after prediction with the STEAM-v1 model and trimming to core enhancers.

**Supplementary File 4.** 336,378 core candidate developmental enhancers were identified in the hg38 genome after prediction with the STEAM-v1 model and trimming to core enhancers.

**Supplementary File 5.** 336,745 core candidate developmental enhancers were identified in the hg19 genome after prediction with the STEAM-v1 model and trimming to core enhancers.

## DATA AVAILABILITY

In addition, to make all data, models, and predicted tracks more accessible, we created a website hosting all resources listed below.

*Count matrices*

● [Cell metadata](https://shendure-web.gs.washington.edu/content/members/cxqiu/public/backup/jax_atac/download/data_matrix/df_cell.csv) (3,937,903 cells, CSV format)
● Cell-by-peak matrices using peaks merged at the [embryo](https://shendure-web.gs.washington.edu/content/members/cxqiu/public/backup/jax_atac/download/data_matrix/atac_peaks_by_sample.h5ad), [cell lineage](https://shendure-web.gs.washington.edu/content/members/cxqiu/public/backup/jax_atac/download/data_matrix/atac_peaks_by_L1_cell_lineage.h5ad), [cell class](https://shendure-web.gs.washington.edu/content/members/cxqiu/public/backup/jax_atac/download/data_matrix/atac_peaks_by_L2_cell_class.h5ad), or [cell type](https://shendure-web.gs.washington.edu/content/members/cxqiu/public/backup/jax_atac/download/data_matrix/atac_peaks_by_L3_cell_type.h5ad) level
● Fragment files (GEO: GSE325776) and [scripts](https://github.com/ChengxiangQiu/jax-atac-code/blob/main/help_code/reorganize_fragments.py) for regrouping by embryo, cell lineage, cell class, or cell type

*CREsted models*

● [Evolution-naive model](https://shendure-web.gs.washington.edu/content/members/cxqiu/public/backup/jax_atac/download/CREsted_model/evolution_naive_model.tar.gz) (CREsted/1.4.0, keras/3.5.0)
● [Evolution-aware model](https://shendure-web.gs.washington.edu/content/members/cxqiu/public/backup/jax_atac/download/CREsted_model/evolution_aware_model.tar.gz) (CREsted/1.4.0, keras/3.5.0)
● [STEAM-v1 model](https://shendure-web.gs.washington.edu/content/members/cxqiu/public/backup/jax_atac/download/CREsted_model/STEAM_v1_model.tar.gz) (CREsted/1.4.0, keras/3.13.2)

*Cell-class-specific chromatin accessibility*

- Single-cell ATAC-seq data

- Pseudobulk scATAC-seq signal tracks for each cell class ([Bigwig](https://shendure-web.gs.washington.edu/content/members/cxqiu/public/nobackup/jax_atac_L2_mouse_raw_data/mm10/))
- Evolution-naive (mouse)

- Predicted accessibility at 100-bp resolution across the mm10 genome ([Bigwig](https://shendure-web.gs.washington.edu/content/members/cxqiu/public/nobackup/jax_atac_L2_mouse_prediction/mm10/))
- 2.79 million top cell-class-specific 100-bp windows (**Supplementary File 1**) ([Bed](https://shendure-web.gs.washington.edu/content/members/cxqiu/public/backup/jax_atac/download/Supplementary_File_1_100bp_Windows_Specific_to_36_Level2_Cell_Classes.bed.gz))
- Evolution-aware (mouse)

- Predicted accessibility at 100-bp resolution across the mm10 genome ([Bigwig](https://shendure-web.gs.washington.edu/content/members/cxqiu/public/nobackup/jax_atac_32_clusters_mouse_prediction/mm10/))
- 318,423 core candidate developmental enhancers (**Supplementary File 2**) ([Bed](https://shendure-web.gs.washington.edu/content/members/cxqiu/public/backup/jax_atac/download/Supplementary_File_2_Peaks_Specific_to_32_Clusters.bed.gz))
- STEAM-v1 (mouse)

- Predicted accessibility at 100-bp resolution across the mm10 genome ([Bigwig](https://shendure-web.gs.washington.edu/content/members/cxqiu/public/nobackup/jax_atac_augmented_mouse_prediction/mm10/))
- 341,552 core candidate developmental enhancers (**Supplementary File 3**) ([Bed](https://shendure-web.gs.washington.edu/content/members/cxqiu/public/nobackup/jax_atac_augmented_mouse_prediction/mm10/))
- STEAM-v1 (human)

- Predicted accessibility at 100-bp resolution across the hg38 genome ([Bigwig](https://shendure-web.gs.washington.edu/content/members/cxqiu/public/nobackup/jax_atac_augmented_human_prediction/hg38/))
- 336,378 core candidate developmental enhancers (hg38; **Supplementary File 4**) ([Bed](https://shendure-web.gs.washington.edu/content/members/cxqiu/public/backup/jax_atac/download/Supplementary_File_4_Evolution_Augmented_Model_Predict_On_Human_Genome.bed.gz))
- 336,745 core candidate developmental enhancers (hg19; **Supplementary File 5**) ([Bed](https://shendure-web.gs.washington.edu/content/members/cxqiu/public/backup/jax_atac/download/Supplementary_File_4_Evolution_Augmented_Model_Predict_On_Human_Genome_hg19.bed.gz))
- STEAM-v1 (other 239 mammals in *Zoonomia*)

- Predicted accessibility at 100-bp resolution ([Bigwig](https://shendure-web.gs.washington.edu/content/members/cxqiu/public/nobackup/jax_atac_augmented_241_mammals/))
- Core candidate developmental enhancers ([Bed](https://shendure-web.gs.washington.edu/content/members/cxqiu/public/nobackup/jax_atac_augmented_241_mammals_core_enhancer/))

*Interactive UCSC genome browser tracks*

- Guide to adding the track hub in UCSC Genome Browser ([PDF](https://shendure-web.gs.washington.edu/content/members/cxqiu/public/nobackup/Read_Me.pdf))
- List of available track hubs ([TXT](https://shendure-web.gs.washington.edu/content/members/cxqiu/public/nobackup/jax_hub_link_list.txt))

- Raw (Tn5 cut sites were aggregated across cells and normalized to CPM)
- Evolution-naive (100-bp resolution of mm10)
- Evolution-aware (100-bp resolution of mm10)
- Evoluation-aware (100-bp resolution of mm10, after QBS scaling and trimming to core enhancers)
- STEAM-v1 (100-bp resolution of mm10)
- STEAM-v1 (100-bp resolution of mm10, after QBS scaling and trimming to core enhancers)
- STEAM-v1 (100-bp resolution of hg38)
- STEAM-v1 (100-bp resolution of hg38, after QBS scaling and trimming to core enhancers)
- [STEAM-v1](https://genome.ucsc.edu/cgi-bin/hgTrackUi?hgsid=4021021427_ycyuSO7Aigo6JalKfHaAYVIb4veg&c=chr9&g=hub_7227887_jaxAtacMammals) (100-bp resolution of 241 mammals from Zoonomia, anchored to hg38)

---

**Supplementary Figure 1.**
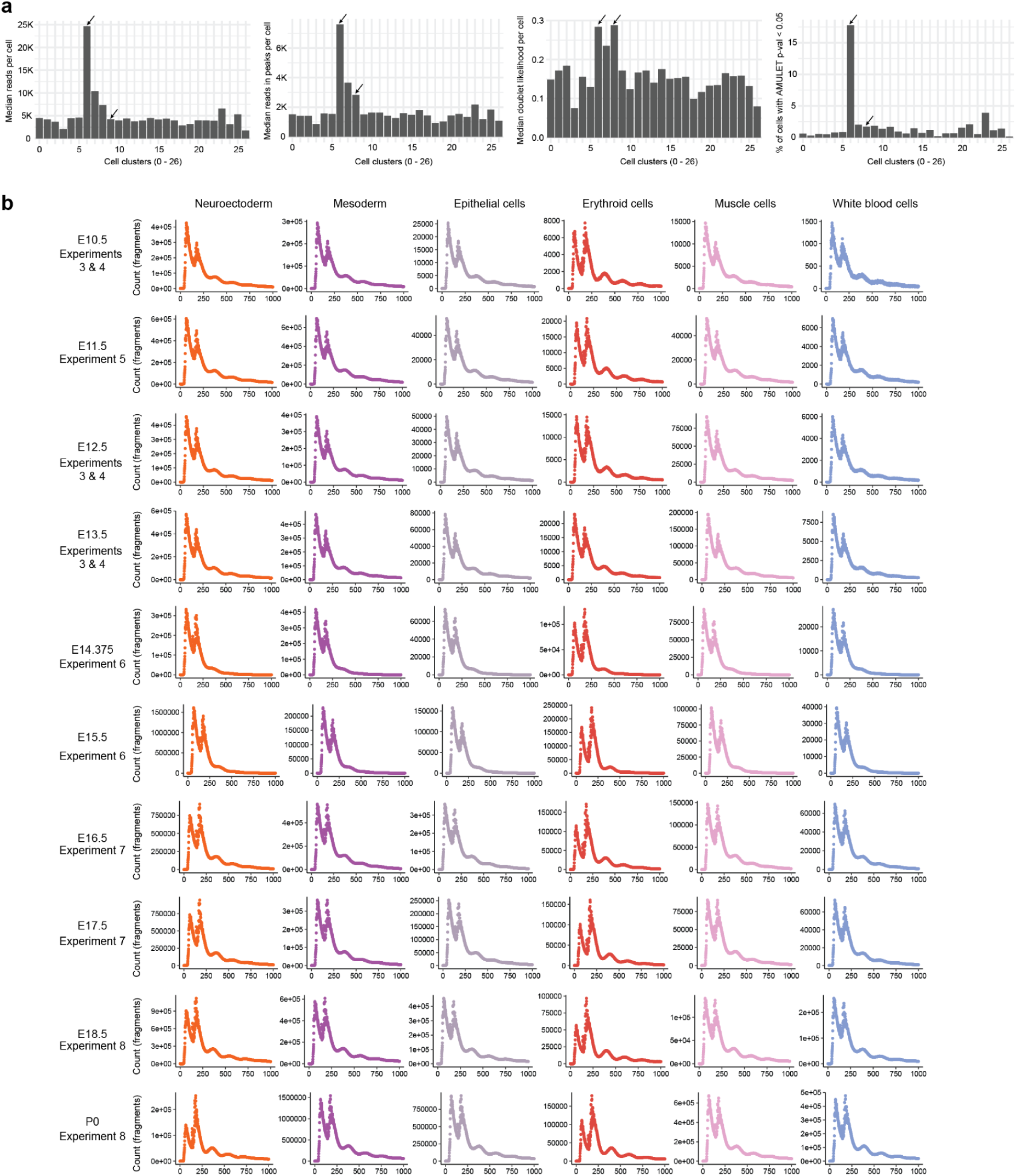
Doublet detection and fragment size distributions. **a,** Prior to doublet removal, we identified 27 cell clusters from 4.3 million cells using LSI-based dimensionality reduction and Leiden clustering with a resolution of one^94^. Doublet detection was then performed with both AMULET^95^ or Scrublet^96^. From left to right: The median number of reads per cell, the median number of reads in peaks per cell, the median doublet likelihood calculated by Scrublet per cell, and the proportion of doublet cells detected by AMULET, are shown plotted for each of the 27 cell clusters. Cells from clusters 6 and 8 (highlighted by arrows in each plot) were identified as doublets and excluded from downstream analyses. **b,** Distribution of fragment sizes varied across timepoints and experiments. Histograms of fragment sizes for cells from 10 representative timepoints (rows; one per day and at least one from each of the 6 experiments) for the six most abundant Level-1 cell lineages (columns) are shown. Fragments greater than 1 kb in length were discarded.

**Supplementary Figure 2.**
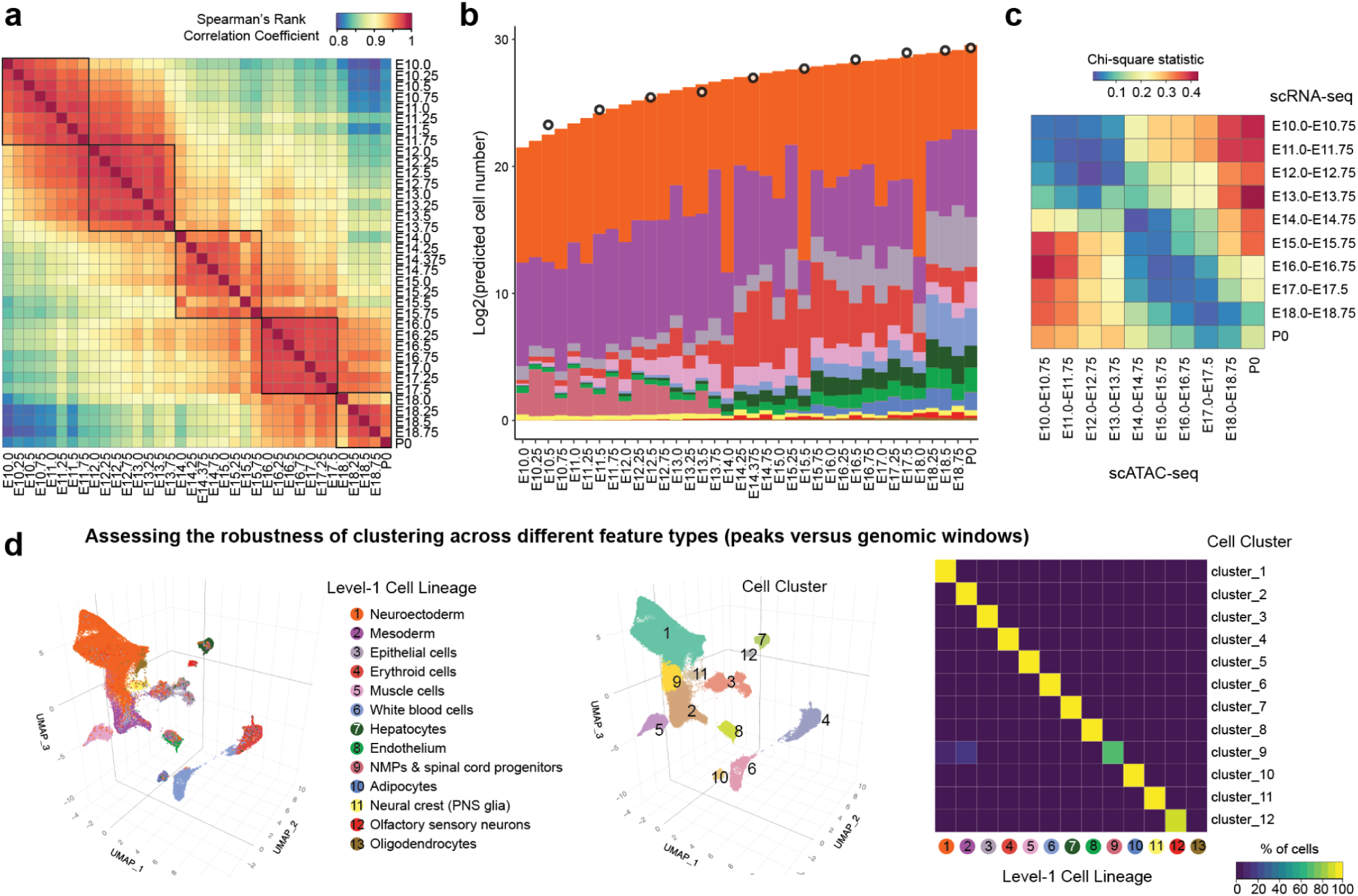
Temporal dynamics and robustness of Level-1 cell lineage annotations. **a,** Spearman correlations were calculated for pairwise combinations of 36 temporal bins, each generated by aggregating scATAC-seq data from a staged embryo after downsampling to a uniform number of cells per sample. Black rectangles are manually added and correspond to two-day intervals. **b,** Composition of embryos from each 6-hour temporal bin by Level-1 cell lineage. The *y*-axis is scaled (log_2_) to the estimated total number of cells in the embryo at each timepoint (see Qiu et al.^42^ for details). Briefly, we isolated and quantified total genomic DNA from whole embryos to estimate total cell number at each of 12 stages spanning E8.5 to P0 (1-day bins, highlighted by black circles), and then applied polynomial regression to predict the log_2_ scaled cell number at each of the 36 timepoints (treating P0 as E19.5 in the regression)^42^. **c,** Compositions across 12 Level-1 cell lineages were compared between scRNA-seq and scATAC-seq using the Chi-squared statistic, where higher values indicate greater differences. For this analysis, cells were grouped in four-day intervals. The NMPs & spinal cord progenitors and neuroectoderm Level-1 scATAC-seq annotations were merged for consistency with scRNA-seq annotations at this level of resolution, while major trajectories from scRNA-seq annotations^42^ were merged as appropriate for consistency with the Level-1 scATAC-seq annotations as well. **d,** To verify the robustness of our clustering strategy to feature choice (peaks vs. genomic windows), we re-performed dimensionality reduction on the read count matrix using 545,114 *x* 5 kb windows of the mouse genome as features. The resulting UMAP is shown twice, with cells colored by Level-1 cell lineage annotations (left) or the major cell clusters identified by the new features (middle). Adjacent major cell clusters were manually merged to achieve a similar resolution. The heatmap (right) shows the percentage of cells from each Level-1 cell lineage annotation (n = 13) within each major cluster (n = 12). Of note, 94% of oligodendrocytes (last column) were included in cluster 1, which predominantly overlaps with neuroectoderm.

**Supplementary Figure 3.**
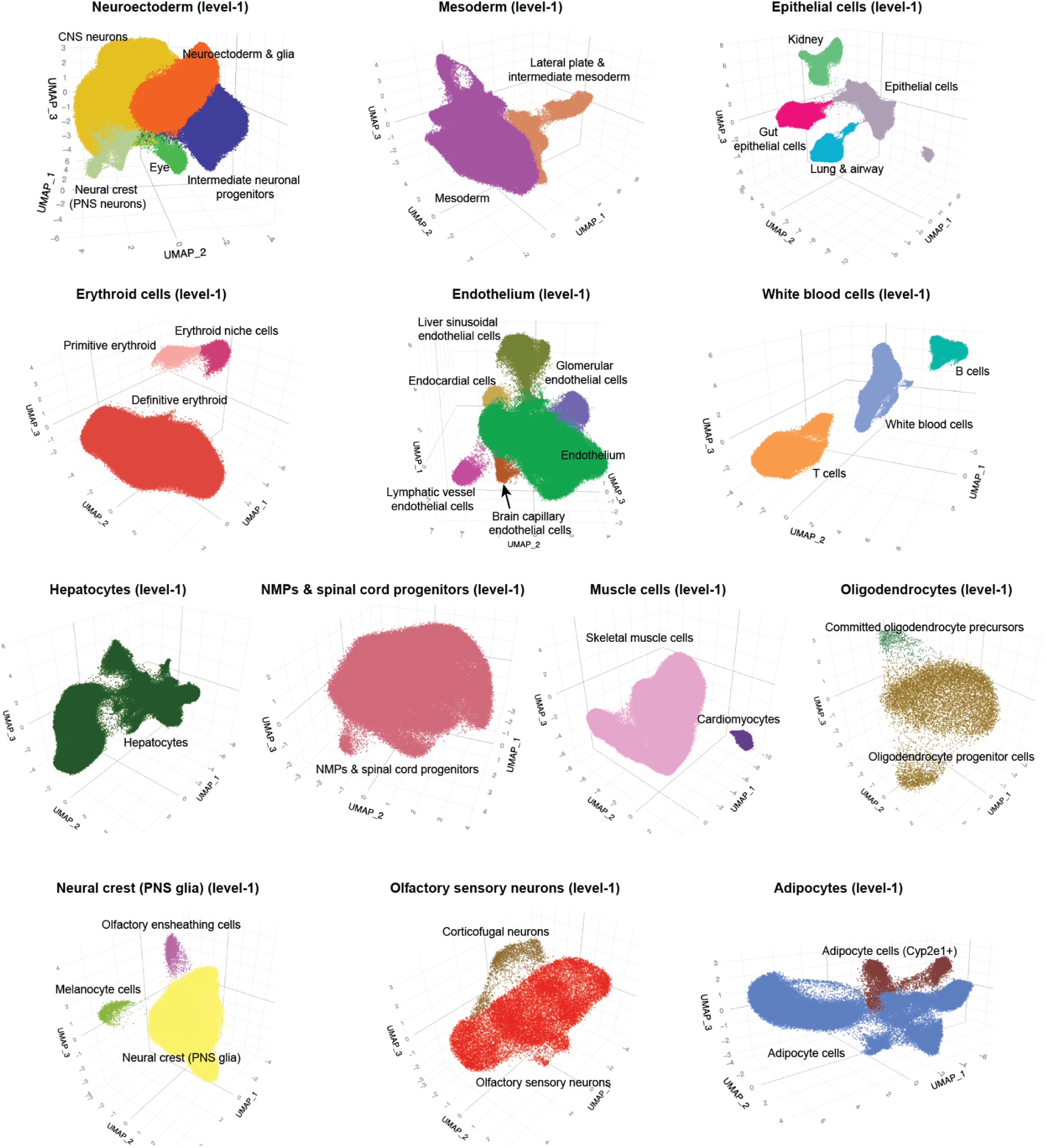
Annotation of 36 Level-2 cell classes. For each of the 13 Level-1 cell lineages, we performed subclustering and annotated the resulting 36 Level-2 cell classes using label transfer from our scRNA-seq atlas^42^ along with at least two literature-supported marker genes per cell type label (**Supplementary Table 3**).

**Supplementary Figure 4.**
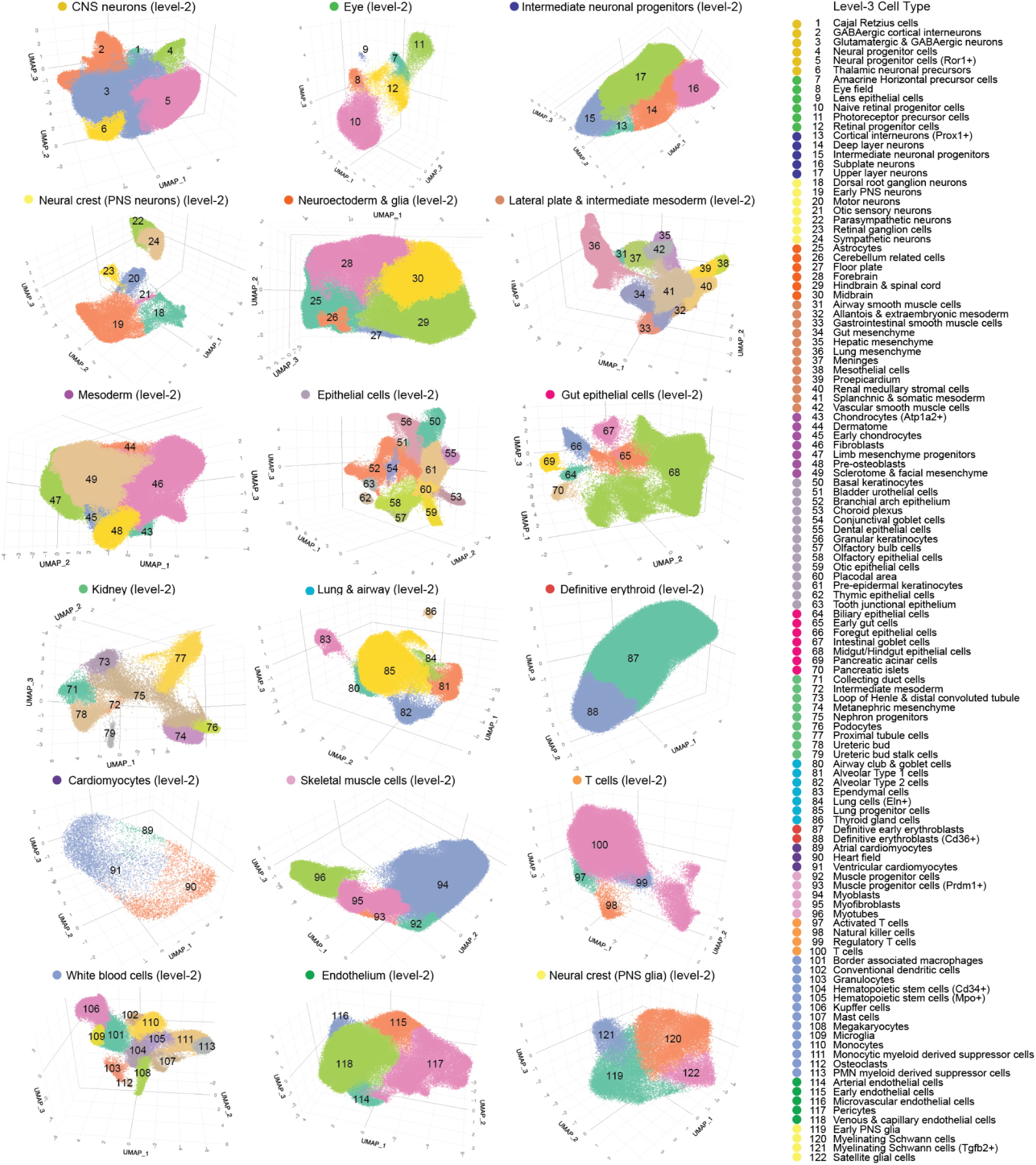
Annotation of 140 Level-3 cell types. For 18 of 36 Level-2 cell classes, we performed subclustering and annotated the resulting 122 Level-3 cell types using label transfer from our scRNA-seq atlas^42^ along with at least two literature-supported marker genes per cell type label (**Supplementary Table 3**). The remaining 18 Level-2 cell classes were not subclustered, and are also represented among Level-3 annotations as cell types.

**Supplementary Figure 5.**
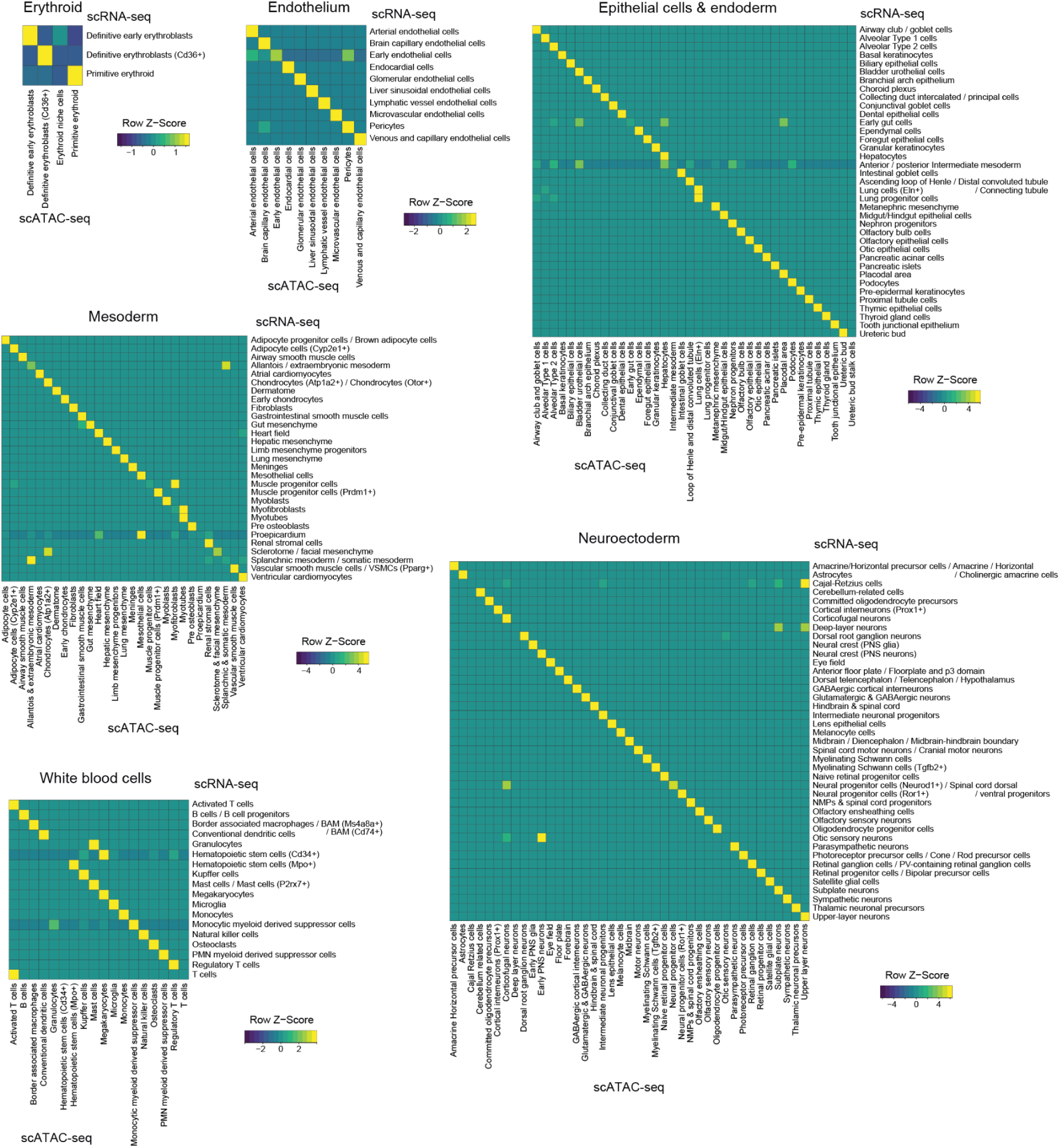
Correlations between scRNA-seq and scATAC-seq cell type annotations based on non-negative least-squares (NNLS) regression. After manually splitting 140 scATAC-seq Level-3 cell type annotations and 170 scRNA-seq individual or grouped cell type annotations^42^ into six major developmental groups, we performed NNLS regression. The resulting heatmaps show the combined regression coefficients (NNLS; row-scaled) for scRNA-seq cell types (rows) vs. 13 scATAC-seq cell types (columns), within each group. Gene-level accessibility scores were used for scATAC-seq (see **Methods**). Prior to this analysis, some subsets of the 56 scRNA-seq cell types were merged prior to performing this analysis, as summarized in **Supplementary Table 4**. CNS: central nervous system. PNS: peripheral nervous system. NMPs: Neuromesodermal progenitors.

**Supplementary Figure 6.**
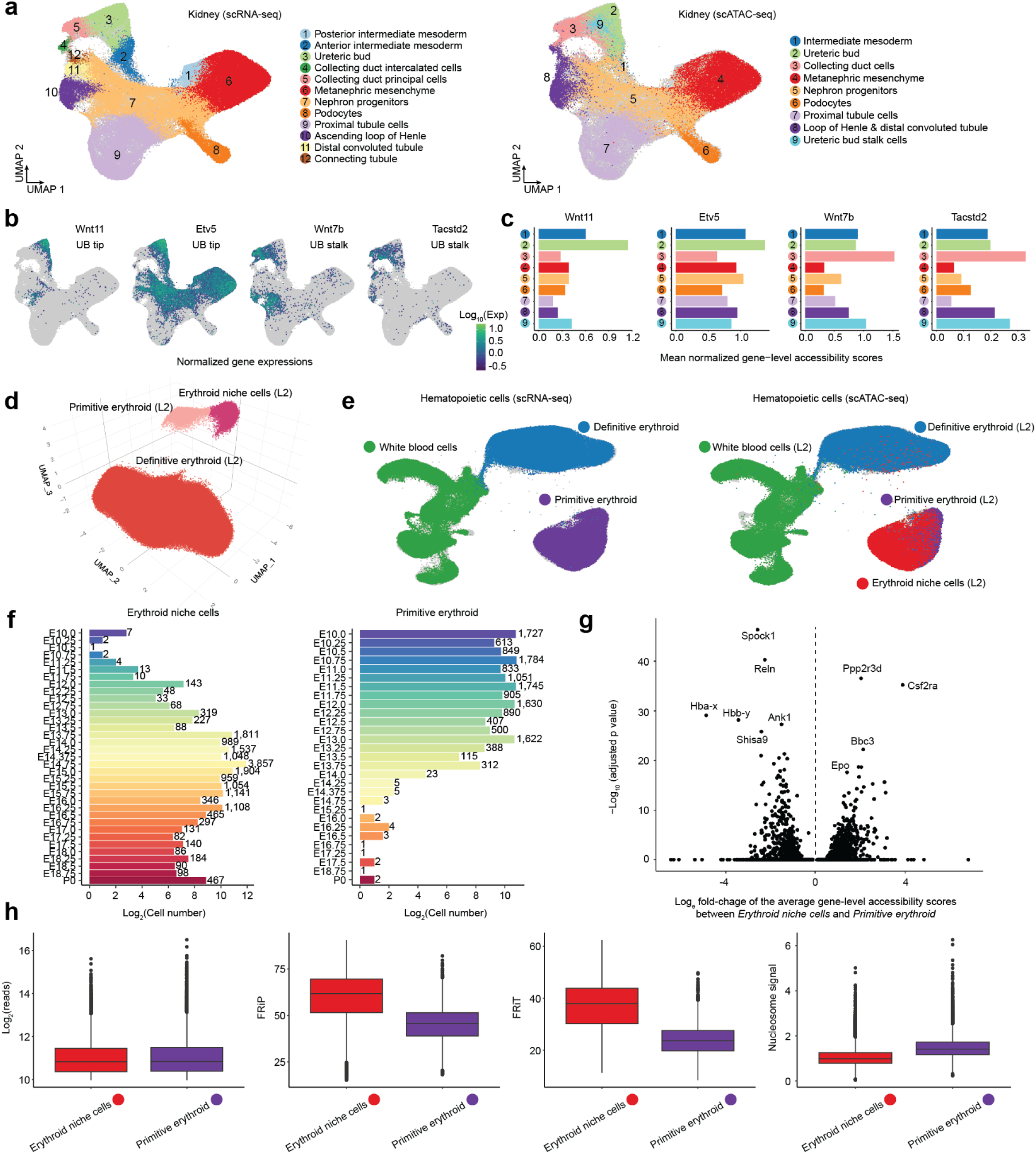
Ureteric bud stalk cells and erythroid progenitors identified uniquely by scATAC-seq. **a,** 2D UMAP visualization of 132,769 co-embedded cells corresponding to renal development, from published scRNA-seq^42^ and scATAC-seq (this study) data, after integration with scGLUE^52^. The same UMAP is shown twice, highlighting cells from either scRNA-seq (left) or scATAC-seq (right) data. **b,** The same UMAP as in panel **a**, colored by expression of marker genes which appear specific to the ureteric bud tip (*Wnt11*+, *Etv5*+) or stalk (*Wnt7b*+, *Tacstd2*+)^99^. **c,** In the scATAC-seq data, gene level accessibility scores were calculated by counting reads overlapping gene bodies plus 200 kb upstream, and those scores were normalized by the total score per cell and multiplied by 10,000. Mean accessibility scores across cells within each renal cell type were plotted for marker genes specific to the ureteric bud tip (*Wnt11*+, *Etv5*+) or stalk (*Wnt7b*+, *Tacstd2*+)^99^. **d,** 3D UMAP visualization of 336,656 scATAC-seq profiles from the erythroid cell lineage, colored by Level-2 cell class. **e,** 2D UMAP visualization of 284,183 co-embedded cells corresponding to hematopoietic development, from published scRNA-seq^42^ and scATAC-seq (this study) data, after integration with scGLUE^52^, colored by Level-2 cell class. Included are all erythroid cell classes and the white blood cell cell class, but not B or T cell classes. The same UMAP is shown twice, highlighting cells from either scRNA-seq (left) or scATAC-seq (right) data. **f,** The number (log_2_ scaled) of scATAC-seq profiles annotated as erythroid niche (left) and primitive erythroid (right) cells, sorted by staging bin. **g,** In the scATAC-seq data, gene level accessibility scores for 21,960 protein-coding genes were compared between the erythroid niche and primitive erythroid annotations after randomly downsampling each group to 1,000 cells, using the *FindMarkers* function implemented in Seurat/v5^97^. The fold-change (natural log scaled) of the average scores between the two groups (*x*-axis) and the negative log_10_ adjusted p-value (*y*-axis) were plotted, with selected significant genes labeled. Full results are provided in **Supplementary Table 5**. **h,** The number of unique reads, the percentage of reads in peaks (FRiP), the percentage of reads falling within TSSs (±1 kb) (FRiT), and the nucleosome signal, were compared between erythroid niche (n = 18,759) and primitive erythroid cells (n = 15,424). The nucleosome signal represents the approximate ratio of mononucleosomal to nucleosome-free fragments^93^. In the boxplots, center lines indicate the medians, and box limits denote the 25th and 75th percentiles.

**Supplementary Figure 7.**
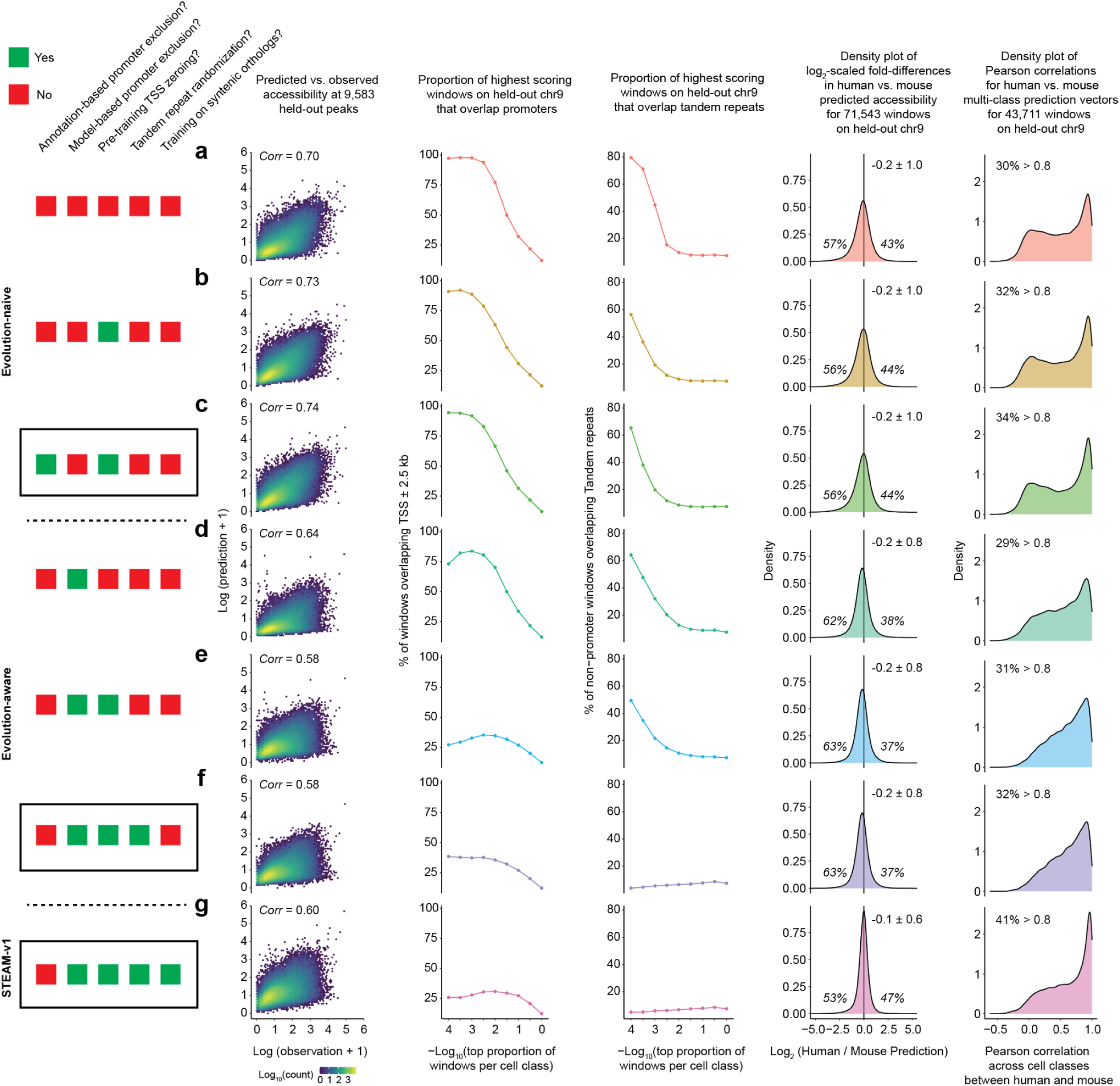
Evaluating successive model iterations against the four modeling objectives. Each row (panels **a**-**g**) corresponds to a model configuration. Each column corresponds to a metric chosen to evaluate one of the four objectives articulated in the main text. Rows are grouped by model iteration. Panels **a**-**c** correspond to several versions of the **evolution-naive** model, trained on 795,998 peaks obtained by merging peaks called on each of the 36 Level-2 cell classes. Panels **d-f** correspond to several versions of the **evolution-aware** model, trained on 354,450 100-bp windows that are syntenically retained, evolutionarily coherent, and excluded from the promoter cluster. Panel **g** corresponds to the **evolution-augmented** model (STEAM-v1), which extends the evolution-aware model by incorporating syntenic orthologs from additional Zoonomia mammals alongside the same 354,450 windows. Within each row group, panels vary along five modifications introduced to address specific failure modes: (i) annotation-based promoter exclusion (TSS ± 2.5 kb); (ii) model-based promoter exclusion (windows assigned to the promoter cluster as shown in Fig. 4d); (iii) manual zeroing of TSS ± 2.5 kb regions prior to training; (iv) tandem repeat replacement during genome-wide prediction (**Supplementary Fig. 10b**); and (v) inclusion of syntenic orthologs from additional species. For each panel, green rectangles indicate a given modification was applied; red indicates it was not. The three models highlighted with black rectangles (panels **c**, **f**, **g**) correspond to the primary evolution-naive, evolution-aware and evolution-augmented models referenced throughout the manuscript. The five columns evaluate the four modeling objectives as follows. **First column (Goal #1, prediction on held-out peaks):** Scatterplots of observed (*x*-axis; normalized Tn5 cut-site counts, log-n scaled) vs. predicted (*y*-axis; log-n scaled) chromatin accessibility across peaks held out during fine-tuning (Fig. 3b), shown for all 36 Level-2 cell classes (evolution-naive) or 32 cell classes (evolution-aware, STEAM-v1), colored by log_10_ scaled density. **Second column (Goal #2, learning distal enhancer rather than promoter grammar):** Predictions across 1.25M × 100 bp windows spanning mouse chr. 9 (Fig. 3g). For the top proportion of windows per cell class (*x*-axis, negative log_10_ scaled), the *y*-axis shows the percentage overlapping with TSS ± 2.5 kb. A model that has learned distal enhancer rather than promoter grammar should show low promoter overlap among its top-scoring windows. **Third column (Goal #3, avoiding overprediction at tandem repeats):** Same prediction outputs as the second column. For the top proportion of windows per cell class, the *y*-axis shows the percentage overlapping with tandem repeats among those not overlapping with TSS ± 2.5 kb. **Fourth and fifth columns (Goal #4, cross-species generalization):** Both evaluate a fixed set of 71,543 100-bp windows on mouse chr. 9 (selected using the evolution-naive model’s prediction and specificity z ≥ 3) with successful liftOver to the human genome. The fourth column shows the density of log₂-scaled fold-differences in predicted chromatin accessibility between human and mouse, with the percentages of windows with log₂ fold differences > 0 and < 0, along with the mean and standard deviation, reported. The fifth column further filters this set to the 43,711 windows overlapping with mouse core enhancers from STEAM-v1 predictions, and plots the density of Pearson correlation across 32 cell classes between mouse and human predictions; the percentage of windows with Pearson correlation > 0.8 is reported. **a,** Evolution-naive CREsted model trained on 306,736 windows (10,000 most specific peaks per class), followed by fine-tuning on 105,671 windows (3,000 most specific peaks per class), to predict chromatin accessibility across 36 Level-2 cell classes. Windows overlapping with annotated promoters (TSS ± 2.5 kb) were not excluded, and promoter accessibility was not zeroed. **b,** Same as panel **a**, but with promoter accessibility zeroed prior to training (297,936 training windows, 107,014 fine-tuning windows). **c,** Same as panel **a**, but with annotated promoter windows (TSS ± 2.5 kb) excluded and promoter accessibility zeroed (299,281 training windows, 106,813 fine-tuning windows). This row corresponds to the primary evolution-naive model referenced throughout the text. **d,** Evolution-aware CREsted model trained on 250,902 windows (10,000 most specific peaks per class) drawn from the 354,450 evolutionarily coherent, non-promoter 100-bp windows, followed by fine-tuning on 95,628 windows (3,000 most specific peaks per class), to predict chromatin accessibility across 32 non-promoter cell classes. Promoter accessibility was not zeroed during training, and no TRF replacement was performed during prediction. However, windows exhibiting promoter grammar were effectively excluded if assigned to the promoter cluster as shown in Fig. 4d. **e,** Same as panel **d**, but with promoter accessibility zeroed prior to training (238,538 training windows, 95,139 fine-tuning windows). **f,** Same as panel **e**, but with TRF replacement performed during prediction. This row corresponds to the primary evolution-aware model referenced throughout the text. **g,** STEAM-v1 model, trained as in panel **f** but incorporating syntenic orthologs across mammals. TRF replacement was performed during prediction.

**Supplementary Figure 8.**
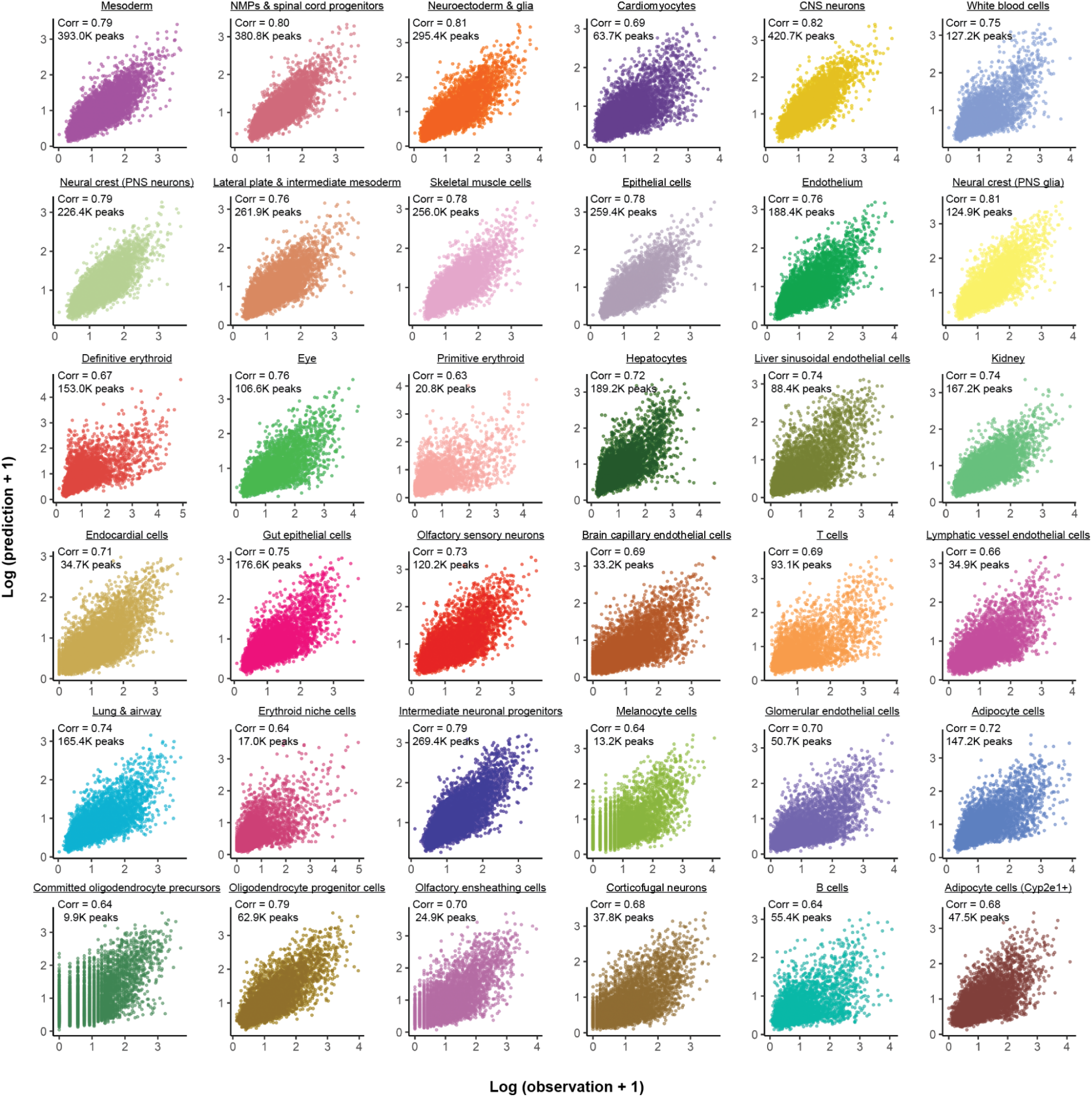
Correlation between observed and predicted chromatin accessibility for each cell class. Scatterplots of observed (normalized Tn5 cut-site counts; log-n scaled; *x*-axis) vs. predicted (evolution-naive CREsted model; log-n scaled; *y*-axis) chromatin accessibility across 9,583 held-out peaks held out during fine-tuning, shown separately for each of the 36 Level-2 cell classes. The Pearson’s *r* and the total number of peaks called in each cell class (before merging to create a master peak list, but after excluding peaks overlapping the ENCODE blacklist or within 2.5 kb of annotated TSSs) are labeled.

**Supplementary Figure 9.**
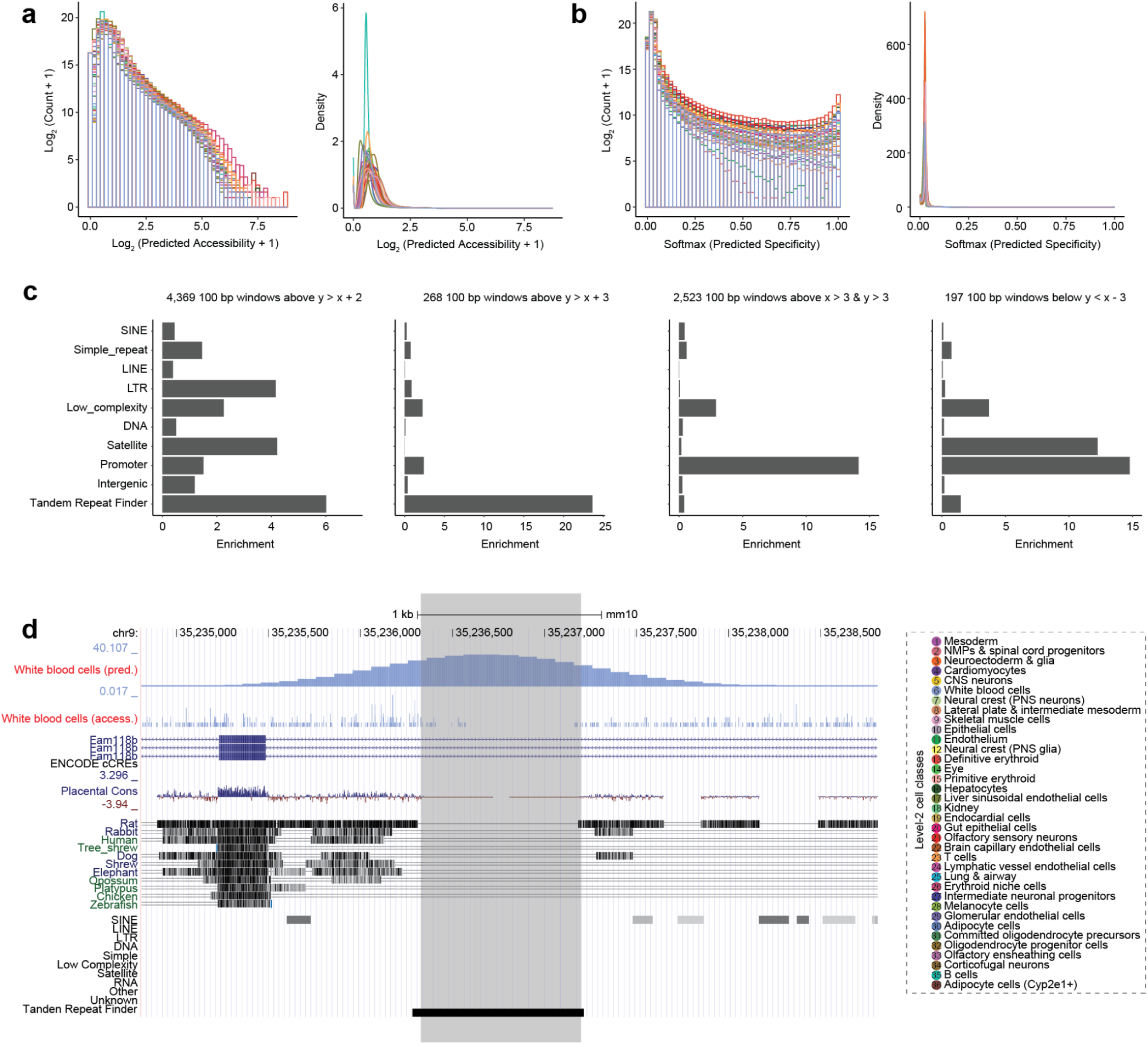
Genome-wide, evolution-naive prediction of chromatin accessibility. **a,** Histogram (log_2_ scaled; left) and density plot (right) of predicted accessibility (log_2_ scaled) for 10% (randomly downsampled) of 28M × 100 bp windows spanning the mouse reference genome. **b,** Histogram (log_2_ scaled; left) and density plot (right) of predicted specificity (log_2_ scaled) for 10% (randomly downsampled) of 28M × 100 bp windows spanning the mouse reference genome. Specificity was calculated by applying a softmax function to the predicted accessibility of each window across cell classes. **c,** Enrichment of various subsets of evolution-naive predictions spanning mouse chromosome 9 for genome annotations, including (from left to right): moderately or grossly overpredicted (*y* > *x* + 2), grossly overpredicted (*y* > *x* + 3), highly accessible and successfully predicted (*x* > 3 and *y* > 3), and grossly underpredicted (*y* < *x* − 3). Enrichment was quantified as the odds ratio: the fraction of each category overlapping a given annotation, divided by the fraction of chromosome 9 covered by that annotation. For this analysis, promoters were defined as regions within 2.5 kb of annotated TSSs. **d,** Genome browser view of observed (normalized Tn5 cut-site counts) and predicted (evolution-naive CREsted model) chromatin accessibility for a representative grossly overpredicted region (highlighted in grey) that overlaps an annotated tandem repeat. Generated using the UCSC Genome Browser (mm10).

**Supplementary Figure 10.**
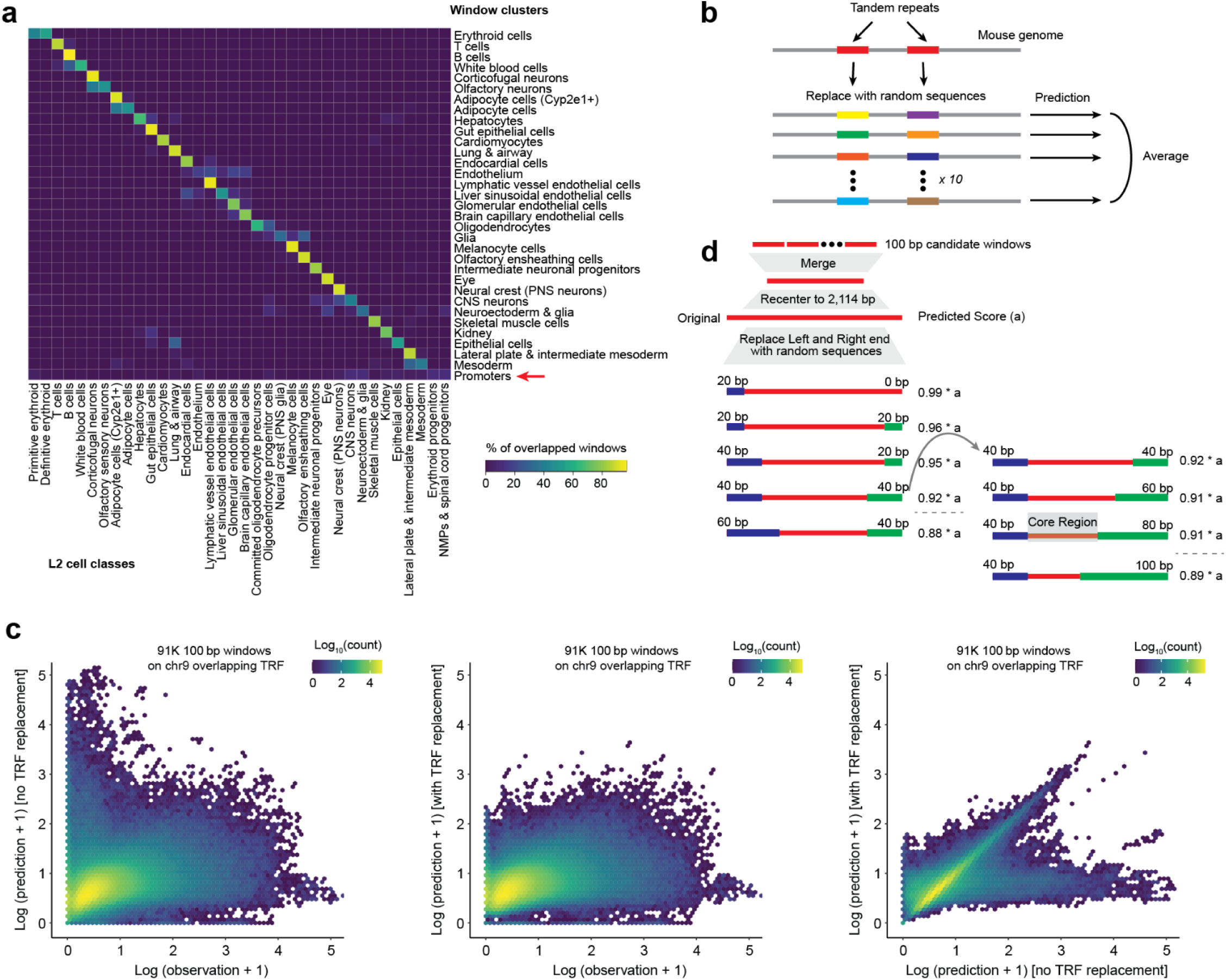
Repeat-aware prediction and core-region trimming refine evolution-aware inference. **a,** The 547,317 × 100-bp windows that exhibited high evolutionary retention and coherence were subjected to Leiden clustering on their 36-dimensional predicted accessibility vectors, which yielded 33 groups, including 32 cell-class-specific clusters and one large cluster dominated by regions overlapping promoters (row with red arrow). For each of the 33 window clusters (rows), we quantified its intersection with 100-bp windows deriving from each of the 36 Level-2 cell classes (columns). The numbers of overlapping windows were first normalized by the total counts within each column and then converted to percentages for each row. Window clusters and Level-2 cell classes exhibited a largely one-to-one mapping, with a few exceptions as discussed in the text. **b,** We performed genome-wide prediction using the evolution-aware model. For this, as with genome-wide prediction using the evolution-naive model, the mouse genome was segmented into sliding 1,000 bp windows, and predictions averaged to 100 bp resolution. However, to mitigate false positive overpredictions induced by tandem repeats, we implemented a heuristic in the mouse reference genome (mm10) was modified by replacing tandem repeats, identified by Tandem Repeats Finder^98^ (TRF), with nearly random sequences that matched to local dinucleotide frequency (± 2,500 bp from the center position of each repeat). We repeated this procedure 10 times (*i.e.* genome-wide prediction on 10 modified versions of the mm10 reference genome) and took the mean for each window as its final prediction score. **c,** Impact of the tandem repeat overprediction mitigation heuristic. Left: Scatterplots of observed (normalized Tn5 cut-site counts; log-n scaled; *x*-axis) vs. predicted (evolution-aware CREsted model <u>without the tandem repeat mitigation heuristic</u>; log-n scaled; *y*-axis) chromatin accessibility across ∼91,000 × 100 bp windows from mouse chr. 9 overlapping TRF-annotated^98^ tandem repeats, for all 32 cell classes. Middle: Same as left panel, but <u>with the tandem repeat mitigation</u> <u>heuristic</u>. Right: Scatterplots comparing predicted chromatin accessibility of these ∼91,000 × 100 bp windows without (*x*-axis) vs. with (*y*-axis) the tandem repeat mitigation heuristic. **d,** Schematic of trimming heuristic used to identify “core regions” driving high scores. For each of the 32 cell classes, we selected 100-bp windows with a GPS score > 24.5 (n = ∼100,000; ∼10 Mb in aggregate) and merged adjacent windows. For each merged region, we extracted a 2,114-bp sequence (the CREsted model input length) centered on the region. We then iteratively replaced increasing amounts of sequence from the left and right ends with random DNA matched to local dinucleotide frequencies (± 2,500 bp from the center), following the pattern [20,0] → [20,20] → [40,20] → [40,40] → …, where each pair indicates the number of base pairs replaced at the left and right ends. At each step, both replaced sequence blocks were regenerated using newly sampled random DNA, and accessibility was re-predicted by the model. When the predicted score first fell below 90% of the original value, trimming from the end whose replacement block had most recently increased was halted, while trimming continued from the opposite end until it also caused the score to drop below the threshold. The core region was defined as the subsequence from immediately before the halt-inducing trim.

**Supplementary Figure 11.**
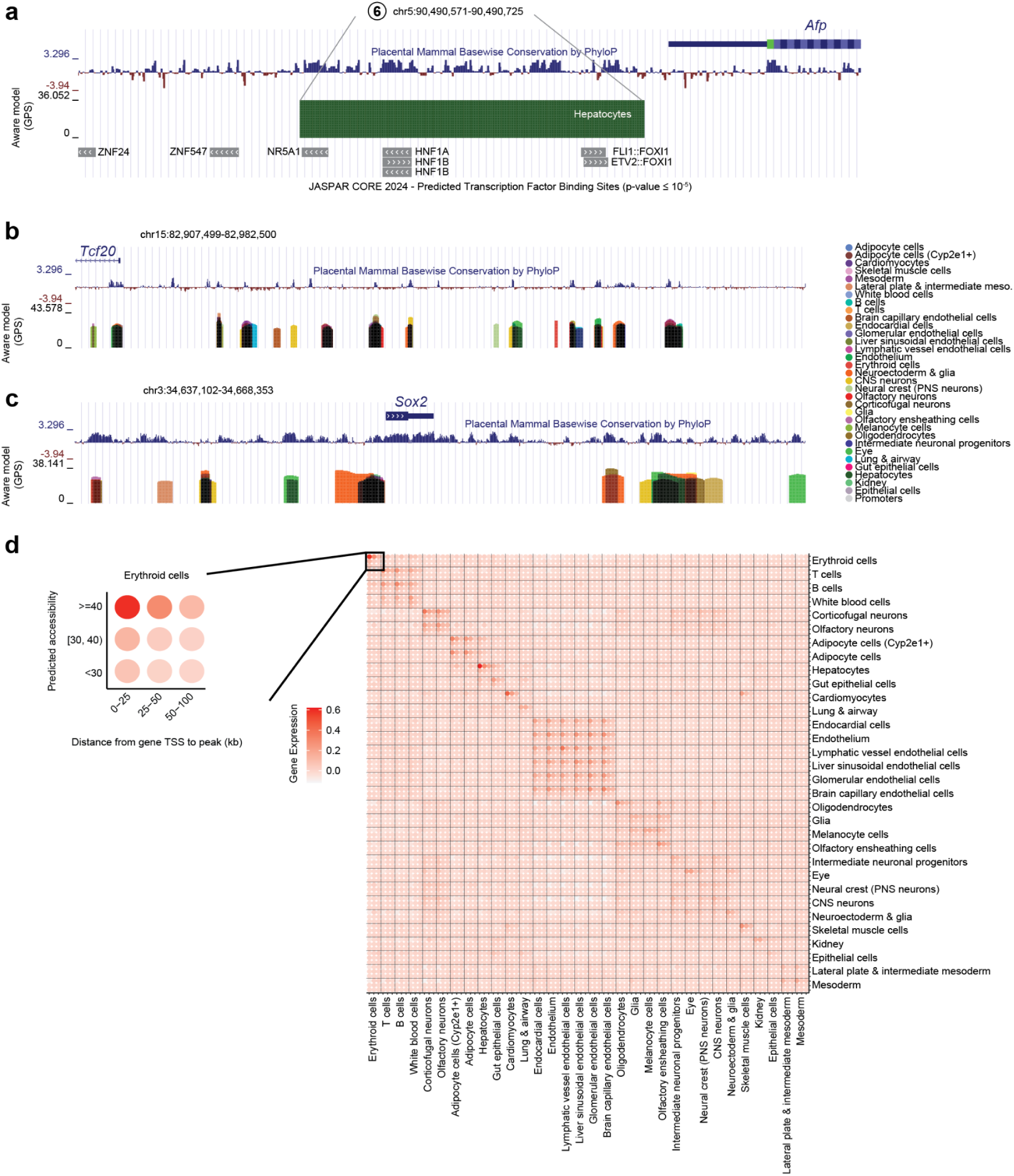
Evolution-aware modeling predicts complex regulatory landscapes near developmental transcription factors. **a,** Detailed genome browser view of candidate hepatocyte enhancer #6 (chr5:90,490,571-90,490,725), plus 100 bp upstream and downstream. Tracks show phyloP conservation, evolution-aware predicted accessibility (GPS-scaled and trimmed to core region) across 32 cell classes (colors), and JASPAR transcription factor binding motif predictions (p ≤ 1×10⁻⁵). The element lies immediately upstream of the *Afp* TSS and is selectively predicted in hepatocytes (green), consistent with either an immediately proximal enhancer or cell-type-specific promoter. **b,** Genome browser view of the region upstream of *Tcf20* (chr15:82,907,499-82,982,500). Tracks show phyloP conservation and evolution-aware predicted accessibility (GPS-scaled and trimmed to core region) across 32 cell classes (colors). Cell class labels are shown on the right. This region contains predicted enhancers with diverse cell-class-specificities. **c,** Same as panel **b** but for the region encompassing *Sox2* (chr3:34,637,102-34,668,353). This region also contains predicted enhancers with diverse cell-class-specificities. **d,** For each pairwise cluster comparison, gene-enhancer pairs were binned into a 3 × 3 grid by genomic distance (*x*-axis) evolution-aware predicted accessibility (GPS-scaled), as highlighted to the left (erythroid cells × erythroid cells). Within each bin of the 3 × 3 grid, the mean cell-type-specific expression was computed and background-corrected using 100 permutations with shuffled peak-cell type assignments.

**Supplementary Figure 12.**
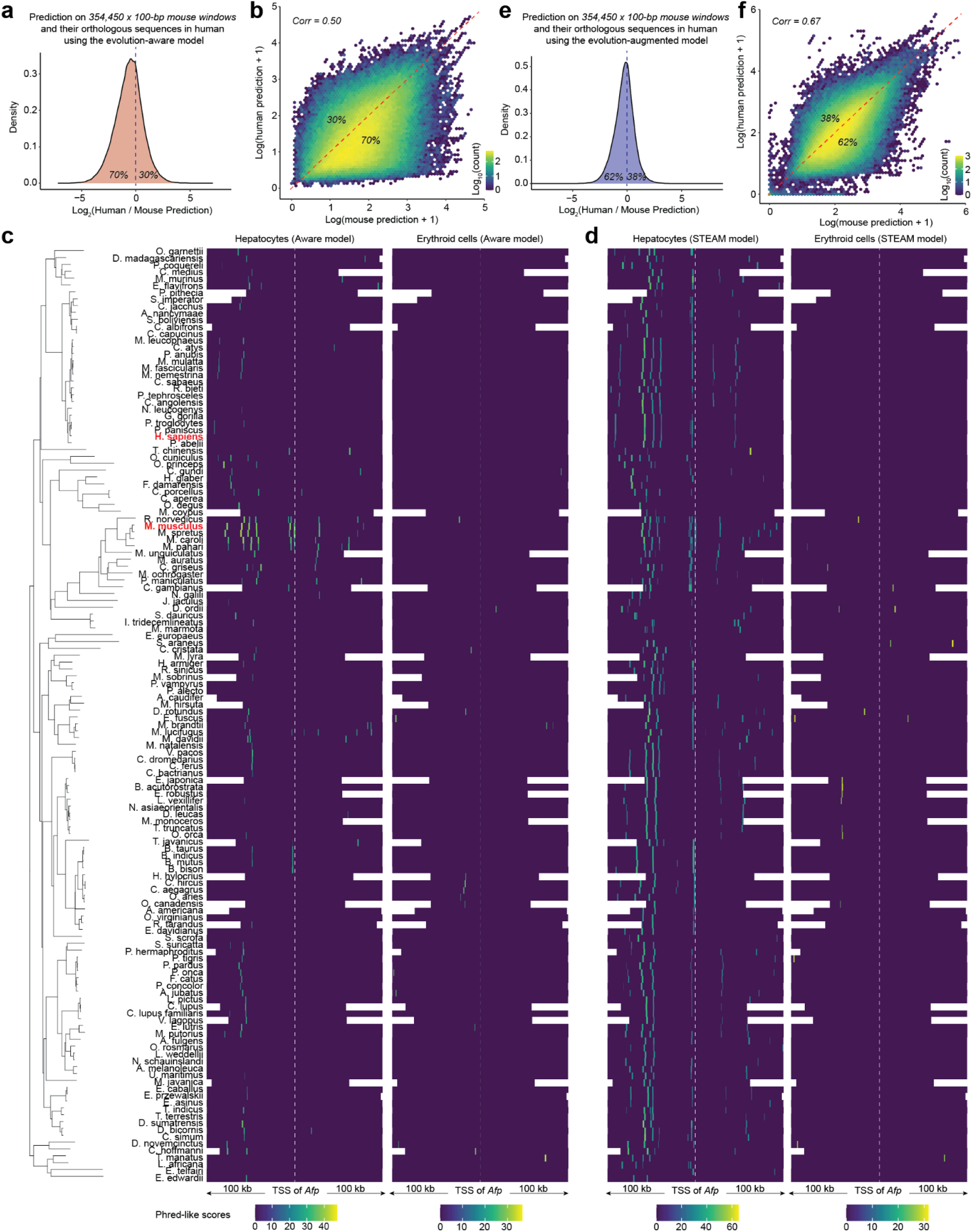
Evolution-augmented modeling improves cross-species inference. **a,** A set of 354,450 × 100-bp mouse windows were lifted over to the human genome. Both the original windows in the mouse genome and the syntenic regions in the human genome were then extended to 2,114 bp and predicted using the evolution-naive model. The density of log_2_-scaled fold-differences between predicted chromatin accessibility for human vs. mouse across 288,982 syntenic pairs is plotted for all 32 cell classes. **b,** Scatterplots show predicted chromatin accessibility in mouse (evolution-aware CREsted model together with tandem repeat mitigation heuristic; log-n scaled; *x*-axis) vs. predicted chromatin accessibility in human (same strategy; *y*-axis) across 288,982 syntenic pairs, for all 32 cell classes, with points colored by log_10_-scaled density. **c,** We examined predicted chromatin accessibility using the evolution-aware model across a 200 kb region centered on the *Afp* TSS (chr5: 90,390,737–90,590,737). The mouse *Afp* TSS was lifted over to other mammalian genomes, and ±100 kb of surrounding sequence was extracted. We retained 136 species with at least 50 kb of contiguous recoverable sequence on each side. Contigs harboring the *Afp* TSS were reoriented to match the top-strand (forward) orientation of the *Afp* transcript in mm10. For each syntenic locus, predictions were performed with tiling, tandem repeat mitigation, and conversion to GPS scores (uniformly calibrated against mouse genome-wide predictions). The left panel shows the phylogenetic tree, restricted to the 136 species included here from the full set of 241 Zoonomia species. The middle and right panels show regions with GPS scores >24.5 from evolution-aware enhancer predictions for hepatocytes (middle) and erythroid cells (right). **d,** Similar to panel **c**, but with predictions made using the STEAM-v1 model instead of the evolution-aware model. **e,** Similar to panel **a**, but with predictions made using the STEAM-v1 model instead of the evolution-aware model. **f,** Similar to panel **b**, but with predictions made using the STEAM-v1 model instead of the evolution-aware model.

**Supplementary Figure 13.**
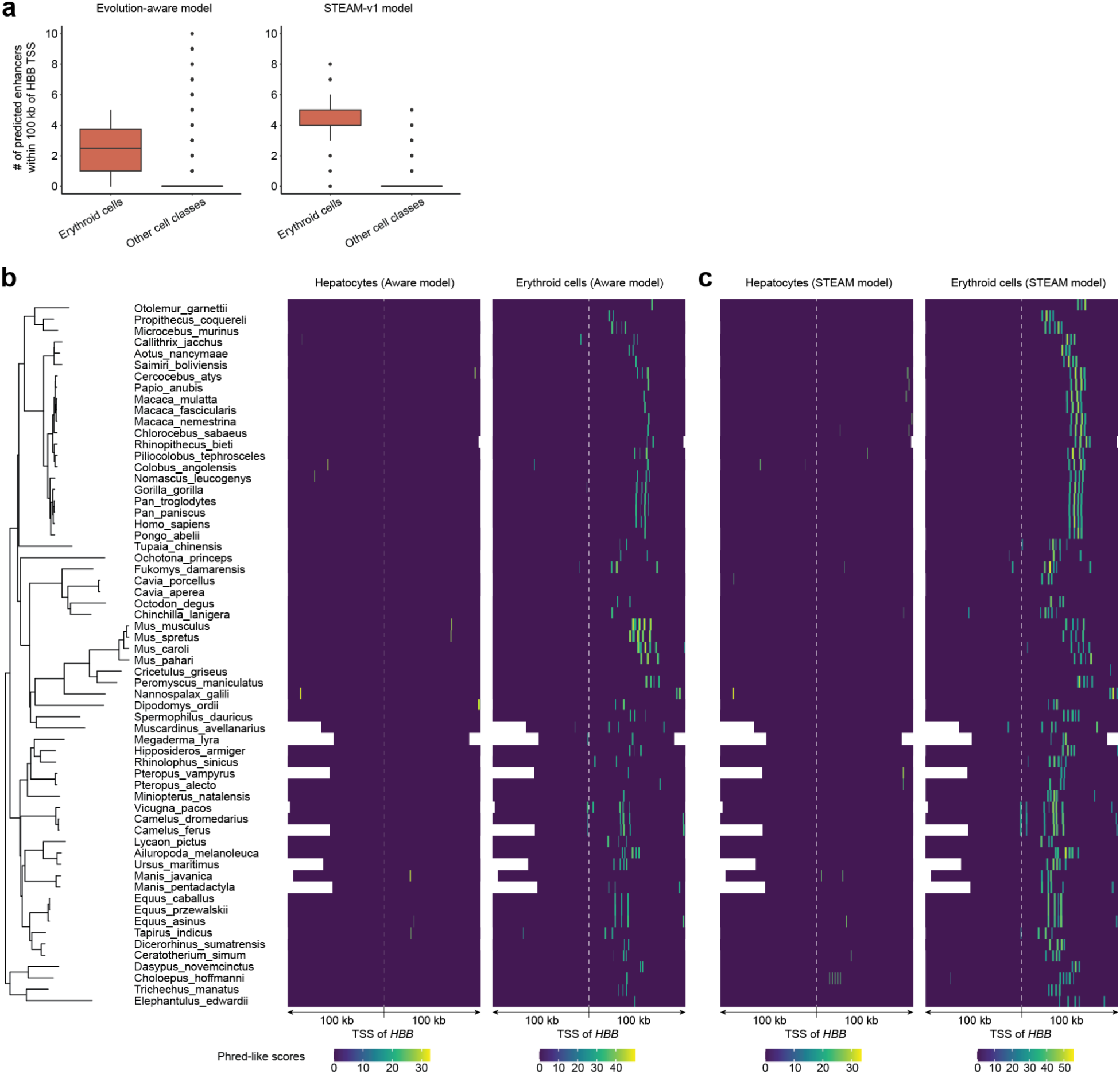
Enhancer prediction at the β-globin locus control region in 62 mammalian genomes. **a,** The human *HBB* TSS was lifted over to 240 Zoonomia^3^ genomes and ±100 kb extracted. 62 species with ≥50 kb of contiguous recoverable sequence on each side were retained. In the human genome, this region includes the classic β-globin locus control region (LCR). The evolution-naive (left) and STEAM-v1 (right) models were applied to each syntenic locus (tiling, tandem repeat mitigation, GPS-based enhancer calling calibrated on mouse genome-wide scale, model-based trimming), and erythroid enhancer counts per species compared against the other 31 cell classes. Boxplot center lines: medians; box limits: 25th–75th percentiles. While the evolution-aware model predicts 2.5 ± 1.4 erythroid enhancers per species (0.6 ± 1.9 for other cell classes), STEAM-v1 predicts 4.2 ± 1.3 (0.2 ± 0.6 for other cell classes). **b,** We examined predicted chromatin accessibility throughout this *HBB* TSS centered region. Contigs harboring the *HBB* TSS were reoriented to match the top-strand (forward) orientation of the *HBB* transcript in hg38. The left panel shows the phylogenetic tree, restricted to the 62 species included here from the full set of 241 Zoonomia species. The middle and right panels show regions with GPS scores >24.5 from evolution-aware enhancer predictions for hepatocytes (left) and erythroid (right) cell classes. **c,** Similar to panel **b**, but with predictions made using the STEAM-v1 model instead of the evolution-aware model.

**Supplementary Figure 14.**
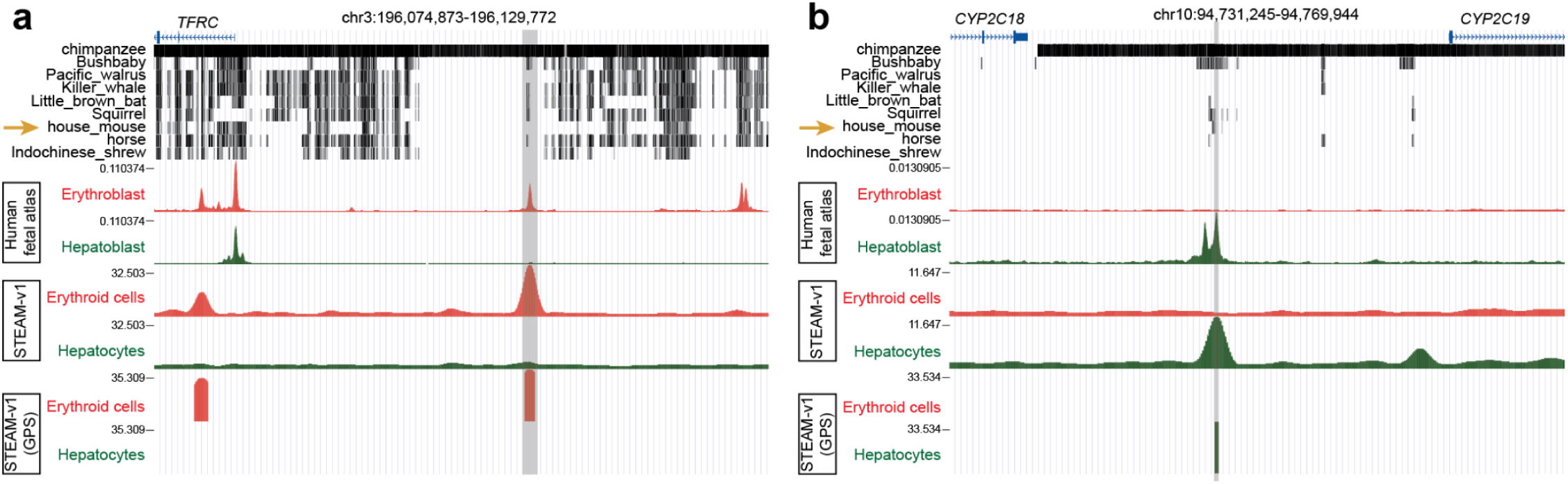
Human-specific enhancers at the TFRC and CYP2C19 loci lack syntenic mouse orthologs. **a,** Genome browser view of a human erythroid-specific enhancer with no syntenic mouse ortholog, located ∼26 kb upstream of the *TFRC* TSS. Tracks show (top to bottom): Zoonomia 241-species alignment^3^, observed liver erythroblast and hepatoblast accessibility from the human fetal atlas^53^, STEAM-v1 predicted erythroid and hepatocyte accessibility, and GPS-scaled core enhancer predictions. Generated using the UCSC Genome Browser (hg38). **b,** Same as panel **a** for a human hepatocyte-specific enhancer with no syntenic mouse ortholog, located ∼15 kb upstream of the *CYP2C19* TSS, in the intergenic region between *CYP2C18* and *CYP2C19*.

**Supplementary Figure 15.**
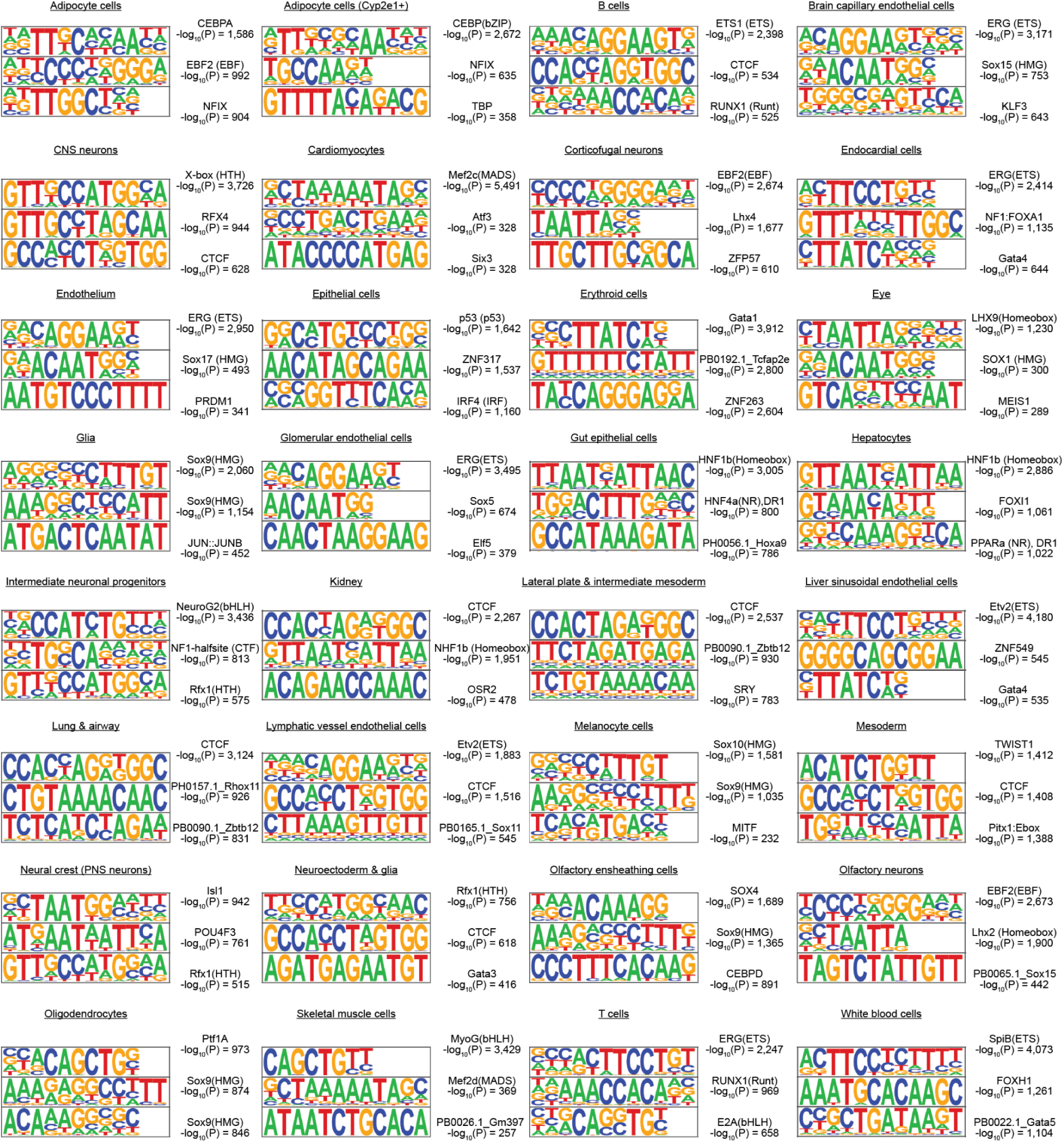
Top enriched TF motifs from STEAM-v1 predicted mouse enhancers across 32 cell classes. The top three *de novo* transcription factor binding motifs (ranked by p-value), along with their p-values and best-matched known transcription factor binding motifs, are shown for each cell class. Motifs were identified from the candidate mouse developmental enhancers predicted by the STEAM-v1 model in each cell class using HOMER^100^ with the findMotifsGenome.pl function.

**Supplementary Figure 16.**
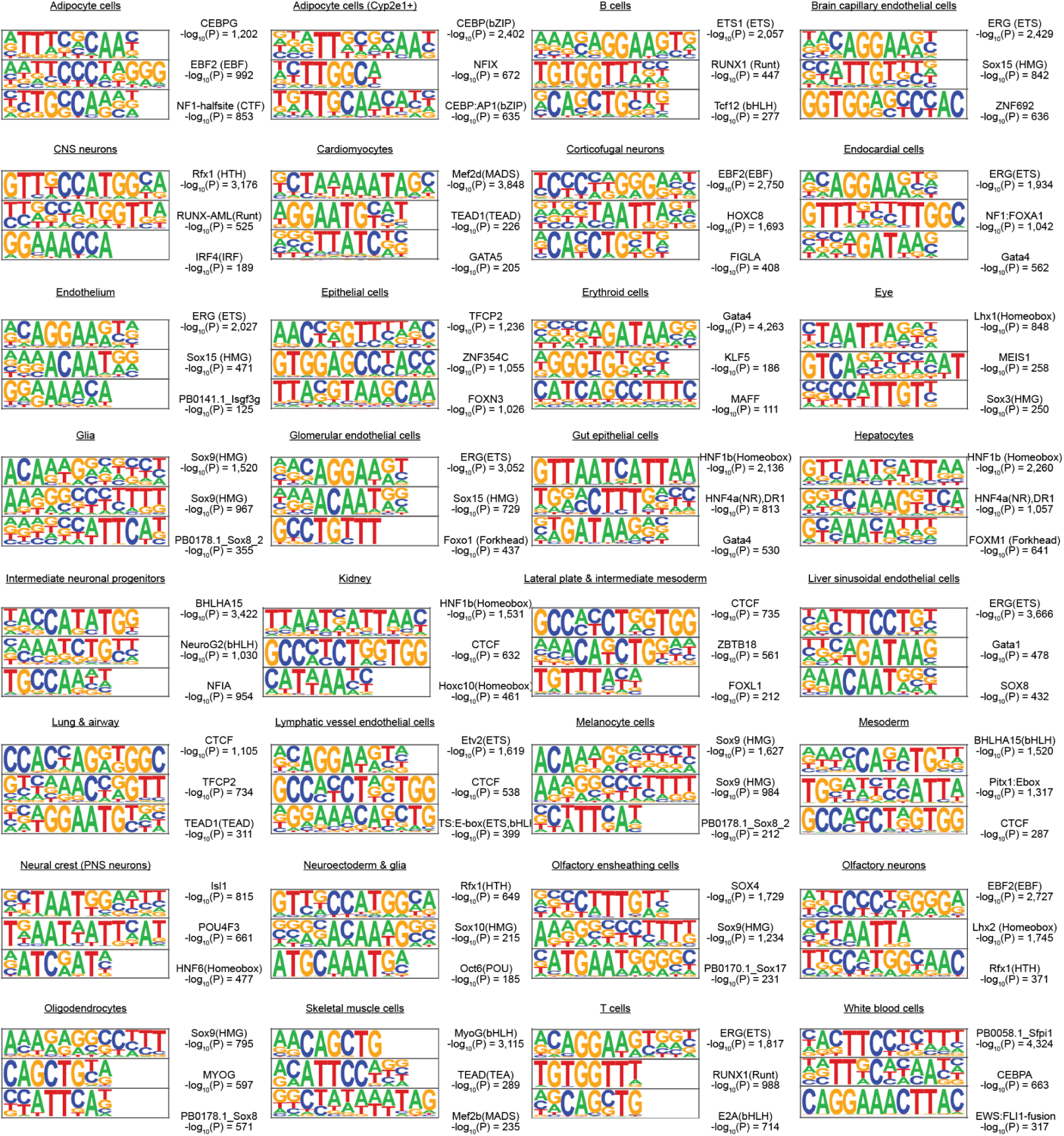
Top enriched TF motifs from STEAM-v1 predicted human enhancers across 32 cell classes. The top three *de novo* transcription factor binding motifs (ranked by p-value), along with their p-values and best-matched known transcription factor binding motifs, are shown for each cell class. Motifs were identified from the candidate human developmental enhancers predicted by the STEAM-v1 model in each cell class using HOMER^100^ with the findMotifsGenome.pl function.

## Notes

https://doi.org/10.62329/hxkk6249

### Summary of Updates

In the revised version, we added a new Supplementary Figure 7 to clarify how the three models were iteratively trained, from naive to aware to STEAM, progressively improving performance toward the goal of genome-wide inference of distal enhancer grammars across all major developmental cell types and all available mammalian genomes.

